# Molecular Organization of the Early Stages of Nucleosome Phase Separation Visualized by Cryo-Electron Tomography

**DOI:** 10.1101/2021.09.01.458650

**Authors:** Meng Zhang, César Díaz-Celis, Bibiana Onoa, Cristhian Cañari-Chumpitaz, Katherinne I. Requejo, Jianfang Liu, Michael Vien, Eva Nogales, Gang Ren, Carlos Bustamante

## Abstract

It has been proposed that the intrinsic property of nucleosome arrays to undergo liquid-liquid phase separation (LLPS) *in vitro* is responsible for chromatin domain organization *in vivo*. However, understanding nucleosomal LLPS has been hindered by the challenge to characterize the structure of resulting heterogeneous condensates. We used cryo-electron tomography and deep learning-based 3D reconstruction/segmentation to determine the molecular organization of condensates at various stages of LLPS. We show that nucleosomal LLPS involves a two-step process: a spinodal decomposition process yielding irregular condensates, followed by their unfavorable conversion into more compact, spherical nuclei that grow into larger spherical aggregates through accretion of spinodal material or by fusion with other spherical condensates. Histone H1 catalyzes more than 10-fold the spinodal-to-spherical conversion. We propose that this transition involves exposure of nucleosome hydrophobic surfaces resulting in modified inter-nucleosome interactions. These results suggest a physical mechanism by which chromatin may transition from interphase to metaphase structures.

## Introduction

The genome of all eukaryotic cells is organized into chromatin, a large complex of DNA and proteins whose structure regulates access to processes such as transcription, replication, and DNA repair. The nucleosome is the basic structural unit of chromatin, and consists of 146 bp of DNA wrapped 1.65 turns around an octameric core of histone proteins (Luger et al., 1997). A diverse set of post-translational modifications of histones and chromatin associated proteins modulate the structural dynamics of chromatin (Bowman and Poirier, 2015), which appears to form two distinct subdomains in the cell nucleus: a lightly packed chromatin or euchromatin state, associated with regions of active transcription, and a condensed heterochromatin state, associated with transcriptionally repressed regions.

Two models have been proposed to explain how chromatin transitions from its transcriptionally active to its inactive form. One model identifies the euchromatin regions with the 10-nm fiber, an extended nucleosome array also referred to as “beads-on-a-string”. According to this model, this structure can fold in a compact, repetitive, and helical structure displaying a 30 nm diameter (Finch and Klug, 1976; Robinson et al., 2006; Song et al., 2014; Widom and Klug, 1985) that corresponds to the heterochromatin domains. In this hierarchical folding model, chromatin condensation occurs through various super-helical structural intermediates formed at different stages of the cell cycle (Belmont and Bruce, 1994). However, this model has been challenged, since *in vivo* studies have failed to reveal regular helical fibers with a 30-nm diameter (Cai et al., 2018; Eltsov et al., 2018; Eltsov et al., 2008; Maeshima et al., 2014a; Maeshima et al., 2014b; Nishino et al., 2012; Ou et al., 2017; Razin and Gavrilov, 2014).

An alternative model for chromatin organization has been proposed recently based on the intrinsic property of chromatin to form liquid-like droplets by liquid-liquid phase separation (LLPS), both *in vitro* and *in vivo* (Gibson et al., 2019; Maeshima et al., 2016; Sanulli et al., 2019; Strom et al., 2017). LLPS is a physical process that underlies the generation of spatially separated membrane-less domains or organelles inside the cell (Banani et al., 2017; Boeynaems et al., 2018; Brangwynne et al., 2009; Erdel and Rippe, 2018; Hubstenberger et al., 2017; Hyman et al., 2014; Sanulli and G, 2020). These condensed phases have been described as LLPS because they appeared under the optical microscope as spherical droplets (Boeynaems et al., 2018) and because chromatin material can diffuse across their phase boundaries (Gibson et al., 2019). In this new model, phase transition of chromatin into droplets drives compartmentalization and organization of long-lasting multiphase systems (Palikyras and Papantonis, 2019; Shakya et al., 2020). Such dynamic chromatin partitioning has been posited to control chromatin accessibility by the cellular machinery (Palikyras and Papantonis, 2019; Shin et al., 2018; Wright et al., 2019). *In vitro*, nucleosome phase transition is stimulated and regulated by nucleosome concentration, nucleosome array length, DNA linker spacing, high-ionic strength, and chromatin-associated proteins such as heterochromatin protein 1 (HP1) and linker histone H1 (Gibson et al., 2019; Maeshima et al., 2016; Sanulli et al., 2019). However, phase transition studies have been circumscribed mainly to the use of fluorescence microscopy, which is limited to study the late-stages of the phase separation process when the condensates have grown to micron size dimensions. A number of questions remain unanswered about the phase separation: first, what is the physical process by which the observed macroscopic domains arise from the microscopic molecular components? Does phase separation and formation of the condensates proceed through a classical mechanism of nucleation and growth, or through the process of spinodal decomposition proposed by Cahn for certain alloys and polymer solutions? (Alberti et al., 2019; Cahn, 1965). Moreover, if condensates are to function as compartments to impede specific enzymatic reactions, chromatin droplets must have a highly compacted structure, while to facilitate reactions, droplets must also be able to adopt a loosely packed structures that enable proteins and other components to diffuse in and out of them (Bancaud et al., 2009; Imai et al., 2017). What is then the arrangement of nucleosomes inside the phase separated condensates? Is the spatial distribution homogeneous or do channels and/or chambers exist that permit molecular diffusion?

To address the above questions requires a technique capable of imaging each nucleosome within the condensates. Yet, given their heterogeneous nature, it is not possible to structurally characterize these condensates using well-established methods such as x-ray crystallography or single particle cryo-EM 3D reconstruction. However, the improvement of electron tomography as a single-molecule 3D imaging method (Ercius et al., 2015; Zhang and Ren, 2012; Zhang et al., 2015), has made it possible to obtain 3D structures at molecular resolution of samples imbedded in vitreous ice (cryo-Electron Tomography or cryo-ET) (Lei et al., 2019a; Lei et al., 2018; Lei et al., 2019b; Yu et al., 2016; Zhang et al., 2016). Here, we have used cryo-ET coupled to deep learning-based 3D reconstruction and segmentation processes to perform a systematic study of the earliest stages of nucleosomal phase separation *in vitro*, using reconstituted tetranucleosome arrays. Tetranucleosome arrays have been identified as the minimal unit capable of generating large-scale chromatin structures (Ding et al., 2021; Schalch et al., 2005; Song et al., 2014), and phase transition experiments with different nucleosome array numbers indicate that tetranucleosomes are at the boundary below which liquid droplets do not form (Gibson et al., 2019).

We find that the initial stages of phase separation, which eventually result in the formation of the large spherical liquid-like droplets observed by fluorescence microscopy, take place through a two-step condensation process. In the first step, uniformly distributed nucleosome arrays undergo global condensation into structures appearing everywhere in the medium without the apparent need of nucleation. This first step resembles a process of spinodal decomposition in which a nucleosome-rich phase composed of irregularly shaped, loosely packed condensates (∼125 nm x 65 nm x 30 nm) emerges throughout the medium at the earliest observation time of 2 min, at room temperature, and in physiological salt. At later times (10 min), under the same conditions, we see the emergence of small, tightly packed spherical nuclei (as small as ∼35 nm in diameter), appearing within the spinodal condensates. These spherical structures are denser than their spinodal counterparts and appear to grow into larger condensates through the accretion of nearby spinodal material or through fusion with other spherical condensates. The spherical shape of these aggregates indicates that their formation is accompanied by an initial unfavorable surface free energy that must be minimized relative to their favorable volume energy. Moreover, the presence of H1 linker histone does not prevent the formation of the initial spinodal phase but catalyzes its transition into spherical condensates more than 10-fold.

## Results

### Observation of the Early Stages of Condensation

To study chromatin LLPS, we used tetranucleosome arrays obtained from the assembly of a DNA fragment that contains four 601-nucleosome positioning DNA sequences separated by 40 bp linkers with recombinant histone octamers (*Xenopus laevis*) (Fig. S1). Nucleosome phase transition time-course experiments at room temperature (20°C) started with the dilution of the tetranucleosomes, stored at low ionic strength (20 mM HEPES-KOH pH 7.5; 1 mM EDTA; 1 mM DTT), to a final concentration of 30 nM using a physiological salt concentration buffer (20 mM HEPES-KOH pH 7.5; 150 mM NaCl; 5 mM MgCl2; 1 mM DTT) (Gibson et al., 2019). To establish the optimal protocol of deposition and time required for the condensate formation, samples incubated for different times were initially subjected to optimized negative staining (OpNS) (Rames et al., 2014) and examined by EM.

Because nucleosome droplets described in fluorescence microscopy experiments exhibit a broad size distribution from ∼0.5 µm to 10 µm in diameter, the samples were imaged at both, low and high magnification (Fig. S2). At the earliest incubation and deposition times (2 min and 20°C), we observed distinct nucleosome condensates adopting irregular shapes (Fig. S2). These condensates were not observed in low salt conditions (1.5 mM Na^+^; Fig. S3A). Interestingly, after 10 min of incubation, small globular condensates (∼35-40 nm in diameter), displaying an apparent higher nucleosomal density, appeared sparsely distributed among the irregularly shaped nucleosome structures (Fig. S2 and S3B). The shapes of these dense condensates were also not uniform. Some of them were spherical, while others were rounded but elongated (Fig. S3B). Significantly, we observed slightly larger structures composed of a dense, rounded condensate, connected with an irregular region (Fig. S3C, white arrow).

After 30 min of incubation, we observed larger spherical condensates (> 100 nm diameter), which tended to increase in number and size with longer incubation times (Fig. S2). Compared to the surrounding irregular condensates, the spherical condensates appeared composed of tightly packed nucleosomes (Fig. S3D), but it is not possible to determine the detailed arrangement of the tetranucleosomes in the spherical condensates using images of negatively stained samples. After 90 min of incubation, we did not observe spherical condensates over 1.8 µm in diameter (Fig. S2 and S4), which suggests that as these dense, spherical condensates grew larger in size, they either became less likely to attach to the surface and/or more likely to detach from the grid surface during the blotting and staining process (Fig. S3E). Epifluorescence microscopy images of samples in the same conditions (Fig. S3F) confirmed that the condensates observed by OpNS correspond to the liquid droplets previously described by fluorescence microscopy (Gibson et al., 2019). Same incubation time series were also repeated at 4°C and 36°C. Statistical measurements of the spherical condensate grid surface density (Fig. S4A) showed that their formation rate is faster at 20°C, followed by 36°C and then 4°C.

### Cryo-Electron Tomography of Nucleosome Condensates

We next used cryo-ET to visualize the nucleosome arrangement in the irregular and the spherical condensates and capture their evolution during the early stages of condensation. To facilitate capture of the condensates on the grid surface, we used a 2D streptavidin crystal as a substrate over the cryo-EM grids (Han et al., 2016) (Fig. 1A) and tetranucleosome arrays harboring a biotin molecule at one DNA end for the crystal surface attachment. The presence of similar spherical condensates on grids prepared with and without the streptavidin crystal, after long incubation times and high salt conditions, indicated that the condensates form in solution, and that the streptavidin crystal simply enabled their attachment to the grid (Fig. S6).

**Figure 1.**
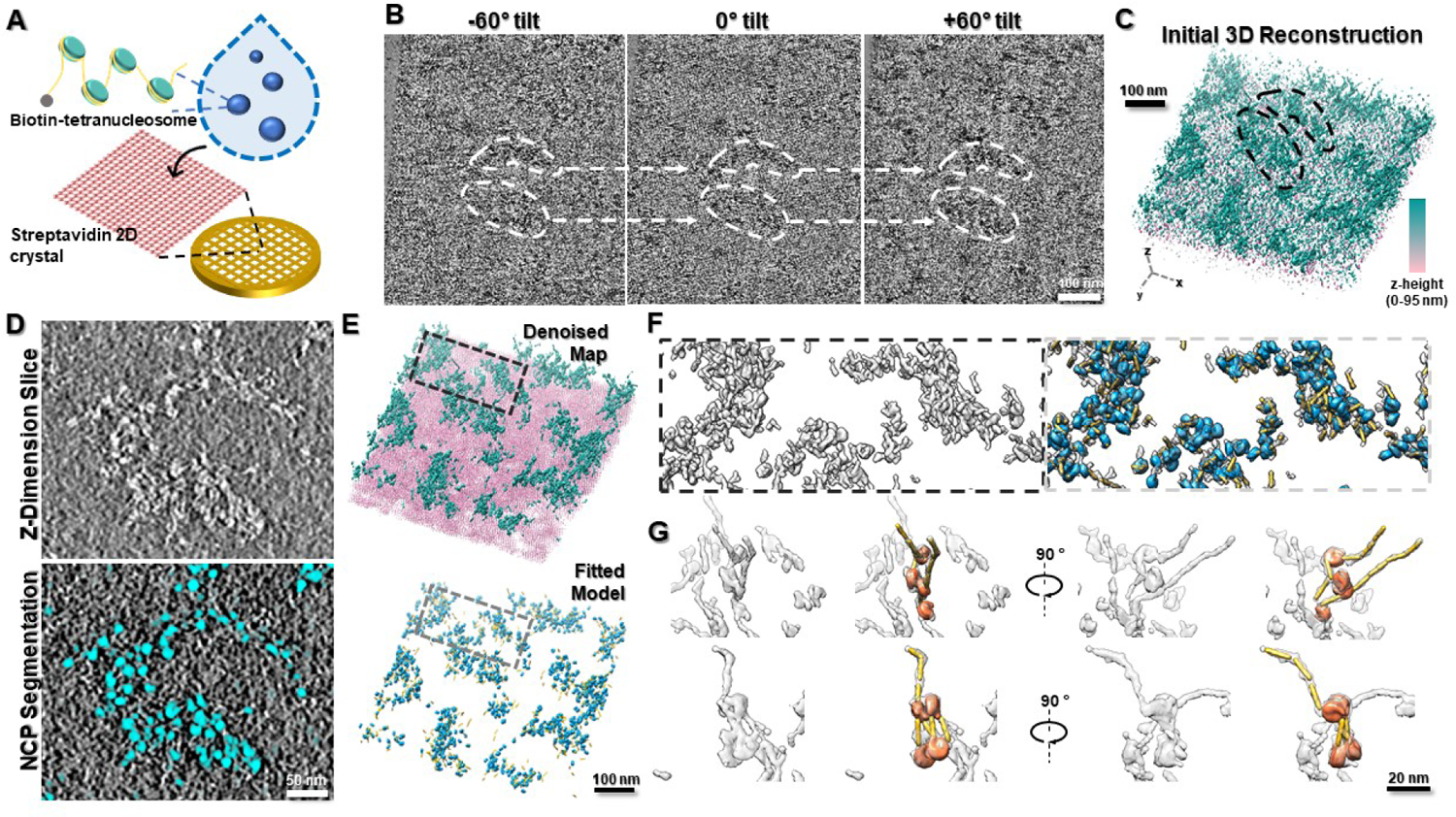
Cryo-ET workflow of data collection, 3D reconstruction, and model docking for samples of the early stages of tetranucleosome phase condensation. (A) Schematic depiction of the incubation of phase-separated condensates of biotinylated tetranucleosomes and their deposition onto EM grids coated with a single layer of a streptavidin crystal. (B) Representative images of tilt series after processing by deep learning denoising. The white dashed contours indicate the tracking of a cluster of visible irregular shape nucleosome condensates along with the tilting. (C) Initial 3D reconstruction of the denoised tilt series after alignment. The reconstructed map is colored by depth from pink to cyan (low to high z-dimensional slice, respectively). (D) Central z-dimensional slice of the initial map (thickness of 1.17 nm, top panel) and the corresponding deep learning-based segmentation of the slice showing predicted NCPs (colored cyan, bottom panel). (E) Missing wedge-corrected final denoised map after reassembling all the tomography pieces shown with the density corresponding to tetranucleosomes depicted in cyan and the density of the SA-crystal is in pink. This final map was then used to generate the fitted model (bottom panel). Blue and yellow correspond to the NCP and a 40 bp DNA model, respectively. (F) Magnified 3D views of the final map (black dashed-line box) and the fitted model (gray dashed-line box). (G) Further zoom of local areas showing clearly distinguishable tetranucleosomes with extended (top panel) or stacked (bottom panel) conformations.

To capture the rapidly formed irregular condensates at the earliest imaging time, we collected tomographic data (Fig. 1B) of the phase transition reaction at 2 min in low-salt (15 mM Na^+^), and at 2 min and 10 min in physiological salt conditions (150 mM Na^+^ and 5 mM Mg^2+^). Given the minimum on-grid incubation time of 2 min for biotin binding to streptavidin (Han et al., 2016), we used the low salt condition to approximate the earliest (0 min) incubation times of the physiological salt concentration. Incubations were initially conducted at 4°C, at which the condensation process occurs more slowly than at 20°C (Fig. S2 and S4A). The cryo-ET of the above three samples under such conditions only showed irregular condensates; no spherical condensates were observed. Spherical condensates were only observed when the nucleosomal arrays were incubated at 20°C for at least 10 min.

In order to determine the spatial organization of tetranucleosomes, we obtained the coordinates of individual nucleosome core particles (NCPs) both free and within each condensate. The cryo-ET workflow we employed is illustrated in figure 1 for a sample incubated for 10 min at 4°C in physiological salt. By using deep learning-based denoising of each tilt image to enhance the low-dose image contrast (Buchholz et al., 2019; Weigert et al., 2018) (Fig. 1B), coupled with a focused refinement strategy of individual-particle electron tomography (IPET) (Zhang and Ren, 2012), we obtained initial 3D density maps in which it was possible to identify nucleosome clusters displaying a low level of condensation (Fig. 1C). The z-dimensional 2D slice of the 3D map (with a thickness of 1.17 nm) showed regions with different degrees of condensation and the irregular arrangement of NCPs within the condensates (Fig. 1D, top panel).

After annotating the initial 3D map using deep learning-based global segmentation (Chen et al., 2019) (Fig. 1D, bottom panel and Fig. S7), we produced a coarse 3D map labelling all NCP positions. To regain NCP molecular details and the connectivity between NCPs in the nucleosome arrays provided by the DNA, the initial and the coarse labelling 3D maps were superimposed, and the later was used as marker to select surrounding connected map density within the former. The resulting maps were then low-pass filtered and served as a mask for local missing wedge correction (Zhai et al., 2020). This procedure yielded the final denoised 3D map depicting NCPs isolated or in condensates, as well as their flanking DNA (Fig. 1E, top panel).

Next, flanking DNA and NCP models (PDB id: 1AOI) were docked into the final 3D map via a weighted iterative search algorithm (see Materials and Methods) (Fig. 1E, bottom panel). We evaluated the result of this fitting by calculating the cross-correlation (cc) score between the local map density and the model. We obtained a cc value of ∼0.8 ± 0.1, corresponding to a high-confidence fitting between the model and the density map when compared to the fitting between the structure model and corresponding modeled density used as a reference (Figure S8 depicts the cc values obtained for all the samples studied here). The final model and 3D map (Fig. 1F, see also Fig. S9-S14) show the distribution of individual NCPs and DNA components. Two representative 3D zoom-in views of individual tetranucleosome arrays show that it is possible to identify their NCPs and flanking DNA arranged in extended and stacked conformations (Fig. 1G top and bottom panel, respectively). Finally, the models obtained were used to determine the spatial coordinates of the individual NCPs.

### The Early Stage of Nucleosomal Condensation Occurs by Spinodal Decomposition

To monitor the process of nucleosome condensation, we calculated the specific space-coordinates of the center of mass of individual NCPs using cryo-ET as described above. We grouped the NCPs’ space coordinates using a density-based clustering non-parametric algorithm (DBSCAN) (Ester et al., 1996). With this algorithm, NCPs surrounded by a minimum of twelve NCPs neighbors (self-included) within 27-29 nm (see Materials and Methods) were classified as nucleosome clusters (Fig. 2A, 2B, and 2C). Each cluster was assigned a different color, with black chosen to specifically depict “free” nucleosomes, i.e., those that could not be associated with any cluster. At 2 min, 4°C, and 15 mM Na^+^, most NCPs appeared dispersed over the imaging area, and only a few of them were grouped into small clusters (Fig. 2A). In contrast, at 2 min of reaction in physiological salt, most NCPs were grouped into larger, irregularly shaped condensates (Fig 2B) that continued to grow after 10 min of reaction (Fig.2C). We propose that the sparse distribution of condensates in 15 mM Na^+^ likely mirrors the earliest stages (0 min) of nucleosome condensation at physiological salt that are otherwise not directly accessible given the earliest possible observation time of 2 min. Thus, the structures observed under this condition will be used as a proxy for the initial state.

**Figure 2.**
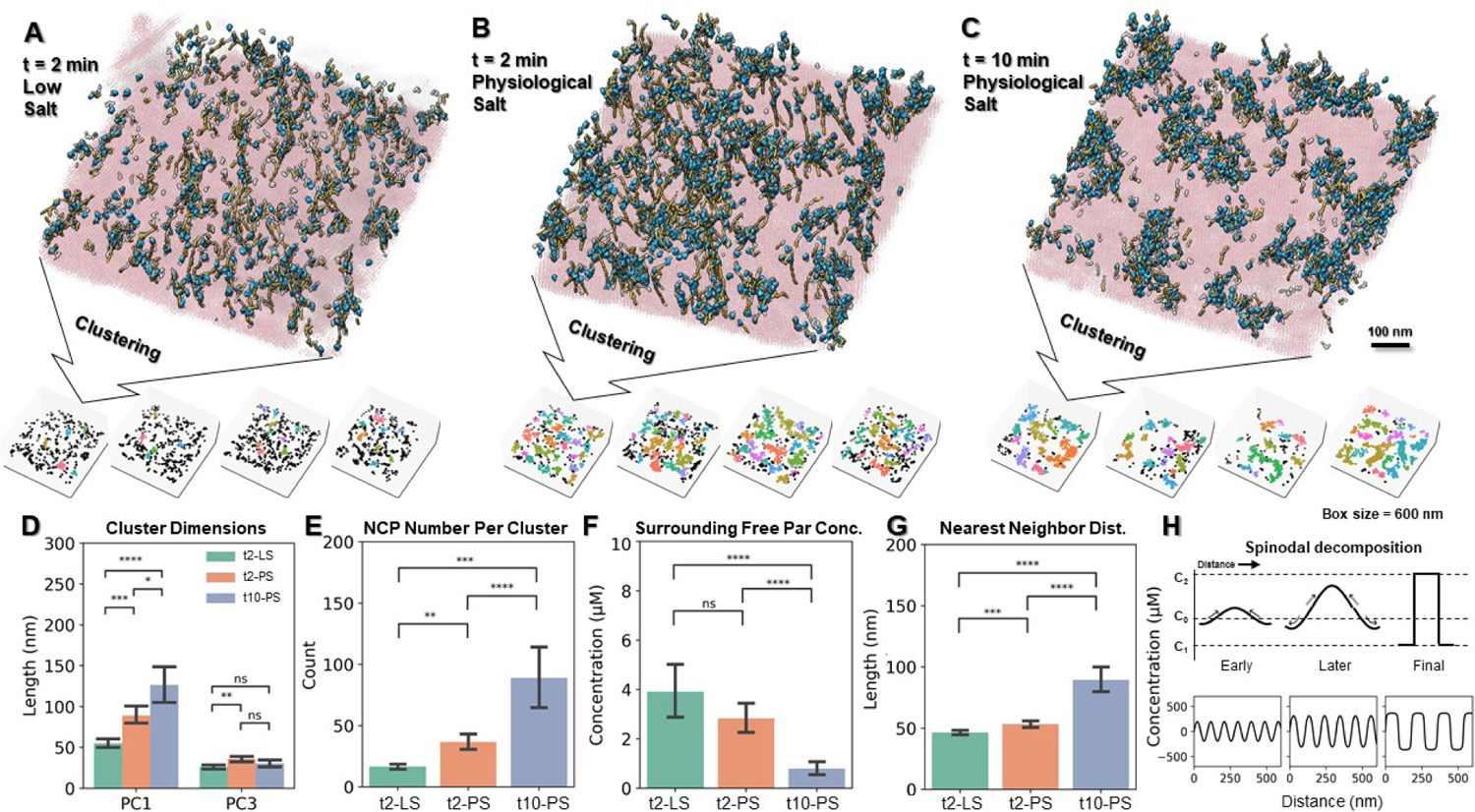
Time evolution of the irregular condensates. Final denoised tomograms with fitted models (tomogram thickness 90-130 nm) displaying irregular condensates obtained at 4°C and 30 nM tetranucleosome concentration. (A; top panel) 2 min incubation at 15 mM Na^+^ (t2-LS); (B; top panel) 2 min at physiological salt (150 mM Na^+^ and 5 mM Mg^2+^; (t2-PS)); and (C; top panel) 10 min at physiological salt (t10-PS). The SA-crystal surface and the nucleosome density maps are depicted in pink and transparent gray, respectively. The fitted NCPs and flanking DNA component are colored cyan and yellow, respectively. The bottom panels show NCP clusters of four representative tomograms for each incubation condition, grouped by the DBSCAM algorithm, with each color representing a unique NCP cluster and black indicating free NCPs that could not be clustered. (D through G) Quantitative analysis of the irregular condensates. (D) Condensates’ dimensions along PC1 and PC3 axes derived from principal component analysis; (E) NCP number per condensate cluster; (F) concentration of free NCPs surrounding a condensate within a 20 nm shell; and (G) nearest neighbor distance between condensates. The data are plotted as mean ± SEM and are shown for the three stages of condensation shown above. All measurements are compared using a two-tailed unpaired t-test, where ∗p < 0.05, ∗∗p < 0.01, ∗∗∗p < 0.001, ∗∗∗∗p < 0.0001; ns, not significant. (H) Schematic representation of the spinodal decomposition process as a function of time obtained from Cahn-Hilliard equation (top panel). Image modified from (Findik, 2013). Evolution of irregular condensates in term of its nearest neighbor distance (wavelength), condensate internal concentration (amplitude). and surrounding free NCP concentration (sharpness) represented by a squdel sine function (bottom panel).

The early condensates exhibited irregular, elongated shapes and were densely distributed all over the surface. We used Principal Component Analysis (PCA) (Hotelling, 1933) to quantitatively characterize these condensates by identifying their long (PC1) and their short (PC3) axes. The average size of the condensate’s long axis after 2 min of incubation at 4°C and 15 mM Na^+^ (the initial reference state) is 55.1 ± 5.2 nm and the average size of the short axis is 26.1 ± 2.4 nm. At 4°C and physiological salt, these values increase to 89.0 ± 10.1 nm and 35.9 ± 3.0 nm after 2 min of incubation, and to 126.9 ± 19.6 nm and 31.6 ± 4.0 nm after 10 min of incubation (Fig. 2D). This analysis indicates that clusters grow asymmetrically along their long axes, and that their short axes stay relatively stable throughout the reaction, with a diameter of ∼30 nm. As a result, their eccentricity (defined here as 1 – short axis/long axis) increases from 0.51 ± 0.05 to 0.56 ± 0.03 to 0.73 ± 0.03 at initial, 2 min, and 10 min of reaction times, respectively (Fig. S15A). The growth of the condensates at these time points was accompanied by an increase in the average number of NCPs per condensate, from 17 ± 3, to 37 ± 6, to 89 ± 26 (Fig. 2E). Next, we quantified the depletion of isolated NCPs surrounding the condensates as the reaction proceeded, by determining the number of isolated NCPs found within a 20 nm shell surrounding the condensate (Fig. 2F). At the initial reference state, the concentration of NCPs within the shell was 3.9 ± 1.0 µM; in physiological salt this value decreased to 2.8 ± 0.5 µM after 2 min and to 0.8 ± 0.3 µM after 10 min of condensation reaction (Fig. 2F). To further validate this analysis, we determined the total number of free NCPs within the imaged area and calculated their concentration in the ice slab. Consistent with the above results, the concentration of free NCPs was found to be 25.0 ± 3.5 µM, 8.9 ± 2.3 µM and 3.9 ± 0.7 µM for the initial reference state, and for the 2 min and 10 min observation times in physiological salt, respectively (Fig. S15C). The total (free + condensed) concentration of NCPs within the ice slab throughout the reaction slightly increased, as it was found to be 31.7 ± 5.4 µM, 41.0 ± 3.7 µM and 48.4 ± 16.8 µM (Fig. S15D) for the three time points analyzed.

We propose that these observations are consistent with a mechanism of spinodal decomposition, which involves negative diffusion of solute material against a concentration gradient, gradually forming irregular condensates with defined borders. Spinodal decomposition does not involve the crossing of an energy barrier, and therefore the formation of condensates is ubiquitous and arises as a result of local, small concentration fluctuations in the medium. The growth of these condensates can be described in terms of a time-dependent periodic distance or wavelength λ(t), which gives the average separation between domains of the condensed phase, ultimately attaining a maximum value λmax (see Supplementary Theory Section, Equation 9) that determines the average distance between condensates (Emo et al., 2014). The spacing among all condensate clusters was estimated by calculating the mean nearest neighbor distance between condensates using a low pass-filtered map (8 nm) (Fig. S16). The average spacing between condensates gradually increased from 46.5 ± 1.8 nm to 53.3 ± 2.8 nm, to 89.5 ± 9.8 nm at 10 min (Fig. 2G). A pairwise correlation test among five of the independent measurements described above, i.e., condensate cluster size, eccentricity, number of NCPs per condensate, concentration of surrounding free NCPs, and nearest neighbor distance, yielded an average value of 0.73 for the corresponding absolute pair-wise r-values (Fig. S17). The strong correlation confirms that the condensates grow by the asymmetric accretion of isolated NCPs, leading to the formation of elongated condensates according to a spinodal decomposition process (Fig. 2H). This type of anisotropic morphology has been described as a characteristic feature of spinodal processes, with the appearance of elongated processes resulting at early times from diffusion effects, and isotropy regained at longer times driven by hydrodynamics effects (Datt et al., 2015).

### Spherical Condensates Arise by Nucleation and Growth in the Later Stages of Phase Separation

The denoised, zero-tilt images of samples prepared in physiological salt that were incubated for 10 min at room temperature revealed distinct, spherically-shaped structures of varying sizes, from ∼40 nm (Fig. 3A; panel I) to ∼400 nm in diameter (Fig. 3A; panel IV) that sparsely appeared among the spinodal condensates. These condensates seemed comparatively denser than the surrounding irregular spinodal condensates and resembled those observed in the OpNS samples (Fig. S2), suggesting that they are an early manifestation of the formation of liquid droplets. Their sparse distribution (Fig. S18A) is consistent with an energetic barrier that must be crossed for these condensates to form.

**Figure 3.**
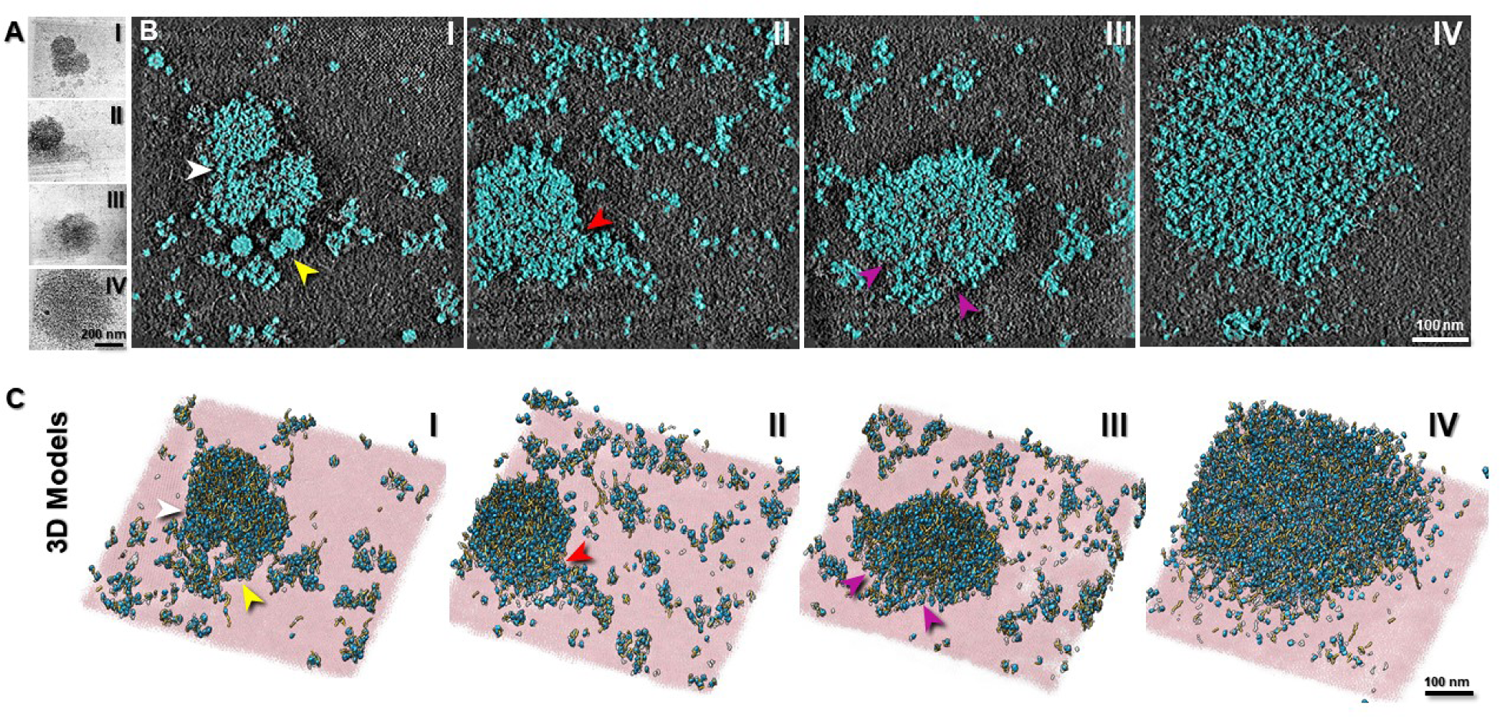
3D reconstructions of the initiation and growth of spherical condensates. Irregular and spherical condensates of different sizes formed at a tetranucleosome concentration of 30 nM, after 10 min of incubation at 20°C in physiological salt (150 mM Na^+^ and 5 mM Mg^2+^). (A) Enhanced cryo-ET zero-tilt images. The tomograms were organized by increasing size of the spherical condensates and labeled I, II, III, and IV. (B) Tomogram slices (1.17 nm thickness) of the same regions as in (A) depicting the NCPs (cyan) identified by the deep learning-based segmentation method. The tomograms show spinodal and spherical condensates of different sizes. The thickness of the tomograms is in range of ∼90-140 nm. (C) 3D models of the tomograms in (B) showing the fitted NCPs (cyan) and the flanking DNA (yellow). The SA-crystal is shown in pink and nucleosome density in transparent gray. The series are interpreted as illustrative of the possible mechanisms of growth of spherical condensates: (I) formation of small, near-spherical ∼35 nm condensates (yellow arrow) and the merging of three larger spherical condensates (∼100 nm, white arrow); (II) accretion of spinodal condensates by a ∼200 nm spherical condensate (red arrow); (III) a ∼300 nm spherical condensate resulting from the apparent fusion (purple arrows) of two smaller condensates; (IV) a larger spherical condensate of ∼400 nm likely arising from similar growth processes.

To characterize the internal structure of these spherical condensates and to establish how it relates to that of the surrounding spinodal condensates, we mapped out the NCPs in areas containing both types of structures (cyan, Fig. 3B). Because under fluorescence microscopy liquid droplets increased their size along the course of the condensation reaction, we used representative spherical condensates ordered by increasing size to illustrate the possible early stages of droplet formation and growth, both in segmented 2D slices (Fig. 3B; panels I, II, III, IV) and in the corresponding 3D models (Fig. 3C; panels I, II, III, IV). The panels in figures 3B and 3C also depict the likely fusion between spherical condensates of small (Fig. 3B and 3C; panel I, yellow arrows) and intermediate sizes (Fig. 3B and 3C; panel I, white arrows). However, the images indicate that growth of these spherical condensates was likely to occur not only by fusion among themselves, but also by accretion of spinodal material, as reflected by the presence of irregular condensates attached to spherical ones (Fig. 3B and 3C; panel II, red arrow). Panel III in figures 3B and 3C illustrates the possible merging (purple arrow) of smaller condensates to form a larger one while maintaining its spherical morphology. Such process suggests a mechanism by which even larger spherical condensates are generated (∼400 nm) (Fig. 3B and 3C; panel IV).

The small, rounded condensates (Fig. 3A; panel I, yellow arrows) were comparable in size to their spinodal counterparts and exhibited a near perfect spherical geometry. We propose that these small spherical condensates result from an early nucleation step that eventually gives rise to larger condensates. To confirm this inference, we analyzed the sample after 20 min of incubation, since nucleus formation is a rare event at early times (Sear, 2014) and the likelihood of capturing them increases with longer incubation periods (Fig. S18B and S19). Under these conditions, we observe small spherical condensates, either isolated or in groups of two or three (Fig. 4A and 4B). Statistical analysis indicates that the probability of finding a group of small spherical condensates together is similar to that of finding them isolated (Fig. 4C). This observation suggests that condensates appearing in clusters all likely arose from the same common large local concentration fluctuation within a spinodal condensate.

**Figure 4.**
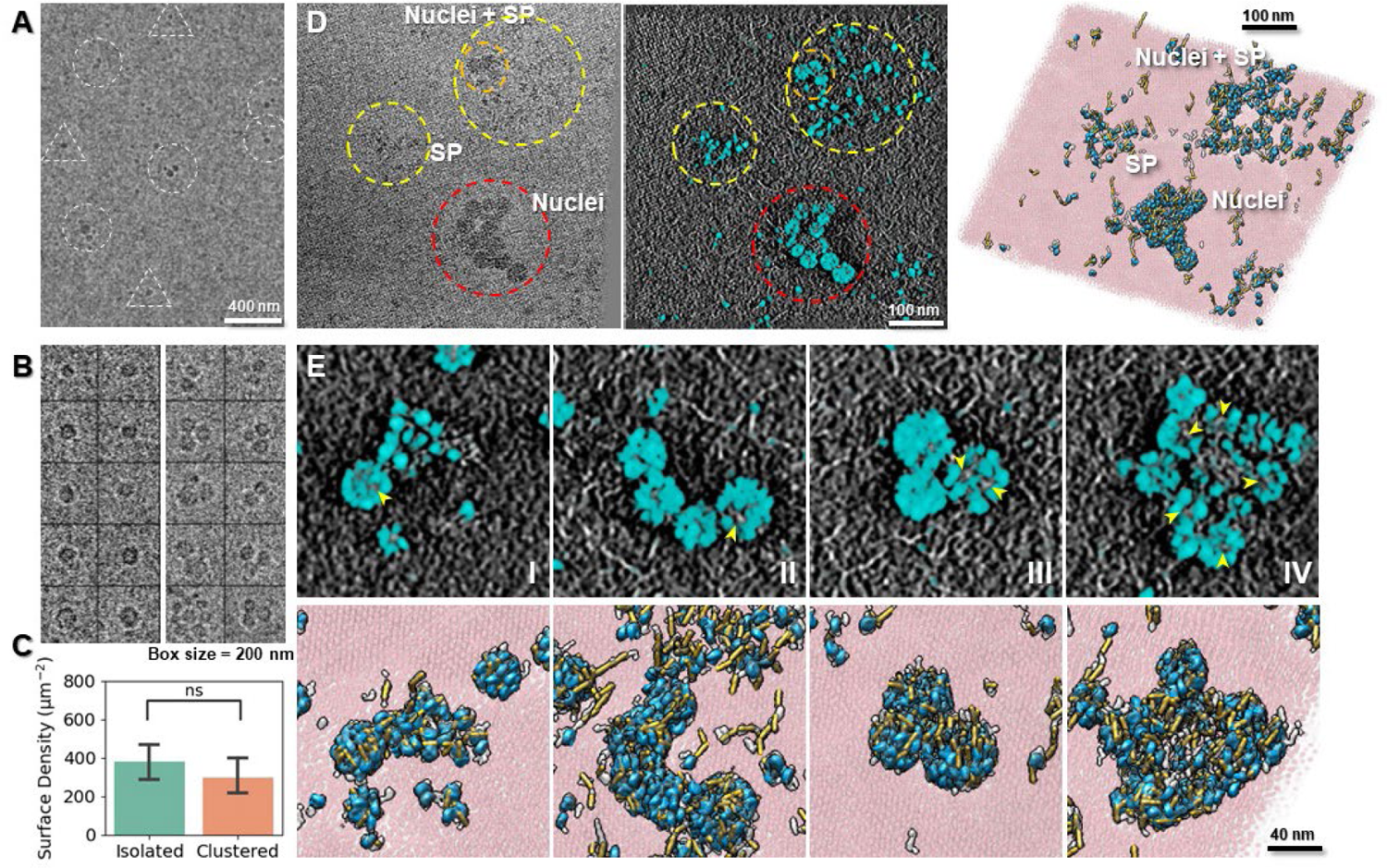
Small spherical nuclei arise from spinodal condensates. (A) Low-magnification images of single (dashed triangles) and multiple (dashed circles) small spherical condensates formed at 20°C in physiological salt after 20 min of incubation. (B) Representative images of individual (left columns) and clustered (right columns) small spherical condensates observed on the SA-crystal surface. (C) Density statistics measured from 12 low-magnification images indicating a similar chance to find condensates in clusters or in isolation. Data plotted as mean ± SEM. ns, not significant. (D) High-magnification cryo-ET zero-tilt image (left panel), tomogram slice (1.17 nm thickness; middle panel), and 3D model (right panel) of a cluster of small spherical condensates (red dashed circle), spinodal condensates (SP; yellow dashed circles), and small spherical condensates in close proximity with spinodal condensates (orange dashed circle). (E) Different stages of the nucleation and growth process (I through IV) of the spherical condensates. The top panels are representative tomographic slices (thickness of the tomograms ∼70 nm) and the bottom panels corresponding 3D models for the four states: (I) conversion of part of a spinodal condensate into a small nucleus; (II) multiple nuclei in side-by-side contact; (III) small nuclei organized in a cluster; (IV) cluster nuclei with smeared boundaries in the process of fusion. The yellow arrows indicate the irregular shape cavities within the small nuclei.

A zoomed-in view of the z-dimensional slices and their corresponding 3D models showed that some of the small spherical condensates were connected to and surrounded by irregular spinodal condensates (Fig. 4D). Figure 4E panel I, shows an example in which only half of the condensate was rounded and made up of 28 NCPs, while the other half, with 36 NCPs, was less organized. The total number of NCPs was similar to that of the late irregular condensates (∼60 NCPs) (Fig. 2E). These hybrid structures were also seen in the OpNS experiments (Fig. S3C, white arrow). While in early reaction times irregular or spinodal condensates were present without spherical condensates, the converse was not true: spherical condensates were only observed among irregular condensates. Together, these observations suggest that the small spherical condensates emerged from the pre-formed irregular condensates, and served as the nuclei that can then grow into larger spherical condensates, either via fusion among themselves and/or from accretion of surrounding spinodal structures.

Analysis of the size distribution of all spherical condensates in the 2D micrographs collected at 20 min of incubation revealed two peaks centered at diameters of 60 ± 14 nm and 161 ± 66 nm, while the smallest spherical structures observed had a diameter of ∼35 nm (Fig. S20). We suggest that the first peak represents the formation, due to large local concentration fluctuations, of thermodynamically unstable initial nuclei. The valley between the two peaks might indicate the dimensions beyond which the initial nuclei begin to overcome the interfacial energy barrier to (irreversibly) grow into larger and more stable structures. (Fig. S20). The probability distribution peak at 160 nm is likely the result of two opposing factors: one favoring the formation of larger condensates by the extra thermodynamic stability gained via the accretion of surrounding spinodal material (Fig. 4D) and/or the fusion with smaller condensates, and another disfavoring further growth because of the depletion of nearby material. The smallest spherical nuclei, although tightly packed, also contained regions of randomly located cavities (Fig. 4E; top panel, yellow arrow).

As before (Fig. 3B), we illustrate a possible temporal evolution and growth of nuclei by assembling representative spherical condensates (droplets) and ordering them by increasing size (Fig. 4E; panels I, II, III and IV). After the earliest stage in which small unstable nuclei are formed (Fig. 4E; panel I), these are often seen forming either linear or cluster arrangements (Fig. 4E; panels II and III); it is possible that the latter are favored over the former by the reduction in interfacial surface, a process that, upon fusion, will lead to a more favorable (smaller) surface-to-volume ratio. However, the high probability of observing clusters composed of spherical nuclei without fusion might be a manifestation of a secondary energy barrier for nucleus fusion. Larger nucleosome droplets result once this barrier is overcome (Fig. 4E; panel IV). Taken together, these observations indicate that nucleosomal phase separation occurs through a two-step process, the first involving organization of condensates by spinodal decomposition and the second in which the spinodal structures undergo re-organization into spherical, tight condensates (See Discussion).

### H1 Catalyzes the Spinodal-to-Spherical Condensate Transition

Linker histone H1 regulates chromatin compaction and packaging (Andreyeva et al., 2017; Song et al., 2014; Zhou et al., 2016). We observed using epifluorescence microscopy that H1 (120 nM, corresponding to a 1 to 1 ratio with the nucleosome concentration) in physiological salt and 20°C increased the number and size of nucleosome droplets compared to the ones formed without H1 (Fig. 5A). This is consistent with previous observations that H1 promotes nucleosome phase separation, and increases the concentration of nucleosomes in the liquid droplets (Gibson et al., 2019). To determine how H1 affects the two-step mechanism of nucleosome phase transition at early times, we used OpNS and cryo-ET imaging to investigated the architecture of H1-condensates at molecular level.

**Figure 5.**
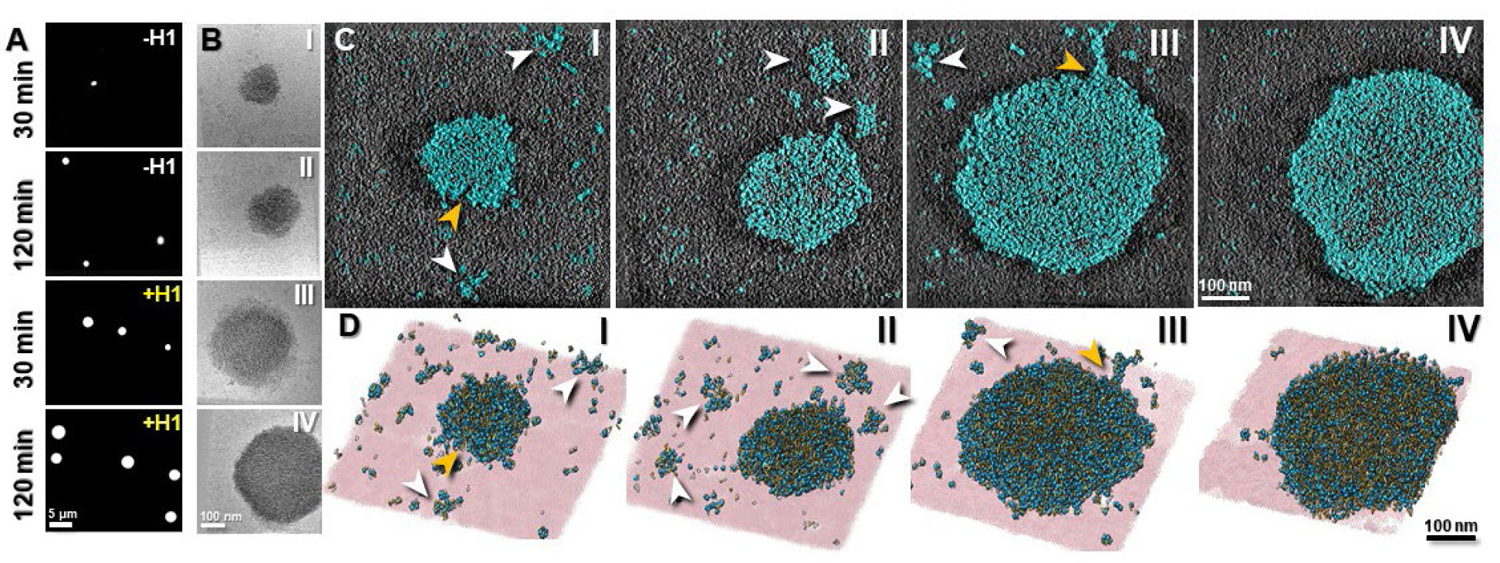
Linker histone H1 accelerates the transition from spinodal to spherical condensates. (A) Fluorescence microscopy images of Cy3 labeled tetranucleosomes incubated at 20°C in physiological salt with and without H1. (B) Representative zero-tilt cryo-ET images of spherical condensates obtained in the presence of H1 at 20°C in physiological salt after 10 min of incubation. (C) Corresponding tomogram slices (thickness 1.17 nm); and (D) 3D models of the corresponding condensates (tomogram thickness in the range of ∼90-130 nm). Tomograms are labeled I, II, III, and IV according to the size of spherical condensates, representing different stages of growth of spherical condensates. White arrows indicate the sparsely distributed spinodal condensates in the surrounding areas. Orange arrows indicate fusion events between condensates.

OpNS imaging showed that H1 induced the formation of large, micron-size spherical condensates at the earliest reaction time (2 min) (Fig. S21A), comparable to those observed after 60 min of incubation under the same conditions but without H1 (Fig. S2A). Next, we compared the surface number density of spinodal material in the presence and absence of H1 in areas devoid of spherical condensates from 2 min to 90 min (Fig. S21B). In the absence of H1, the number density of NCPs decreased only by ∼7% after 90 min of incubation (Fig. S4B), but in the presence of the linker histone, this parameter decreased exponentially to one-third of its initial value during the same time period (Fig S22B). Same incubation time series were also repeated at 4°C and 36°C in the presence of H1, which showed a similar trend but at lower rates (20°C > 36°C > 4°C, Fig. S22B). This result is consistent with the observation that the rate of spherical condensates formation does not monotonically depend on the temperature of incubation, being maximum at 20°C followed by 36°C and then by 4°C without H1 (Fig. S2 and S4A). Interestingly, a similar non-monotonic behavior with temperature has been described for tau protein condensates (Dong et al., 2021).

Using cryo-ET, we captured spherical condensates formed in the presence of H1 after 10 min of incubation at 20°C in physiological salt. Cryo-ET reconstruction of these condensates (Fig. 5B, 5C, and 5D) showed that they seem to have a tighter nucleosome packing (Fig. 5C; panel III) than those obtained under the same conditions in the absence of H1 (Fig. 3A; panel IV). The central z-dimensional slice of the reconstruction and the corresponding 3D models shown in figures 5C and 5D, respectively, display spherical condensates that appeared to be in the process of fusing with other spherical condensates or accreting spinodal material (Fig. 5C and 5D, yellow arrows). The sparse distribution of spinodal condensates in figures 5C and 5D, compared to their larger abundance around spherical condensates formed in the absence of H1 (Fig. 3B and 3C), once again supports our inference that H1 catalyzes the conversion of spinodal into spherical condensates without preventing the formation of the spinodal phase.

### Nucleosome Organization Inside the Spinodal and Spherical Condensates

We used the spatial coordinates of the individual NCPs to investigate the structural arrangements of NCPs inside the different types of condensates and then calculated the internal density of NCPs inside the condensates and their inter-nucleosome distance distribution. In order to determine the concentration of NCPs in the condensates, we first defined a series of 3D contours with the shape of each condensate, separated by a distance of 2 nm and placed at the center of mass of the condensate (See details in Materials and Methods, Fig. 6A; left panel). To calculate the nucleosome concentration within contour *n* (internal volume concentration), we divide the number of nucleosomes inside this contour by the volume it encloses (Fig. 6A; right panel). The difference of the number of nucleosomes enclosed by contours *n* and *n*-1 divided by the difference in their volumes, gives the nucleosome concentration of shell *n* (shell concentration). Next, we average the concentrations enclosed by the innermost contour, defined as contour *n=* 1 for all condensates of the same type (Fig. 6B). Then we repeated this procedure to calculate the average concentration enclosed by contour *n=* 2, and so on. We also obtained the average concentration of NCPs inside shell *n=* 1, 2, 3, etc. This analysis was performed only on condensates whose dimensions were similar or smaller than ∼150 nm, since they are not severely flattened due to thickness of the vitreous ice.

**Figure 6.**
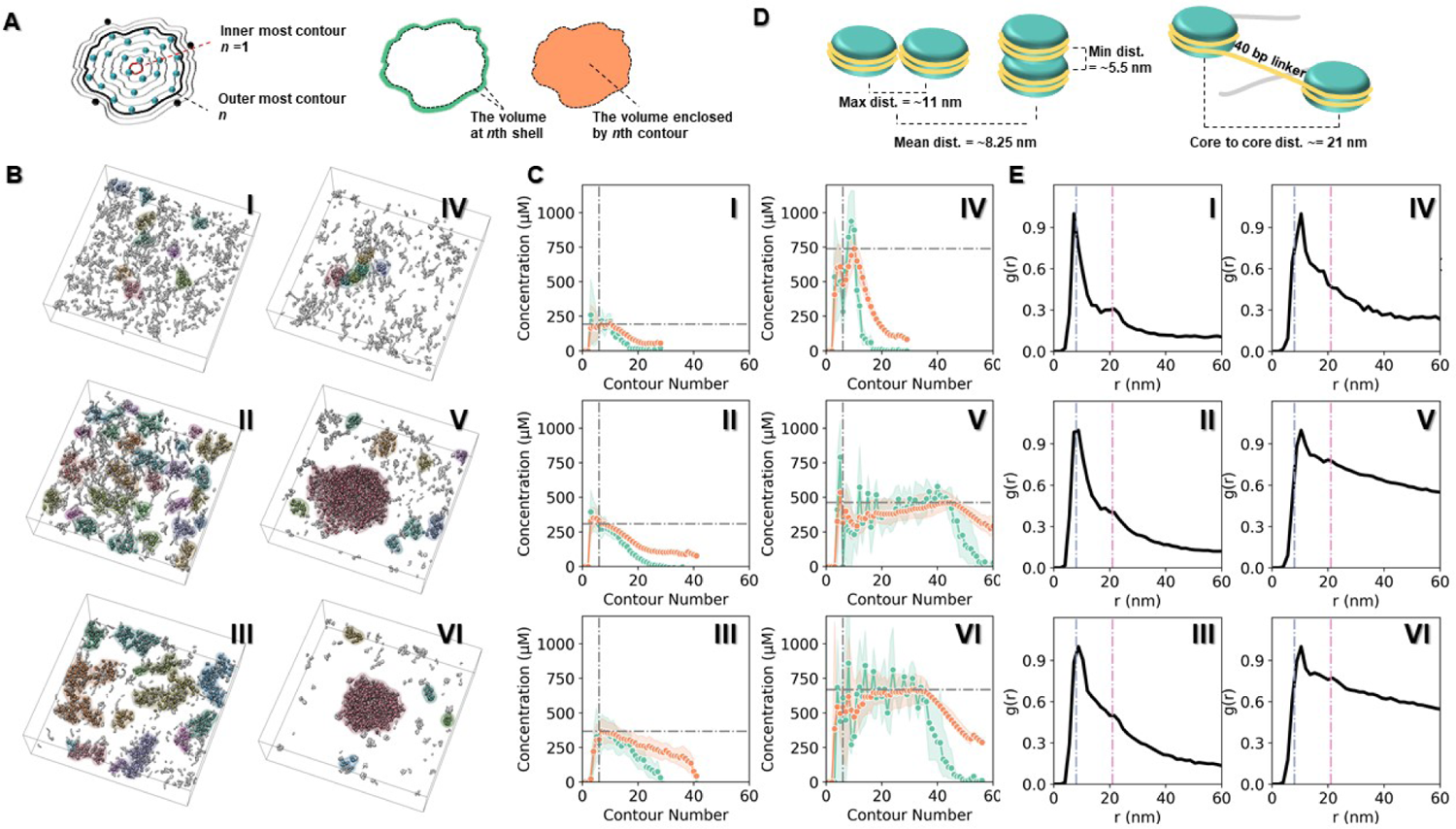
Quantitative analysis of NCP concentration and organization during different stages of condensate growth. (A) Schematic showing concentric contours, 2 nm apart and following the condensate shape, with the innermost contour labeled *n*=1 (left). Cyan and black dots represent internal and free NCPs around the condensate, respectively. Depiction of the *n*th-shell delimited by contours *n* and *n*-1 (green region) and the volume the *n*th contour encloses (orange region) (right). The nucleosome *n*th-shell concentration is defined as the number of NCPs inside shell *n*, divided by the corresponding shell volume. The *n*th-volume concentration is defined as the number of NCPs inside contour *n,* divided by the volume it encloses. (B) The shape of contour series was defined by the 10 nm low-pass maps shown in different colors. The numbering of the maps follows the proposed two-step phase separation mechanism from early (I), intermediate (II), and late stage (III) spinodal condensates to the small nuclei (IV) and spherical condensates without (V) and with (VI) H1. (C) Quantitation of the *n*th-shell (green) and *n*th-volume (orange) nucleosome concentration, corresponding to stages I through VI. The vertical and horizontal dash lines mark the highest contour concentration when *n* > 5. (D) Theoretical inter-nucleosome distances for side-to-side (11 nm) and stacked (5.5 nm) configurations (left), and distance between neighboring NCPs within a tetranucleosome (21 nm) (right). (E) Radial distribution function g(r) from NCPs’ centers of mass for stages I through VI. The blue and pink dash lines indicate peaks at 8 nm and 21 nm, respectively. The g(r) calculation for spinodal condensates (I, II, and III) was obtained from all the NCPs within the ice slab. The g(r) calculation for spherical condensates (IV, V, and VI) excludes the surrounding spinodal condensates.

We examined both the shell and internal volume concentration starting from *n* > 5, because both concentrations cannot be not well-defined when the measured volume approaching the size of a single NCP. The shell concentrations for *n*= 6 obtained for the initial, intermediate, and late stage spinodal condensates were 217 ± 43, 274 ± 36 and 356 ± 66 µM, respectively, and the internal volume concentration enclosed by contour *n*= 6 were 195 ± 25, 309 ± 21 and 364 ± 47 µM, respectively (Fig. 6C; left panels). The shell concentrations for all three stages of spinodal condensates decreased towards the periphery of the condensates, and vanished for shell *n*= 17, 24, 28 for the three stages, respectively (Fig. 6C; left panels). In comparison, the initial spherical nuclei, the spherical condensates, and the spherical condensates in the presence of H1, all displayed overall higher shell concentrations than their spinodal counterparts at shell *n=* 6, yielding values of 286 ± 97, 339 ± 74 and 612 ± 152 µM, and correspondingly higher internal volume concentrations enclosed by contour *n*= 6 of 494 ± 68, 402 ± 45, and 550 ± 138 µM, respectively (Fig.6C; right panels). Notably, unlike the spinodal condensates, the spherical condensates did not display a decrease but instead a slight increase in shell concentrations towards the periphery, reaching a peak shell concentration of 939 ± 99, 501 ± 36 and 688 ± 65 µM and a peak internal volume concentration of 743 ± 64, 471 ± 42 and 667 ± 62 µM, for the initial spherical nuclei, the spherical condensates, and the spherical condensates formed in the presence of H1, respectively (Fig.6C; right panels). Thus, the concentration of the spherical condensates is ∼1.3 to 3.6-fold higher than that of the spinodal condensates, and increases significantly in the presence of H1. Notably, the condensates with the highest concentrations were found to be the initial small nuclei. As these nuclei grow by accretion or fusion, their initial cores must rearrange so that the resulting spherical condensates display more concentration towards the periphery than at their center. Interestingly, the large discrepancy between the peak shell and peak volume concentrations (∼200 µM) observed in the spherical nuclei suggests that formation of a nucleus involves an outer shell of tightly packed NCPs with an interior somewhat lower in NCP concentration; this conclusion is supported by z-dimensional slice images of nuclei, clearly displaying these features (Fig 4 D and 4E).

Next, in order to characterize the spatial organization of individual NCPs within the condensates, radial distribution functions g(r) centered at each NCP were calculated and the results were averaged over all the particles in condensates of the same type in the data set (See Materials and Methods). This analysis reveals two characteristic peaks residing at distances of ∼8 nm and ∼21 nm (Fig. 6E, dash lines). The first peak, which corresponds to the distance between nearest NCPs centers, moves from 7.5 nm, to 8.1 nm, to 9.0 nm for the early, intermediate and late spinodal condensates, respectively. The value of 7.5 nm can be rationalized as the distance between two NCPs averaged from their various contact conformations. The stacking conformation of *n* and *n*+2 NCPs within the same array (Fig 6D; left panel) and the side-by-side conformation adopted by NCPs belonging to neighboring arrays (Fig 6D; left panel) delimits the minimum and maximum core to core distance of 5.5 nm and 11 nm, respectively. The rest of the NCPs’ contact conformations should contribute distances that lie somewhere in between the above two extremes. (Bilokapic et al., 2018). The small shift in the peak distance as the spinodal condensates mature and grow might reflect a decrease in the statistical weight of intra-array *n*/*n*+2 stacking in favor of inter-array interactions. Similarly, the early spherical nuclei and the grown spherical condensates, both with and without H1, also display a first peak at 10.6 nm, 10.5 nm, and 10.6 nm, respectively (Fig. 6E, blue dash line). These increased peak values relative to those found for the spinodal condensates probably indicate that the higher compaction in spherical condensates further discourage intra-array *n*/*n*+2 stacking in favor of inter-array NCPs interactions, a conclusion supported by chromatin dynamics simulations (Farr et al., 2021). The second peak position (Fig. 6E, pink dash line), which appears as a shoulder, matches the distance expected between *n* and *n*+1 NCPs that are connected by a 40 bp DNA linker in every array (Fig. 6D; right panel). Note that the prominence of this peak decreases monotonically from the early spinodal to the grown spherical condensates. This trend may reflect the bending of the linker DNA within an array, and/or the increased statistical weight of neighboring nucleosomes belonging to different arrays as a result of the condensation process. Overall, the g(r) decay trend agrees well with the concentration measurements in the spinodal and spherical condensates: the high-density spherical condensates have a slower g(r) decay, indicating a higher chance to find neighbor NCPs at a close distance when compared to the early spinodal condensates.

The strong tendency of nuclei to adopt a spherical shape indicates that their formation is accompanied of an unfavorable nucleosome-water surface interaction energy. Accordingly, we investigated if the orientation of NCPs on the surface of these condensates differs from those in the interior. For a given nucleosomal disk in the spherical condensates, we determined the angle θ between each fitted discoidal plane and the radius of the condensate that crosses the center of the disk. For the spinodal condensate, we determined the same angle between the fitted discoidal plane and a line perpendicular to the central axis of the condensate (its ‘skeleton’, see Material and Methods) that passes through the center of the nucleosomal disk. If the nucleosomal disks are randomly distributed, the probability density distribution of their angular orientation should follow a cosine distribution, *P*(θ)*d*θ = *cos*(θ)*d*θ (see Materials and Methods). Deviation from this distribution indicates a preferential orientation of the nucleosome disks either on the surface or in the interior of the condensates. The results of this analysis (Fig. S23) showed that the nucleosomes in the interior exhibited very small deviations from the random distribution profile in the spinodal condensates and in the spherical condensates with or without H1. Only the nucleosomes in the exterior of the initial spherical nuclei displayed a moderate skewed angle distribution, where the disks tended to orient more parallel to the condensates’ surface. In contrast, the NCPs at the surface of the spinodal condensates, the spherical condensates, and the spherical condensates formed in the presence of H1, all showed an increased tendency to align perpendicular to the condensate surface (θ = 0). Again, only for the initial nuclei, the NCPs at the surface tended to align parallel to the condensates’ surface (θ = π/2). This analysis indicates that the energetic and global structural differences between spinodal and spherical condensates do not arise from a distinct spatial arrangement of the NCPs within the respective condensates. Instead, the transition between these two condensed forms must involve a change in the nature of the inter-nucleosome and nucleosome-solvent interactions (See discussion).

## Discussion

We have used cryo-ET to capture, for the first time, the 3D molecular organization of tetranucleosome arrays during the initial stages of their phase separation, when the condensates are only a few tens of nanometers in size. The identification of the resulting condensates and their structural characterization was possible because of the large dimensions of the nucleosome compared to other biological systems that also undergo phase separation, and the use of an optimized 3D reconstruction method. We have obtained snapshots of condensates at different times (2 min, 10 min, and 20 min) of incubation in physiological salt conditions (150 mM Na^+^ and 5 mM Mg^2+^), from which it has been possible to infer their likely process of growth and maturation. We show that nucleosome phase transition occurs through a two-step mechanism (Fig. 7 and movie S1). In the first step, dispersed nucleosomes (at a concentration of 120 nM) first experience a fast and system-wide condensation resulting in the ubiquitous appearance of irregularly shaped clusters. These condensates attain a nucleosome concentration of ∼360 µM and average dimensions of (∼125 nm x 65 nm x 30 nm). During their formation, they grow by enlarging their major axis, while their minor axes remain nearly constant. This step closely resembles a spinodal decomposition process in which aggregate formation occurs throughout the system without the crossing of an energy barrier and where domain anisotropy arises as a consequence of diffusive effects with a driving force that tends to minimize the surface tension between the two phases (Datt et al., 2015). According to the Cahn-Hilliard theory of spinodal decomposition (See supplementary theory section), the distance among the resulting condensates are determined by a “critical wavelength λ” that describes the spatial pattern of separation between the two phases. This wavelength, in turn, is given by the ratio of the parameter that describes the cost of creating a concentration fluctuation in the medium and the “curvature” of the free energy surface of the system with respect to the concentration of the dispersed and the dispersing phases (see supplementary theory section, equation 9). We do not know what factor limits the length of the spinodal condensates along their longer axis. It may simply reflect the limit of mechanical stability of condensates made up of small tetranucleosome elements or the length attained before the concentration of free arrays falls below a critical value for the condensates to continue to grow. In the second step, the system undergoes a second condensation process in which highly packed spherical nuclei emerge locally and sparsely from the existing spinodal condensates. This step resembles a more traditional nucleation-and-growth process in which an initially unfavorable event (nucleation) is increasingly stabilized by a favorable volume energy through the formation of nucleosome-nucleosome interactions distinct to those adopted in the spinodal condensates. These new inter-nucleosomal interactions must be accompanied, however, by an unfavorable surface energy with the solvent, which forces the system to adopt a spherical shape and minimize its surface-to-volume ratio (see below). Throughout their growth, the initially small unstable nuclei become increasingly stabilized through a process of accretion of the surrounding spinodal material, or by fusion with other spherical nuclei. We identify that the phase separation described here is a two-step process—as opposed to a simple spinodal phase that undergoes coarsening or Ostwald ripening (Ostwald, 1897)—because we observed a clear separation of timescales between the formation of the two phases. In detail, the spinodal decomposition occurs and achieves a steady state, indicated by the arrested growth of condensates’ dimensions before 10 min, 2 min, and 10 min for the 4°C, 20°C, and 36°C reaction (Fig S2 and S4D), respectively. After that, a secondary nucleation process emerges within those spinodal condensates, which was observed after 60 min, 2 min, and 10 min for reaction at 4°C, 20°C, and 36°C (Fig S2 and S4C), respectively. In contrast, a process of Ostwald ripening will occur continuously with the formation of the matured spinodal condensates without a clearly defined time scale separation from the initial spinodal phase and without a nucleation step.

**Figure 7.**
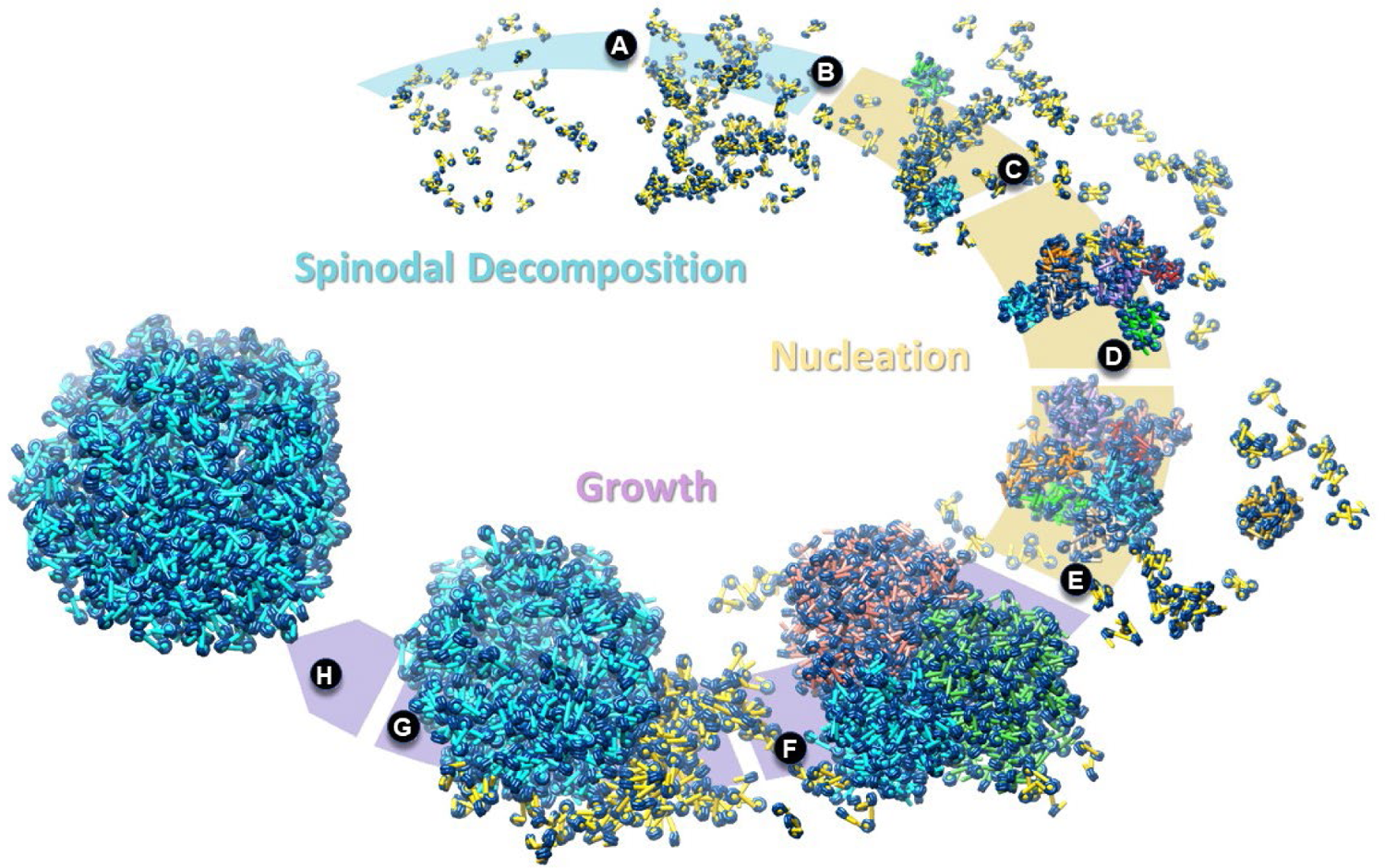
Proposed two-step mechanism of tetranucleosome phase separation. (A) Initial, dispersed phase of tetranucleosome arrays; (B) organization of tetranucleosome into irregular, spinodal condensates; (C) emergence of spherical nuclei within the spinodal phase and growth of these nuclei from accretion of the spinodal material; (D) bundle of spherical nuclei; (E) fusion of small nuclei bundles enable growth into larger, energetically stable spherical condensates; (F) fusion between spherical condensates; (G) growth of the spherical condensates by accretion of surrounding spinodal material; and (H) large spherical condensate corresponding to the early stages of liquid droplet. The light blue, pale brown, and purple background indicate the major steps of nucleosome condensation: spinodal decomposition, spherical nuclei formation, and growth of spherical condensate, respectively. H1 catalyzes step (C) through (H). Spinodal condensates are shown in yellow, and the spherical condensates are shown in blue, green, orange, pink, and red. All states are illustrated using experimentally observed condensates.

The 3D mapping of NCPs obtained here by using deep learning segmentation, missing wedge correction, and a spatial clustering algorithm, reveals that the small nuclei formed at the beginning of the second condensation step are very dense, attaining a nucleosome concentration of ∼740 µM and involving the tight packing of a large number of NCPs within a spherical volume as small as ∼35 nm in diameter. The sparse appearance of these spherical nuclei suggests that their formation is both kinetically and thermodynamically inaccessible from the initial solution of nucleosome arrays, and that they only arise from the pre-existing spinodal condensates as a result of rare, large local concentration fluctuations, that are capable of overcoming a significant energy barrier. Given the localized and rare appearance of these nuclei, it is somewhat surprising that we often observe them close together forming clusters. It is statistically unlikely that these condensates arose at widely separate loci in the medium and came together to form a bundle. Instead, it is more likely that large, localized concentration fluctuations gave rise to multiple nucleation sites at a given location. Because the conditions under which we observe the formation and growth of spherical nuclei by cryo-ET also leads to the observation of large spherical condensates (liquid droplets) by fluorescence microscopy, we propose that the spherical condensates described here reflect the early stages in the formation of those droplets.

We find that the transition from spinodal to larger spherical condensate is not accompanied by a significant difference in the orientation of NCPs relative to one another or to the outer surface of the condensates. This observation implies that this transition, which is accompanied by an approximately 2-fold increase in nucleosome concentration, involves a qualitative change in the interaction of the nucleosomes among themselves and with the solvent. Accordingly, to rationalize our observations we propose a model in which, in the spinodal process, the condensates retain significant amount of solvent, and the contacts of nucleosomes among themselves and with the solvent are mediated by polar interactions. These condensates do not experience a surface energy cost, as the difference between the interactions among nucleosomes in the interior and on the surface are not qualitatively different, only quantitatively so, as revealed by their angular orientation. During the nucleation step, in contrast, solvent is significantly excluded, and it is likely that hydrophobic nucleosome surfaces are exposed that mediate the nucleosome-nucleosome and nucleosome-solvent interactions. Indeed, Swi6, an HP1-family protein in *S. pombe*, has been shown to induce liquid-liquid phase separation nucleosomes by promoting a conformational change that increases the solvent accessibility and dynamics of hydrophobic residues isoleucine, leucine and valine of the octamer core (Sanulli et al., 2019). Based on these observations, these authors have proposed that nucleosome plasticity is key for the action of Swi6, favoring nucleosomal states that promote liquid-liquid phase separation by increasing multivalent interactions between nucleosomes. Other authors have also called attention to the importance of hydrophobic residues as adhesive elements in liquid-liquid phase separation (Alberti et al., 2019). Consistent with these observations, treatment of 1,6-hexanediol (which inhibits weak hydrophobic interactions) to living cells caused a more uniformed chromatin distribution (Ulianov et al., 2021) and increased chromatin accessibility (Henikoff et al., 2020). While thermodynamically favorable in the interior, these interactions result in an unfavorable energy cost for nucleosomes at the surface, forcing the condensates to minimize the area exposed to the solvent, by adopting spherical shapes. Interestingly, the isotropic distribution of the NCPs disks observed in the spinodal and the larger, stabilized spherical condensates, indicate that these condensates do not possess the well-defined structure of a solid, but rather the random orientation associated with a liquid-like state.

The observation of a two-step phase separation pathway in the formation of large spherical phase droplets, as opposed to a conventional one-step (classical) nucleation mechanism, has precedence in the literature. Examples include the phase separation observed in inorganic systems such as nanogold particles, Fe-Mn alloys, and in molecules of biological origin, such as ferritin (Houben et al., 2020; Kwiatkowski da Silva et al., 2018; Loh et al., 2017a).

Along the cell cycle, chromatin exhibits different levels of compaction, from a less condensed state at interphase, to a highly condensed state of the chromosomes during mitosis. The internal concentration of the irregular spinodal condensates formed by the tetranucleosome arrays ranged from ∼195 µM (early spinodal stage) to ∼360 µM (late spinodal stage), which are in the range of the *in vivo* chromatin concentration estimated for late interphase and the beginning of mitosis, respectively (Hihara et al., 2012). However, if we consider the characteristic spacing *λ* that is generated concomitantly with the formation of spinodal condensates, the local nucleosome concentration decreases down to ∼30-50 µM. This observation suggests that the spinodal condensates could represent the less constrained and active form of chromatin at interphase (DNA replication and transcription). The irregular organization of spinodal condensates supports the current model of interphase chromosome organization, in which the 10-nm fibers self-interact and condense to form a polymer-melt structure (Hansen et al., 2018). The average dimensions of the spinodal condensates (∼55-125 nm x ∼30 nm for the long and short axis, respectively) suggest an inherent tendency of nucleosomal arrays to spontaneously self-assemble into a geometry whose short axis may explain the earlier description of a regularly packed 30-nm chromatin fiber (Garcia-Saez et al., 2018; Grigoryev et al., 2009; Robinson and Rhodes, 2006; Song et al., 2014). Note, however, that regular 30-nm chromatin fibers have not been observed inside the nucleus (Eltsov et al., 2008; Maeshima et al., 2014b; Ou et al., 2017; Razin and Gavrilov, 2014). Consistent with these observations, we found only one condensate displaying a regular structure out of the great majority of irregularly packed spinodal structures (Fig. S3C; left column second row).

The transition from the spinodal condensates to droplets involves the formation of a spherical condensate, and a ∼1.3-fold increase in nucleosome concentration (∼470 µM) as determined here by cryo-ET (Fig. 6). We note that this high nucleosome concentration is in the range estimated for metaphase chromosomes (Hihara et al., 2012), and that it is similar to both the liquid droplet nucleosome concentration determined *in vitro* by fluorescence microscopy using 12-mer nucleosomes arrays (∼340 µM) (Gibson et al., 2019) and to the concentrations predicted by Monte Carlo simulations (∼500 µM) (Maeshima et al., 2016). It is interesting to estimate how an increase in nucleosome concentration affects the ability of a molecule such as RNA Polymerase II (MW= 550 kDa) to diffuse inside the condensates. Empty chamber evaluation analysis (Fig. S24) shows that concomitant to the increase of internal nucleosome concentration, there is a nearly 10-fold decrease in the number of Pol II molecules that could fit in these structures, from the late stage spinodal condensate to the spherical condensate without H1, and then to the spherical condensate in the presence of H1. This analysis also indicates that spherical condensates, especially those formed in the presence of H1, should strongly repress transcription initiation, given the even larger dimensions of the transcription pre-initiation complex. In contrast, smaller molecules may still be able to diffuse inside liquid droplets, since experiments have shown that linker DNA within the liquid droplets can be readily cleaved by micrococcal nuclease (17 kDa) (Maeshima et al., 2016) and by Cas12 (120 kDa) (Strohkendl et al., 2021). Similarly, Tet Repressor (23 kDa) was shown to be strongly recruited to nucleosome droplets in the presence of its operator (Gibson et al., 2019).

Formation of the early spherical nuclei requires at least 10 min of reaction under physiological salt conditions at 20°C, and it is accompanied by an ∼2-fold increase in nucleosome concentration (∼740 µM). The difficulty of stochastically generating tightly packed nuclei within the spinodal condensates indicates that this process is the rate limiting step in the formation of LLPS which, in turn, makes it a possible target of regulation for chromatin condensation in the cell. Accordingly, conversion of chromatin condensation states in the cell from interphase to metaphase chromosomes could be controlled by the regulation of the intrinsic two-step nucleosome phase separation mechanism established here. Indeed, we find that incubation of nucleosome arrays with linker histone H1 catalyzes the rate-limiting step that leads to the formation of spherical condensates at the expense of the surrounding spinodal material (Fig. 5). Thus, H1 and other protein factors could function as catalysts that reduce the energy barrier required to transition from the spinodal to the spherical condensates, thus regulating the conversion of chromatin from interphase to metaphase. Likewise, because spinodal condensates have the intrinsic ability to condense into spherical condensates in physiological salt (Fig. 3), external factors such as proteins and chemical modifications may be needed to maintain an open chromatin state during interphase. This inference is supported by the observation that proteins such as FoxO1 are able to bind nucleosomes and induce chromatin opening (Hatta and Cirillo, 2007), and that histone acetylation has been shown to disrupt chromatin droplets (Gibson et al., 2019). Future efforts will be directed toward determining how histone acetylation and removal of core histone tails affect the early stages of nucleosome spinodal condensation and spherical nuclei formation.

Collectively, it is tempting to extrapolate our *in vitro* observations and propose that chromatin, under physiological conditions in the cell, can exist in an equilibrium between a loosely packed spinodal-like state, fully accessible for transcription, repair, and replication, and a highly packed spherical condensate arising from a process of nucleation-and-growth inside the spinodal structures. The former would be associated with transcriptionally active chromatin, while the droplet state would represent transcriptionally silent states. We propose that it is through a tight spatial and temporal regulation of the interconversion between these two forms that cell identity and differentiation are established, and physiological homeostasis is attained in response to external cues.

## Materials and Methods

### Purification of Histones Proteins

Purification of *Xenopus laevis* histones H2A, H2B, H3, and H4, was performed as previously described (Wittmeyer et al., 2004). Briefly, histone plasmids were transformed into *E. coli* BL21(DE3) and bacterial cells were grown while shaking at 37°C in LB media supplemented with ampicillin and chloramphenicol. Histone expression was induced at an OD600 of ∼0.6 by adding Isopropyl b-D-1-thiogalactopyranoside (IPTG) at a final concentration of 1 mM. After 3 h of induction, cell cultures were pelleted, suspended in wash buffer (50 mM Tris-HCl pH 7.5; 100 mM NaCl; 1 mM EDTA; 5 mM 2-mercaptoethanol (BME); 1% Triton X-100 [w/v]; and protease inhibitors (Roche)) and frozen at −80°C for later use. To purify the inclusion bodies, cell pellets were thawed, sonicated, and centrifuged at 25,000 x g for 30 min. Supernatant was discarded, and the pellet, corresponding to the inclusion bodies, was rinsed, suspended in wash buffer, and centrifuged three times. To remove Triton X-100, inclusion bodies were rinsed and centrifuged three times with wash buffer without Triton X-100. Inclusion bodies were solubilized with a buffer containing 20 mM Tris-HCl pH 7.5; 8 M Urea; 1 mM EDTA; 10 mM DTT, and purified by anion and cation exchange. Purification was checked by 15% SDS-PAGE and fractions containing pure histones were pooled and dialyzed against 1 L of 10 mM Tris pH 8.0, with three buffer exchanges. Histones were centrifuged to remove aggregates, concentrated by centrifugation (∼10 mg/mL), lyophilized, and stored at −80°C. The integrity and correct positioning of the four nucleosomes was corroborated using Atomic Force Microscopy (AFM) and Electron Microscopy (EM) (Fig. S1).

### Fluorescence Labeling of H2A

For fluorescence microscopy experiments, *Xenopus laevis* histone H2A119C (The Histone source, Colorado State University) was chemically modified with the fluorescent dye Cy3-maleimide (Lumiprobe) as previously shown, with some modifications (Zhou and Narlikar, 2016). Lyophilized H2A119C was resuspended to 2 mg/mL using labeling buffer (20 mM Tris-HCL, pH 7.0; 7 M guanidine hydrochloride; 5 mM EDTA) and incubated for 2 h at room temperature. Unfolded histone was reduced by the addition of 100 mM TCEP (Sigma) and incubated for 2 h in the dark with occasional mixing. A second round of 100 mM TCEP reduction was done overnight. Cy3-maleimide was dissolved with dimethylformamide (DMF) to a final concentration of 20 mM and H2A119C was labeled by the addition of 15-fold molar excess of Cy3 maleimide (∼2 mM). Labeling reaction was performed at 21°C overnight, in the dark, and with constant shaking. To remove non-conjugated Cy3-maleimide, the labeling product was concentrated by centrifugation using a Microcon of 3.5 kDa membrane cutoff. Concentrated Cy3-histone was diluted with labeling buffer and concentrated again. This procedure was repeated six times, and every other time, the Microcon filter was changed for a new one. Concentration of labeled histone and degree of Cy3 labeling was ∼50%, which was calculated by measuring the absorbance at 278 nm and 555 nm and using the extinction coefficients of 4,400 Mcm^-1^ and 150,000Mcm^-1^ for H2A119C and Cy3, respectively.

### Histone Octamer Reconstitution

To reconstitute the wild type histone octamer (Wittmeyer et al., 2004) and the Cy3-labeled octamer (H2A-Cy3 replaces H2A), lyophilized histones were solubilized with unfolding buffer (20 mM Tris-HCl pH 7.5; 7 M guanidine hydrochloride; 10 mM DTT), combined at a H2A:H2B:H3:H4 ratio of 1.2:1.2:1:1, and dialyzed 4 times against 1 L of refolding buffer (10 mM Tris-HCl pH 8.0; 2 M NaCl; 1 mM EDTA; 5 mM DTT), with three buffer exchanges, for a total of 48 h. Refolded octamer was centrifuged, concentrated to ∼0.5 mL and loaded onto Superdex 200HR (Amersham Bioscienes) equilibrated with refolding buffer. Gel filtration separated octamers from aggregates, tetramers, and dimers. Fractions were checked by 15% SDS-PAGE, and fractions containing the four histones in equimolar quantities (based on Coomassie blue staining) were pooled, concentrated to ∼10 mg/mL, and stored at −80°C. Fractions containing H2A-H2B heterodimer were also pooled, concentrated to ∼10 mg/mL, and stored at −80°C.

### Synthesis of Tetranucleosome DNA Template

The tetranucleosome DNA template array contains four repeats of the 601-nucleosome positioning sequence separated by a 40 bp linker length, and flanked to the left by 100 bp DNA, and to the right by 50 bp DNA (100-4W-50; left and right sides correspond to the 5’ and 3’ ends, using the 601 sequence (W) as a reference). Tetranucleosome DNA template was synthetized by ligation of four fragments containing one 601 sequence each. Each fragment was produced by PCR using primers containing the restriction recognition sites for enzymes BbsI and BsaI (NEB), which recognize asymmetric DNA sequences and cut outside of their recognition sequence. This allowed for the ligation of fragments in tandem and for control over the number of repeats. In our design, the tetranucleosome is flanked by two BsaI sites, and the BbsI sites were used to ligate the four nucleosome fragments. In the first cleavage step, the four fragments were digested with BbsI and ligated at an equimolar ratio using *E. coli* ligase (NEB). The longest product, corresponding to the tetranucleosome, was purified from 0.8% agarose gels, digested using the BsaI enzyme, and cloned into BsaI-restricted PGEM 601 plasmids using T4 ligase (NEB). Ligation product was transformed into *E. coli* DH5α for plasmid extraction by miniprep. The product of the tetranucleosome DNA synthesis was checked by DNA sequencing.

To produce large quantities of the tetranucleosome DNA template, the tetranucleosome plasmid was grown in *dam-/dcm-E. coli* cells (NEB) and was purified by maxiprep. Tetranucleosome DNA template was excised from the vector backbone by digestion using the BsaI enzyme. Tetranucleosome template was purified from the backbone by preparative acrylamide electrophoresis using a Model 491 Prep Cell (Bio-Rad).

### Synthesis of Biotinylated Tetranucleosome DNA Template

To incorporate a biotin molecule to the 100-4W-50 template without modifying its length, a 70-4W-50 template was ligated with a biotinylated 30 bp dsDNA to produce the biotin-100-4W-50 DNA template. The 70-4W-50 template was designed and produced in the same way as the 100-4W-50. The 30 bp dsDNA was synthesized by the annealing of two complementary oligos, forming a 5’ biotinylated end and a 3’ end with a BsaI overhang, which is complementary to the left side of 70-4W-50. Biotin-30 bp ds oligo and 70-4W-50 were ligated using T4 ligase at a molar ratio of 20:1, respectively. Ligation reaction was inactivated with SDS loading buffer (purple dye, NEB), and the biotin-100-4W-50 product was purified by preparative 7% acrylamide electrophoresis.

### In vitro Reconstitution of Tetranucleosomes Arrays

Wild type histone octamers and tetranucleosome DNA templates (100-4W-50 and biotin-100-4W-50) were combined at a ratio of 1:1.2, respectively, in 200 µL of high-salt buffer (10 mM Tris-Cl pH 8.0; 2 M NaCl; 1 mM EDTA; 0.5 mM DTT; and 1 mM PMSF) at a final concentration of 100 ng/uL of DNA template. H2A-H2B dimer was also incorporated at a molar ratio of 0.2 compared to the octamer to promote nucleosome, rather than hexasome formation. For fluorescent nucleosome arrays, wild type octamers were doped with 25% Cy3-octamer and combined with the 100-4W-50 template. Tetranucleosome assembly solutions were dialyzed against 500 mL of high-salt buffer for 1 h at 4°C, followed by a lineal gradient dialysis against 2 L of low-salt buffer (10 mM Tris-Cl pH 8.0; 1 mM EDTA; 0.5 mM DTT; and 1 mM PMSF) using a peristaltic pump at 0.8 mL/min and with continued stirring. A final dialysis of 3 h in 500 mL of low salt buffer was done to reduce the residual NaCl concentration, and the nucleosome reconstitution was checked by 4% acrylamide native electrophoresis.

Reconstituted arrays were loaded into 4.8 mL of 10-40% lineal sucrose gradient (20 mM HEPES-NaOH pH 7.5; 1 mM EDTA; 1 mM DTT) and centrifuged for 16 h at 38,000 rpm at 4°C, using an ultracentrifuge Beckman Optima MAX-XP with the rotor MLS-50 (Beckmann). Gradients were fractionated into 100 µL fractions using the Brandel gradient fractionator, and the resulting purification was checked by 4% acrylamide native electrophoresis. Fractions containing 1-2 bands were pooled, concentrated, and dialyzed against 20 mM HEPES-NaOH pH 7.5; 1 mM EDTA; 1 mM DTT.

### Atomic Force Microscopy of Nucleosome Arrays

Purified tetranucleosomes were diluted to 10 nM and crosslinked with 1% formaldehyde for 1 h at room temperature. Crosslinked sample was dialyzed against 20 mM HEPES-NaOH pH 7.5; 1 mM EDTA; 1 mM DTT, and centrifuged at 20,000 x g to remove aggregates. Crosslinked tetranucleosome sample was diluted to 1 nM using 10 mM MOPS pH 7.0 and 5 mM MgCl2, and 3 µL of sample were deposited and incubated for 2 min on freshly cleaved bare mica V1 (Ted Pella Inc.), after which was rinsed with Milli-Q water, and then gently dried under a stream of N2 perpendicular to the mica surface. AFM micrographs were taken with a MultiMode NanoScope 8 atomic force microscope (Bruker Co.) equipped with a vertical engagement scanner E. The samples were excited at their resonance frequency (280-350 kHz) with free amplitudes (Ao) of 2-10 nm and imaged in tapping mode using silicon cantilevers (Nanosensors). The image amplitude (set point As) and A0 ratio (As/A0) was kept at ∼0.8 in a repulsive tip-sample interaction regime, and phase oscillations were no greater than ± 5 degrees. The surface was rastered following the fast scan axis (x) at rates of 2 Hz, capturing the retrace line to reconstruct the AFM micrographs. All samples were scanned at room temperature in air, at a relative humidity of 30%.

### Tetranucleosome Phase Transition Visualized by Epifluorescence Microscopy

For phase transition experiments using fluorescence microscopy, tetranucleosome stocks 100-4W-50 (300 nM) and Cy3-100-4W-50 (300 nM) were combined at a molar ratio of 3:1, respectively, in 10 mM HEPES-NaOH pH 7.5; 1 mM EDTA; 1 mM DTT. Nucleosome phase transition was induced by diluting the tetranucleosome stocks tenfold in 50 mM Tris-HCl pH 7.5; 150 mM NaCl; 5 mM MgCl2; 1 mM DTT. Phase transition reactions were incubated at room temperature, and 4.5 µL aliquots were combined at different times with 0.5 µL of 10X GODCAT visualization buffer (50 mM Tris-HCl pH 7.5; 2 mg/mL Glucose Oxidase (SIGMA cat. no. G2133); 350 ng/mL Catalase (SIGMA cat. no. C1345); 20% Glucose; 10 mg/mL AcBSA (Thermo); 50% glycerol) and added into the PEGylated and BSA passivated slide/coverslip chamber.

To prepare the visualization chamber, glass coverslips and slides were passivated using polyethylene glycol (PEG) as previously described (Chandradoss et al., 2014). A microfluidic chamber was assembled with both the PEG-coverslip and the PEG-slide, and the channel formed between both surfaces was blocked for 30 min by the addition of 1 mg/mL of AcBSA (Thermo) in 20 mM Tris-HCl pH 7.5; 50 mM NaCl, and 1 mM EDTA, followed by rinsing with 50 mM Tris-Cl pH 7.5; 150 mM NaCl; 5 mM MgCl2; 1 mM DTT. 5 µL of the phase transition reaction combined with the GODCAT was applied to the channel chamber and visualized using epifluorescence microscopy. Nucleosome phase transition in the presence of human H1.0 linker histone (The Histone Source, Colorado State University) was performed in the same way as explained above, but including H1 in the phase transition reaction at a concentration of 120 nM to keep a ratio of H1: nucleosome 1:1.

Epifluorescence micrographs were taken using a laboratory built wide-field fluorescence microscope. An oil-immersion 100x objective lens (Olympus 100× UPlansApo, N.A. 1.4) was used with a diode-pumped 532-nm laser (75 mW; CrystaLaser) for excitation and a red LED for brightfield illumination. Fluorescence emissions and brightfield illumination were splitted via a laboratory-built multichannel imaging system and projected side-by-side on an EMCCD (IXon EM+ 897; Andor). The illuminated area on each channel was ∼50 × 25 μm with ∼110 nm pixel widths.

### Tetranucleosome Phase Transition visualized by OpNS-EM

Tetranucleosome stocks 100-4W-50 and biotin-100-4W-50 were combined at a molar ratio of 3:1, respectively, in 20 mM HEPES-NaOH pH 7.5; 1 mM EDTA; 1 mM DTT. Nucleosome phase transition was induced by diluting the tetranucleosome stocks tenfold in buffers with different salt concentrations (20 mM HEPES-NaOH pH 7.5; 0-150 mM NaCl; 0-5 mM MgCl2; 1 mM DTT). Phase transition reactions were incubated at 4°C, 20°C, and 36°C, and were analyzed at different incubation times by the optimized negative staining protocol (OpNS) as previously described (Rames et al., 2014). In brief, 4 μL of the phase transition reaction was deposited on a glow-discharged ultra-thin carbon-coated 200-mesh copper grid (CF200-Cu-UL, Electron Microscopy Sciences, Hatfield, PA, USA, and Cu-200CN, Pacific Grid-Tech, San Francisco, CA, USA). After 30 seconds of incubation, phase transition solution on the grid was blotted with filter paper. The grid was stained for 10 s with 30 µL of 1% (w/v) uranyl formate (UF), UF was blotted with filter paper, and the staining procedure was repeated two more times, followed by air-drying with nitrogen. OpNS images were acquired under a defocus of 0.5-1 µm on a Zeiss Libra 120 Plus transmission electron microscope (TEM) (Carl Zeiss NTS GmbH) equipped with a Gatan UltraScan 4 k × 4 k CCD. The TEM was set to a high-tension of 120 kV with energy filtering at 20 eV. Carbon-coated grids are usually glow-discharged in residual air to promote the absorption of biological material deposition from the aqueous solution, a process that makes the surface hydrophilic by depositing negative charges. However, the negatively charged surface does not efficiently bind the negatively charged nucleosome arrays (Fig. S5; blue section). An analysis of the effect of Na^+^ and Mg^2+^ indicates that in absence of Na^+^, spherical droplets were formed and attached to the surface above 0.5 mM Mg^2+^, but irregular condensates were not observed (Fig. S5A), and when tetranucleosomes were incubated in absence of Mg^2+^, with increasing Na^+^ concentrations, tetranucleosome formed irregular condensates above 1.5 mM Na^+^, which became attached to the surface (Fig. S5B). When Mg^2+^ and Na^+^ are present at physiological concentrations the surface becomes correspondingly more positive and robustly binds the condensates (Fig. S5C).

### Preparation of Streptavidin 2D Crystal Grids and cryo-EM Sample Deposition

The streptavidin (SA) crystal grids were prepared as previously described (Han et al., 2016) with a few modifications. Lacey carbon grids with 5 µm holes (LC200-Au-FF, Electron Microscopy Sciences) and Quantifoil carbon grids with 7 µm holes (S 7/2, Electron Microscopy Sciences) were cleaned with 100% chloroform to remove the plastic cover. Since functionalization of carbon grids involves hydrophobic interactions, no glow discharge was applied on the grids. To form a biotinylated monolayer of lipids, 30 µL of castor oil (BRAND) was added to 5 ml of crystallization buffer (50 mM HEPES-NaOH pH 7.5; 150 mM KCl; 5 mM EDTA) in a 50 mm plastic petri dish, followed by the addition of 1 µL of 1 mg/mL Biotinyl Cap PE (1,2-dipalmitoyl-sn-glycero-3-phosphoethanolamine-N-(biotinyl); Avanti 870277)). To functionalize the grids, cleaned carbon grids were put in contact with the lipid monolayer for 1 s, then washed three times with 100 µL of crystallization buffer. To add the streptavidin surface to the grid, 4 µL of 0.2 mg/mL of streptavidin (NEB; N7021S) in crystallization buffer were added to the biotinylated lipid surface and incubated for 30 min in a humidity chamber at room temperature. To remove excess SA, SA-crystal grids were washed and blocked with 30 µl of rinsing buffer (10 mM HEPES-NaOH pH 7.5; 50 mM KCl; 5 mM EDTA; 10% trehalose), followed by blotting and air drying. A thin layer of carbon was deposited by evaporation onto the backside of the SA crystal grid to protect the crystal from being damaged during grid manipulation. SA-crystal grids were stored at room temperature in a sealed box for further use.

For sample deposition, excess trehalose was removed from SA crystal grids by rinsing the grids two times with 30 µL of crystallization buffer for 10 min. Grids were blotted at the edge with filter paper, then 4 μL of the tetranucleosome phase transition reactions (prepared as for OpNS-EM visualization) were added to the SA grids and incubated in a humidity chamber to allow for biotin and streptavidin binding. After 2 min, grids were blotted for 2 s and immediately flash frozen in liquid ethane at 100% humidity to form an amorphous ice, using a Leica EM GP rapid-plunging device (Leica, Buffalo Grove, IL, USA). Humidity chamber and the plunging device were set to the temperature used in each phase transition reaction.

### Cryo-ET Data Acquisition

We found that the negatively charged nucleosome arrays did not efficiently bind to the negatively charged carbon grids, and while the use of Mg^2+^ increased deposition (Fig. S5), it made it difficult to control the degree of condensate adsorption at different ionic strengths. For our cryo-EM studies and to avoid using a charged grid surface, we used the 2D streptavidin crystal method (Han et al., 2016) (Fig. 1A) and tetranucleosome arrays harboring a biotin molecule at one DNA end. We observed similar spherical condensates on grids prepared with and without the streptavidin crystal, after long incubation times and high salt conditions, indicating that the condensates form in solution and are then absorbed onto the SA-crystal (Fig. S6).

Frozen nucleosome samples on SA-crystal grids were subjected to tomography data collection on a FEI Titan Krios transmission electron microscope, operated at 300 kV with an energy filter at 20 eV (Gatan BioQuantum). Tilt series were acquired with SerialEM (Schorb et al., 2019) using the automated data collection function, including fine eccentricity, item realignment, and autofocus scheme. The range for tomography collection was set from −60° to +60°, with an angular increment of 3° per tilting step. The target defocus was set between 2.5-3 μm. The total electron dose for the whole tomography was calculated between 200 and 250 electrons per Å^2^, with an adjustment of the exposure time following the 1/*cos* schemes. For each tilt angle, 8-10 frames were acquired every 0.25 s from 2 to 2.5 s of exposure using a Titan Krios Gatan K2/K3 camera with a pixel size of 1.45/1.47 Å.

### Deep Learning-based Denoising

For deep learning denoising, we followed the description of the PROJECTION2PROJECTION method (Buchholz et al., 2019). Frames of tilt angles were aligned using MotionCor2 (Zheng et al., 2017) to correct for image drift. Aligned frames were split into even and odd frame stacks and averaged separately to generate noise independent pairs for the same object. To select representative areas of the averaged image pair, 1,120 local box path-pairs of 600×600 pixels were generated at each apparent nucleosome core particle (NCP) position. Inside each NCP position, 10 patches of 128 × 128 pixels were further randomly cropped to train a CARE network in the NOISE2NOISE regime (Weigert et al., 2018). The neural network was constructed with a U-Net of depth two, a kernel size of three, one hundred training epochs, and a linear activation function at the last layer. The loss function was evaluated with a per-pixel mean squared error (MSE). After training, the network was applied to restore the full-size image pairs at each tilt. The final image of each tilt is the result of averaging two individual restorations of even and odd images.

### Preprocessing of cryo-ET Tilt Series

The denoised tilt series were pre-aligned using the IMOD software package with the patch tracking method (Kremer et al., 1996). Then, further alignment was performed on the local image patches (200 x 200 pixels) by using the focused refinement strategy in individual-particle electron tomography (IPET)(Zhang and Ren, 2012). The alignment of the image tilt axis onto its central vertical axis allowed for the determination of the Contrast Transfer Function (CTF) of the microscope, using the GCTF software package (Zhang, 2016). CTF of the center slice of each micrograph and the cosine of the tilt axis angle were used to estimate the CTF of the flanking strip bands. Both the phase and amplitude were corrected for each strip band using TomoCTF (Fernandez et al., 2006).

### Cryo-ET 3D Reconstruction and NCPs Segmentation

To obtain the single-molecule 3D images, the CTF corrected tilt series were then submitted for a focused electron tomographic refinement (FETR) algorithm in IPET (Zhang and Ren, 2012) followed by 3D reconstruction using weighted Fourier back projection using the e2tomo software package (Chen et al., 2019; Tang et al., 2007) without a further alignment step. The reconstructed maps were subjected to low pass (abs= 0.25), normalization and median shrink filters enhancement (bin*=* 8; pixel size= 11.68 Å). To implement the deep learning-based segmentation method in e2tomo, areas containing representative features for network training (crystal lattice, free distal DNA, NCPs, and background noise) were boxed out from z-dimensional slices of the reconstruction. The application of the trained network to the tomograms allowed for the segmentation of the NCPs, DNA, and SA crystals from the background. NCP segmentation was used as a marker to select surrounding connected map densities (within 10 Å) using the zonsel function of Chimera software (Pettersen et al., 2004). SA crystal segmentation was used as a marker to remove the surrounding SA crystal densities (within 10 Å). To produce a soft boundary mask for the missing wedge correction, the processed density maps were low-pass filtered to 30 Å and clipped to box of 512 x 512 x 200 pixels.

### Missing Wedge Correction of 3D Density Map Reconstruction

To reduce the tilt limitation caused the elongation artifact in the 3D reconstruction, the missing wedge correction was performed by the low-tilt tomographic reconstruction (LoTToR) method (Zhai et al., 2020), which is a model-free iterative process, under a set of constraints in the real and reciprocal spaces, aiming to fill the missing wedge zone in Fourier space. To implement the algorithm, a soft boundary 3D mask was generated using the previously denoised density map after 30 Å low-pass filtering. This 3D mask was paired with the initial 3D reconstruction to generate nine cubic tiles of 200 x 200 x 200 pixels with overlapping regions of 30 x 200 x 200 pixels as reference for realignment. These cubic tiles were subjected to iterations of the missing wedge correction until the final corrected 3D map converged with a stable Fourier shell correlation value. The final 3D density map of 512 x 512 x 200 pixels was reassembled by aligning the corresponding missing wedge corrected density cubes.

### Modelling of the final denoised 3D Electron Density Map

A 3D global template-matching was obtained by competitively docking the nucleosome model (PDB ID:1AOI) and a linear 40 bp DNA model into the final missing wedge corrected density map. This procedure consisted of four major steps: i) NCP segmentation maps were split into domains using the watershed method of Chimera (4 smoothing steps with a step size of 3 voxels). High-density weight centers of each domain were calculated at different map contour levels (from 10-90% of its maximum in a step of 0.5). ii) NCP and 40 bp DNA models were fitted into the weight centers with a random shift and rotation. iii) The competitive docking of positioned NCP and DNA models was determined using the sequential multi-model fitting function in Chimera (Pettersen et al., 2004). All docked models were ranked based on the cross-correlation value (CC-score) between the models and the map. Models with either a CC-score < 0.7 or exhibiting a clash event with higher-ranked models were removed from the analysis. iv) Steps ii) and iii) were repeated multiple times until a stable fitting was identified as explained below.

A stable fitting of each map domain was obtained by the process of maximizing the average CC-score and minimizing the map/model fitting residue (MaFR/MoFR). MaFR corresponds to the remaining density volume after subtracting the combined model from the density map, and MoFR corresponds to the remaining density volume after subtracting the density map from the combined model. A score to evaluate the minimization of the fitting residue compared the current fitting with the best previous fitting and was defined as the ratio between (MaFRcurrent - MaFRbest)/MaFRbest and (MoFRcurrent - MoFRbest)/MoFRbest. A negative value of both (MaFRcurrent - MaFRprevious) and (MoFRcurrent - MoFRprevious) indicated that a better fitting was found. A score threshold of < −0.1 was set to accept the current fitting as the best fitting. The fitting loop was stopped when no better fittings were found in ten consecutives iterations. The translations and rotations of each NCP and DNA linker models were recorded for the statistical analysis described in the following sections.

### Identification of Free and Condensed NCPs

The weight centers of NCPs in the fitted model were classified based on their distance from their neighbors, using the density-based spatial clustering of applications with noise algorithm (DBSCAN) (Ester et al., 1996). DBSCAN required two input paramaters, *eps* and *min_samples*, where *eps* define the maximum distance between two points to consider them as neighbors, and *min_samples* define the minimum number of neighbors to consider them as clusters. An initial classification to identify free NCPs and condensates was performed using an *eps* value of 250 Å, which is the minimal theoretical distance between two consecutive nucleosomes in the same array (2 x 55 Å NCP (disc radius) + 40 bp DNA x 3.4 Å/bp= 250 Å), and a *min_sample* value of 8, which is, by definition, the smallest cluster composed of two tetranucleosome units. A second classification was performed based on the previous classification, using an *eps* value between 270-290 Å, and a *min_sample* value of 12. This produced a cleaner segmentation that differentiated free NCPs from NCPs grouped in condensates, representing the light and dense phase of spinodal decomposition, respectively. In the case of small spherical condensates that interact among themselves or with irregular condensates, a subclassification was manually performed to distinguish each spherical condensate within the cluster.

### Calculation of Tetranucleosome Condensates Geometry

The shape of irregular and spherical nucleosome condensates was determined by measuring its physical dimensions and eccentricity. The condensate dimensions are represented by the length of its long and short axis. The direction of the long and short axis was calculated using the coordinates of the center of mass of NCPs and the principal component analysis (PCA)(Pearson, 1901), which produced the vector PC1 for the long axis and the vector PC3 for the short axis. The lengths of the long and short axis were calculated as the distances between the two most separated NCPs along the PC1 and PC3 vectors. The overall size of the condensate was estimated by averaging the length of the long and short axis. The eccentricity was calculated using the formula “1-(short axis length / long axis length)”, which yield a score between zero and one, where zero represents a perfect spherical shape, and one represents an elongated structure.

### Calculation of NCP Condensate Boundary Sharpness

To define the overall shape of condensates, a boundary mask was produced using the electron density of condensates with a 10 nm low-pass filter. To calculate the surrounding free NCPs concentration, which represents the condensate boundary sharpness, a shell of 20 nm extending from the boundary mask was established to determine the number of the surrounding free NCPs, which in turn was divided by the volume of the external 20 nm shell.

### Determination of NCP Concentration in Condensates

The nucleosome ice-slab (global) concentration for the early stage of phase separation (when light phase and dense phase are formed under ∼100 nm scale) was calculated by dividing the total number of NCP within the tomogram by the volume of the smallest cuboid that enclosed all NCPs. To measure the concentration of NCPs in spinodal and spherical condensates, an initial condensate-shaped mask was produced based on the electron density after a 10 nm low-pass filter, and a series of 3D contours were established either by shrinking or expanding the mask in steps of 2 nm. The innermost contour was labeled as *n=* 1, and the next consecutive contour was labeled as *n +* 1. A shell defined as the 3D space between two consecutive contours was also labeled starting from inside the condensate (*n=* 1). To calculate the concentration of NCPs inside a specific contour *c*, the number of NCPs within contour *n* was divided by the volume enclosed by the contour. To calculate the concentration of NCPs inside a specific shell *n*, the number of NCPs within the shell *n* was divided by the volume of the shell. To avoid double counting NCPs that cross two shells or contours, the center of mass of NCP was used to determine its residing shell or contour. The mean NCP concentration was obtained by averaging the NCP concentration inside a specific contour or shell *n* at the same position in the same type of condensates. Since condensates vary in size, the total number of contours within a condensate is different from each other, which may cause more variance in concentration measurements for the exterior contour/shell.

### Calculation of the Pair Distribution Function (*g(r)*) of NCPs Within Condensates

The *g(r)* function was determined using the distribution of the pairwise distances between the weight centers of NCPs. For each NCP weight center, a series of spheres separated by 1.6 nm (*dr*; small step of radius extension) were generated using a radius (*r*) ranging from 0 to 60 nm. This resulted in shells formed by two consecutive concentric spheres (i.e., the shell between *r* and *r* + *dr* away from the reference NCP weight center). The NCP density of each shell was calculated by dividing the number of NCPs within the shell by the shell volume (*4πr^2^ x dr*), and the shell densities of different spheres with the same radius were averaged and divided by the ice-slab (global) density for normalization. However, the shell density estimation for NCPs that reside at the border of the tomography is not accurate. To correct this, a periodical boundary was applied to the *x-* and *y*-direction of NCP models, lengthening the respective dimensions of the tomogram 2-fold. If the shell surpasses the *z*-direction boundary of the tomogram, a correction factor (the ratio between the overall shell volume and the partial shell volume inside the tomogram) was multiplied with the corresponding shell density because the *z*-direction is non-extensible, due to contact with ice (top) and the SA-crystal surface (bottom). For tomograms containing spherical condensates, only the condensate region was selected for *g(r)* analysis, and due to the large background density variation among different condensates, the *g(r)* density was normalized by setting the highest peak equal to one.

### Calculation of the NCPs Orientation Within Condensates

To calculate the orientation of NCPs on the surface and in the interior of different types of condensates, each NCP within the fitted model was simplified as a triangular shape which centered on the nucleosomal discoidal plane. For spherical condensates, the orientation of each NCP relative to its containing condensates was measured as θ which is the angle between the triangular plane and the radius of the condensate that crosses the triangle center (i.e., the center of mass of NCP). Similarly, for irregular shape condensate that cannot define the radius, θ was measured as angle between the triangular plane and the line that is perpendicular to the spine of the condensate and crossing the triangle center. To calculate the spine of the condensate, the density map of the condensate was eroded to irregular lines by using Lee’s 3D skeletonizing algorithm (Lee et al., 1994). The θ angle distribution of NCPs for each type of condensate was evaluated by comparing with a *cos*(θ) function, which represent a randomly distributed orientation of NCPs. The *cos*(θ) represents the distribution of orientations of random unit vector relative to a fixed plane, which in turn equivalent to the probability of finding a point on the unit hemisphere contained in a differential ring-shape area: *P*(θ)*d*θ =2π(*Rcos*(θ))*Rd*θ/2π*R*^2^ = *cos*(θ)*d*θ.

## Acknowledgments

We thank Professor Geeta Narlikar for providing the expression vectors of *X. laevis* histones, and Dr. Hataichanok “Mam” Scherman (The Histone Source) for providing histone H1. We thank Professor David Limmer and Alex Tong for helpful discussions. Data was collected at the Cal-Cryo facility, Berkeley QB3 Institute. This research was supported by the Nanomachine program (KC1203) funded by the Office of Basic Energy Sciences of the U.S. Department of Energy (DOE) contract no. DE-AC02-05CH11231 and by contract no. DE-AC02-05CH11231 (C.B.). The work was also partially supported by grants from the US National Institutes of Health (R01HL115153, R01GM104427, R01MH077303, and R01DK042667 to G.R.; R35GM127018 to E.N; and R01GM032543 to C.B). C.B and E.N. are Howard Hughes Medical Institute Investigators.

## Author contributions

C.B., M.Z., C.D.C. conceived the study, and C.B., M.Z., G.R., and E.N. designed the research. C.D.C. and M.V purified and labeled histones, reconstituted and purified tetranucleosome samples. C.C.C. and C.D.C. performed the fluorescence microscopy imaging and B.O. collected the AFM images. M.Z. prepared EM specimens. M.Z. and J.L. collected the EM data. M.Z. and G.R. designed the cryo-ET work flow. M.Z. performed image processing and 3D reconstruction, M.Z. conducted the structural fitting, modelling and statistical analyses. M.Z. interpreted the data. M.Z prepared the EM related figures and videos. M.Z. and C.B. wrote the original draft and C.D.C., E.N., G.R., B.O., K.I.R., C.C.C., M.V. edited the manuscript. K.I.R. supplemented the theory section. C.B., and G.R., supervised the work.

## Competing Interests

The authors have declared that no competing interests exist.

## Supplemental Figures

**Figure S1.**
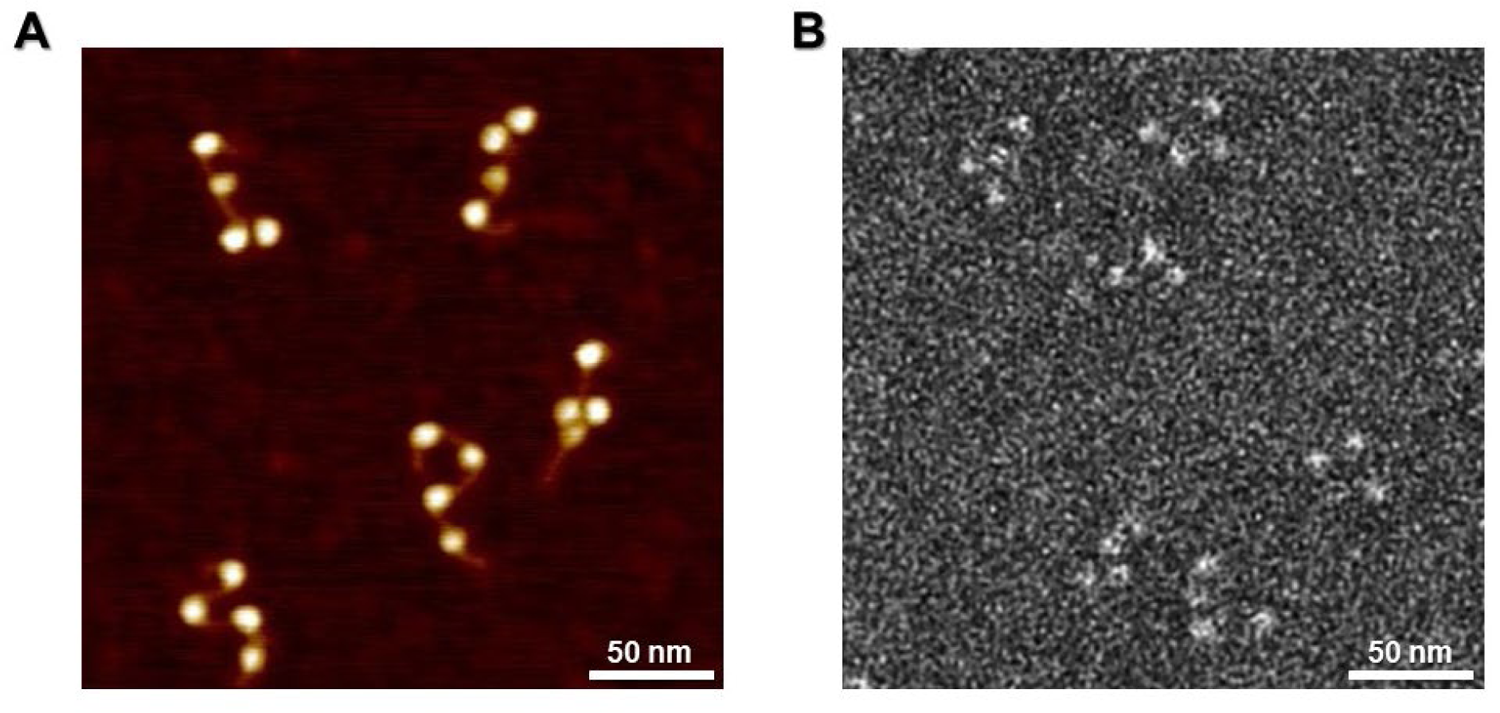
Tetranucleosomes reconstituted using recombinant *Xenopus laevis* histone octamers. Samples imaged by (A) Atomic Force Microscopy (AFM); and (B) Optimized negative staining (OpNS).

**Figure S2.**
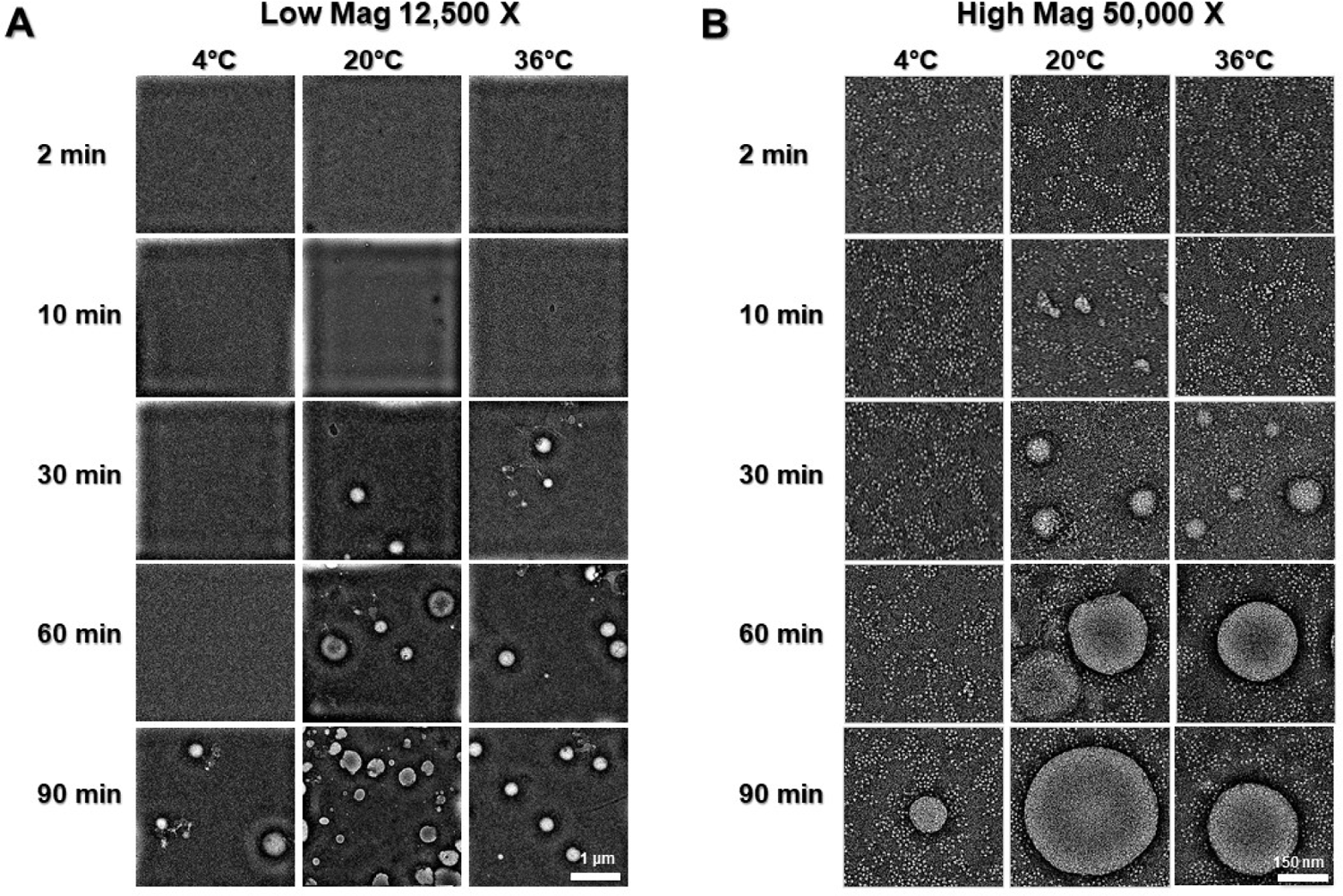
Tetranucleosomes phase transition visualized by OpNS. Representative time series of 30 nM tetranucleosome in physiological salt (20 mM HEPES-KOH pH 7.5; 150 mM NaCl; 5 mM MgCl2; 1 mM DTT) at 4°C, 20°C, and 36°C imaged at magnifications of (A) 12,500 X; and (B) 50,000 X.

**Figure S3.**
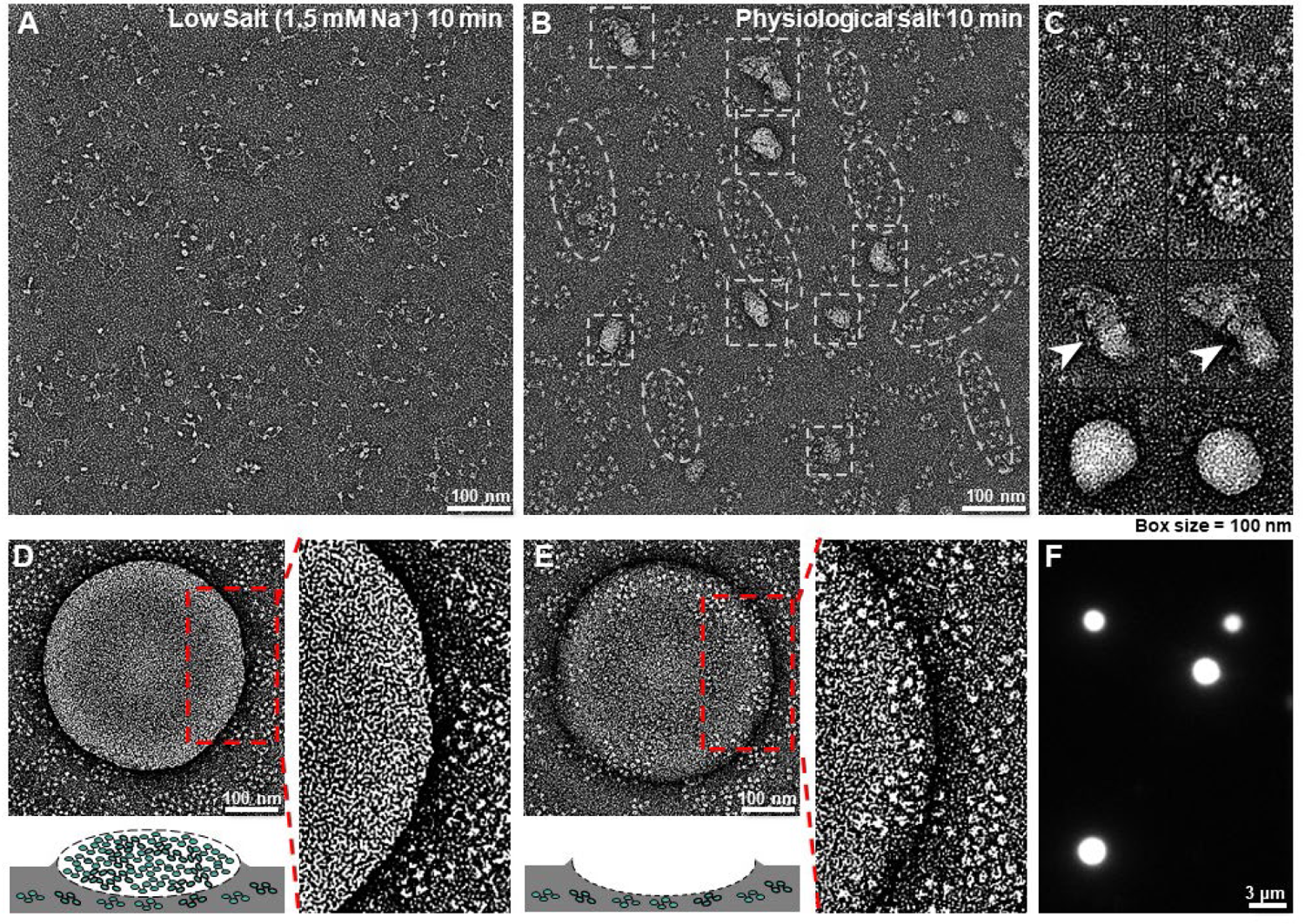
OpNS and fluorescence imaging of tetranucleosome phase transition. (A) Tetranucleosomes visualized by OpNS in low salt buffer (1.5 mM Na+) after 10 min incubation at 20°C. (B) Tetranucleosome condensates formed in physiological salt buffer after 10 min incubation at 20°C. Dashed-line ellipsoids enclose condensates displaying an irregular shape and loosely packed nucleosomes. Dashed-line boxes mark globular condensates displaying tighter nucleosomal packaging. (C) Representative images of nucleosome clusters in physiological salt showing different degrees of condensation, from loosely packed structures to tightly packed condensates (top to bottom). (D) OpNS image of a flattened spherical condensate in physiological salt (left top panel). The zoom-in view of the spherical condensate’s edge (right panel), corresponding to the rectangular dash line box, reveals tightly packed nucleosomes inside the condensate, surrounded by more loosely packed irregular condensates. The high density of the spherical condensate prevents visualization of the irregular condensates below it, as depicted schematically in the left bottom panel. (E) OpNS image of a spherical condensate landing area (left top panel). The condensate did not attach to the carbon surface during the staining process, leaving a dent on the staining material behind it upon drying, as shown schematically in the left bottom panel, and confirmed by the visualization of irregular condensates below its original landing place (right panel). (F) Fluorescence microscopy imaging of Cy3-labeled nucleosome liquid droplets formed in physiological salt after 30 min incubation at 20°C.

**Figure S4.**
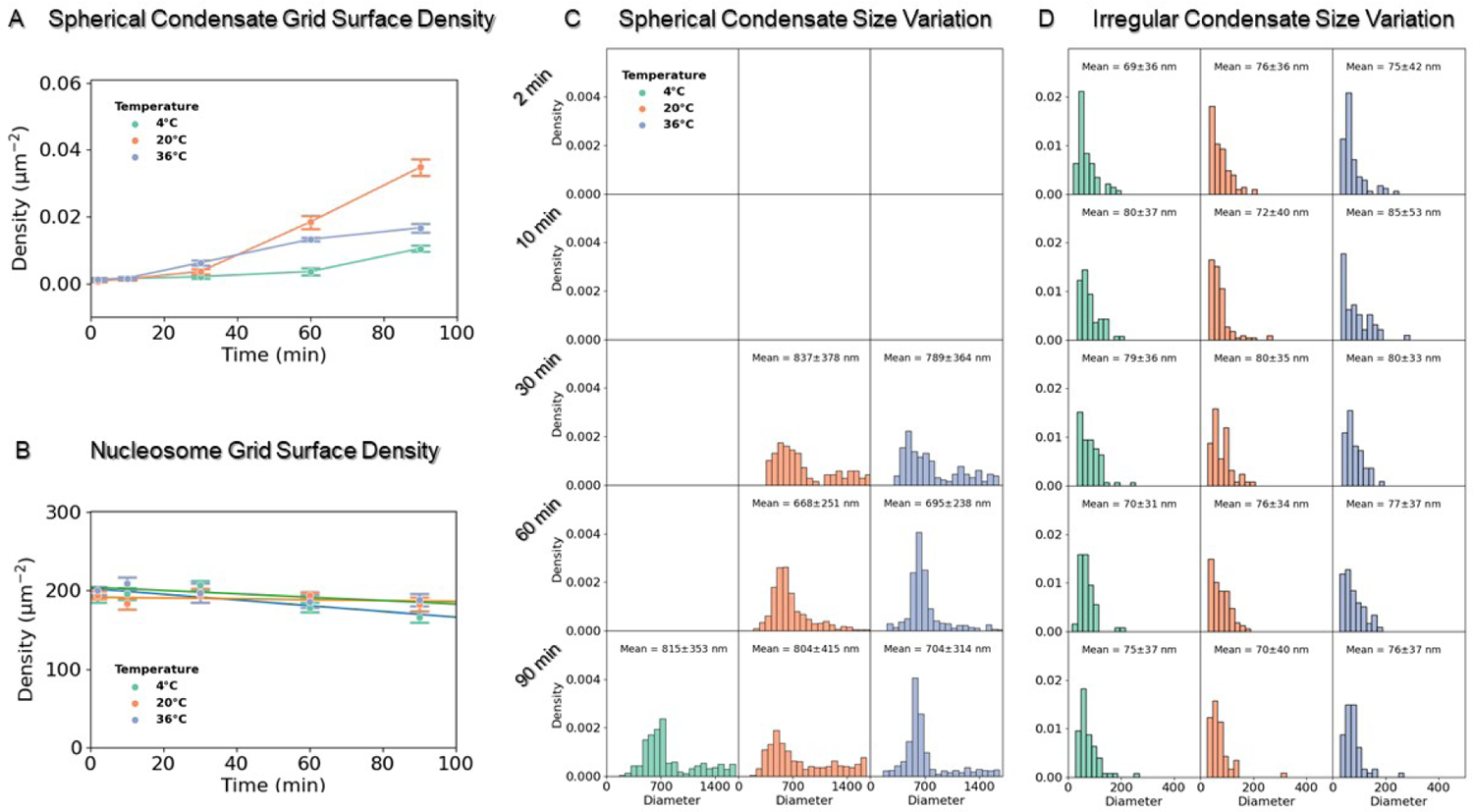
Quantitive measurement of tetranucleosome and spherical condensates grid surface density and condensates size. (A) Statistics of spherical condensate number density on carbon grid surface (number of condensates divided by the corresponding imaged area) obtained at various incubation time from OpNS samples (Fig. S2A). The incubation time series were conducted at 4°C (green), 20°C (orange), and 36°C (blue). (B) Statistics of nucleosome number density (number of nucleosomes within spinodal condensate divided by the corresponding imaging area) measured from the same time series samples at higher magnification (Fig. S2B). Data for (A) and (B) was obtained from 10-12 images and are presented as mean ± SEM. (C and D) Size (diameter) distribution of spherical condensates and irregular condensates measured from (A) and (B), respectively.

**Figure S5.**
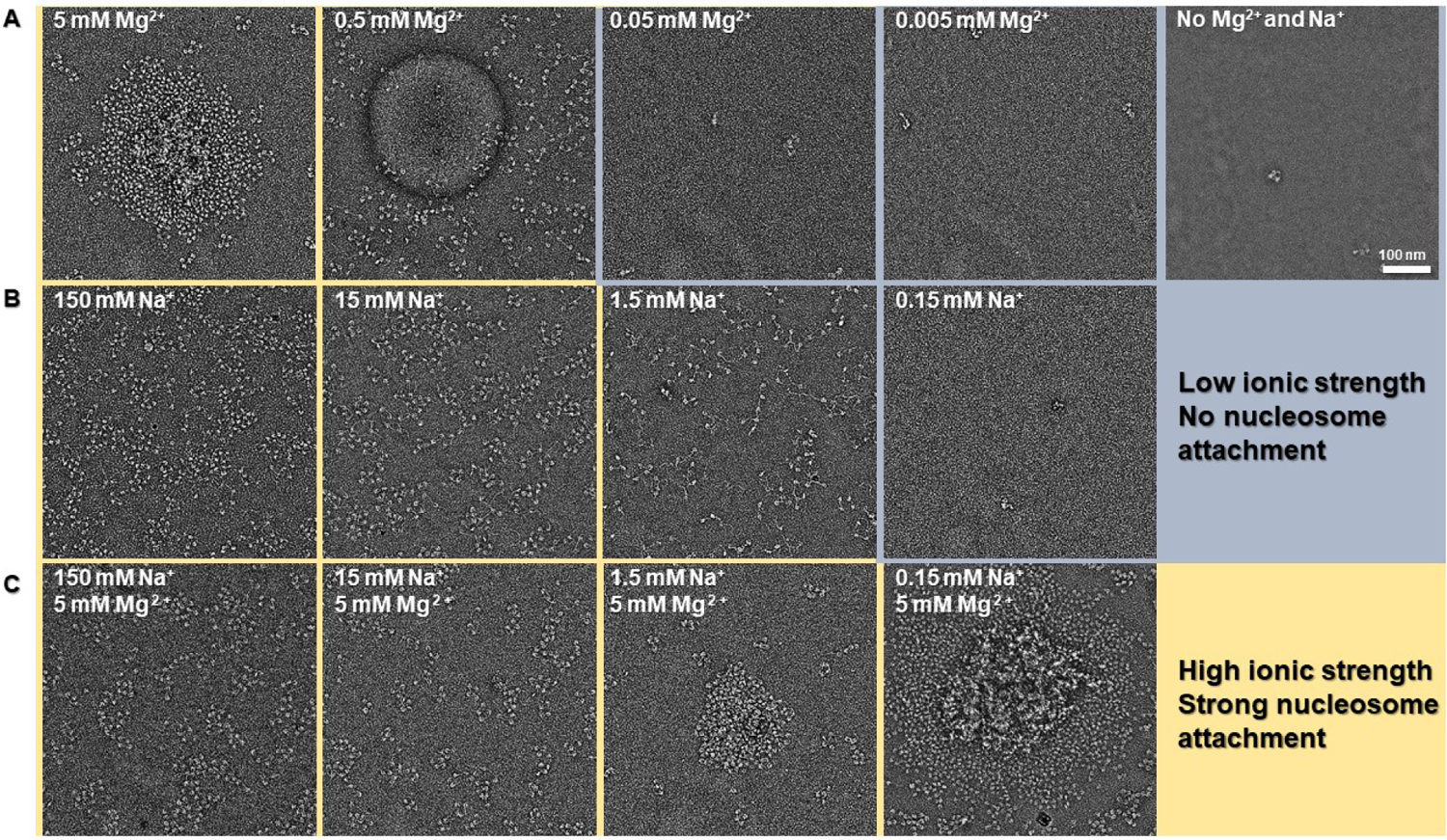
Effect of ionic strength on tetranucleosome conformation, condensate formation, and condensates attachment to a glow discharged carbon surface. In all conditions, tetranucleosome samples (30 nM) were incubated for 10 min at 20°C before OpNS. (A) Effect of Mg^2+^ concentration on condensate formation (20 mM HEPES-KOH pH 7.5; 5-0.005 mM MgCl2; 1 mM DTT). Spherical condensates are observed when Mg^2+^ concentration is above 0.5 mM. 5 mM Mg^2+^ causes the collapse of spherical condensates onto the grid surface, reflecting their high affinity to the carbon surface. Neither tetranucleosomes nor their condensates were observed on the surface when Mg^2+^ concentration falls below 0.5 mM (blue section). Interestingly, no irregular-shaped nucleosome condensates were observed with only Mg^2+^. (B) Effect of Na^+^ concentration on condensate formation (20 mM HEPES-KOH pH 7.5; 150-0.15 mM NaCl; 1 mM EDTA; 1 mM DTT). Irregular condensates are observed for Na^+^ concentrations above 15 mM. Tetranucleosomes do not attach to the grid for Na^+^ concentrations below 1.5 mM (blue section). Tetranucleosomes convert from an extended conformation at 1.5 mM Na^+^ to a relatively more compact conformation at 15 mM Na^+^. (C) Effect of Na^+^ concentration on condensate formation at constant Mg^2+^ concentration (20 mM HEPES-KOH pH 7.5; 150-0.15 mM NaCl; 5 mM MgCl2; 1 mM DTT).

**Figure S6.**
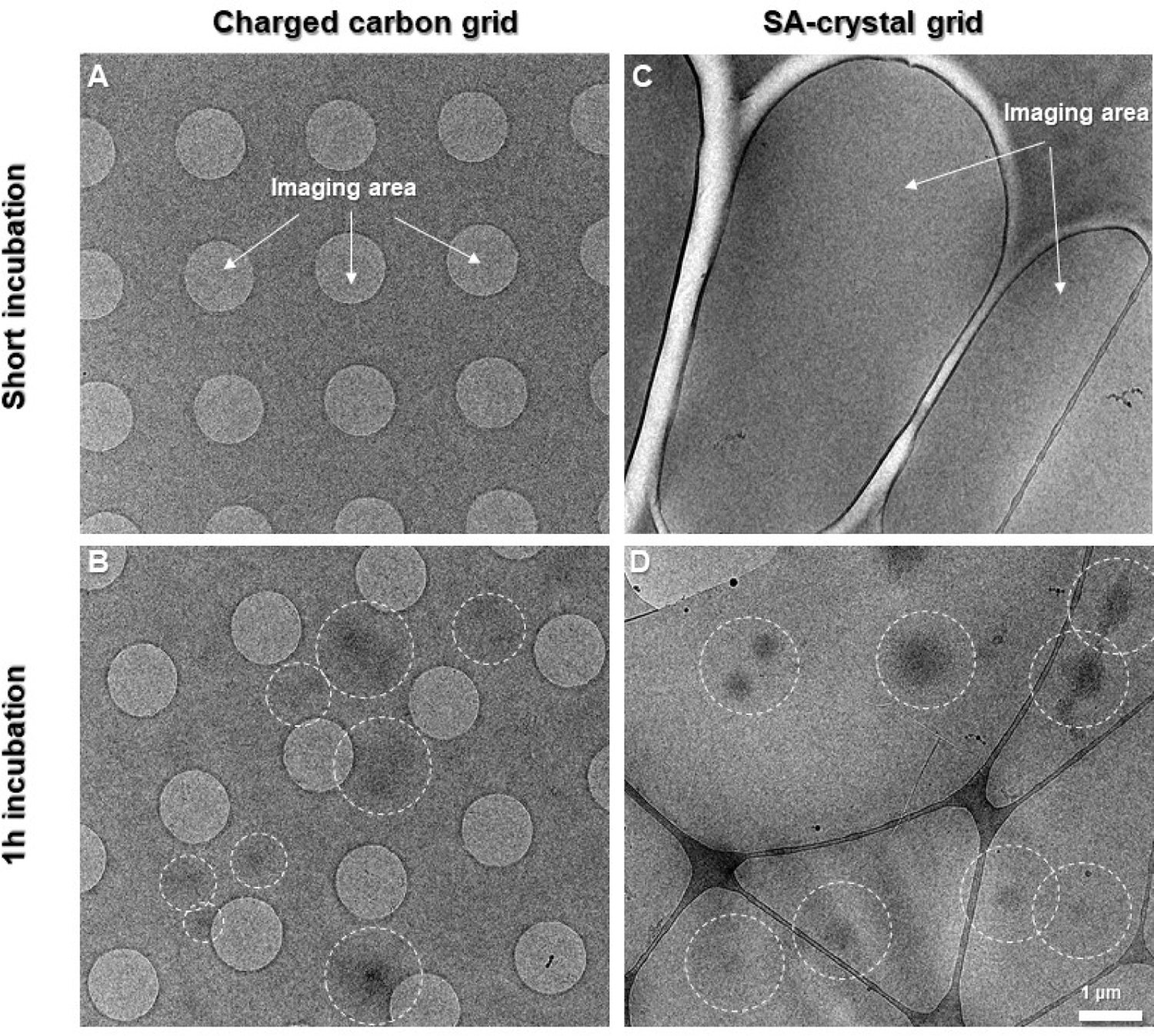
Streptavidin 2D crystal grid favors the binding of tetranucleosome condensates within the vitreous ice area. 30 nM tetranucleosome samples were incubated at 20°C in physiological salt (20 mM HEPES-KOH pH 7.5; 150-0.15 mM NaCl; 5 mM MgCl2; 1 mM DTT). (A and B) cryo-EM sample of tetranucleosomes prepared using a conventional glow discharged carbon quantifoil grid after short incubation (2 min) and long incubation (1 h). The circles are the areas of vitreous ice. Dashed line circles indicate the deposition of large-scale condensates onto the carbon area, away from the vitreous ice. (C and D) cryo-EM sample of tetranucleosomes prepared using a streptavidin 2D crystal substrate on a lacey carbon grid after short incubation (2 min) and long incubation (1 h). Similar size condensates are observed in the vitreous ice areas after long incubation (dashed circles).

**Figure S7.**
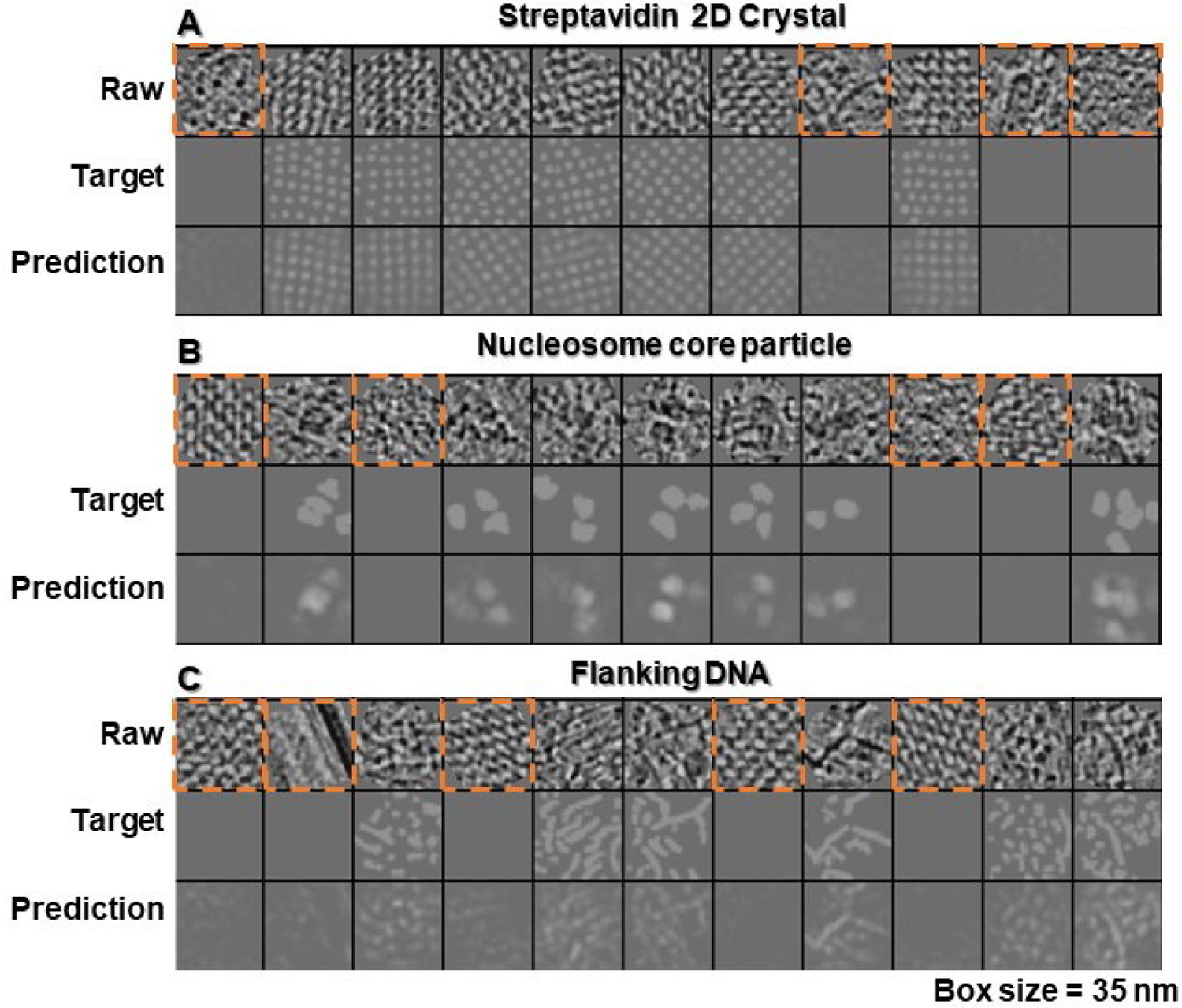
Deep learning-based segmentation of the initial 3D reconstruction. Network training and prediction for the (A) streptavidin crystal; (B) nucleosome core particle; and (C) flanking DNA component within tomograms. (A through C) First row: representative z-dimensional slice raw images containing the feature of interest. Representative negative examples containing features that do not belong to the category were also selected (orange dashed-line box). Second row: annotation of the first-row images by labeling out target features. Negative example images were targeted to a blank image. The raw and target images pairs were used to train the network. Third row: network prediction of the corresponding features after applying the trained network. The similarity between the target and prediction reflects the quality of the constructed network.

**Figure S8.**
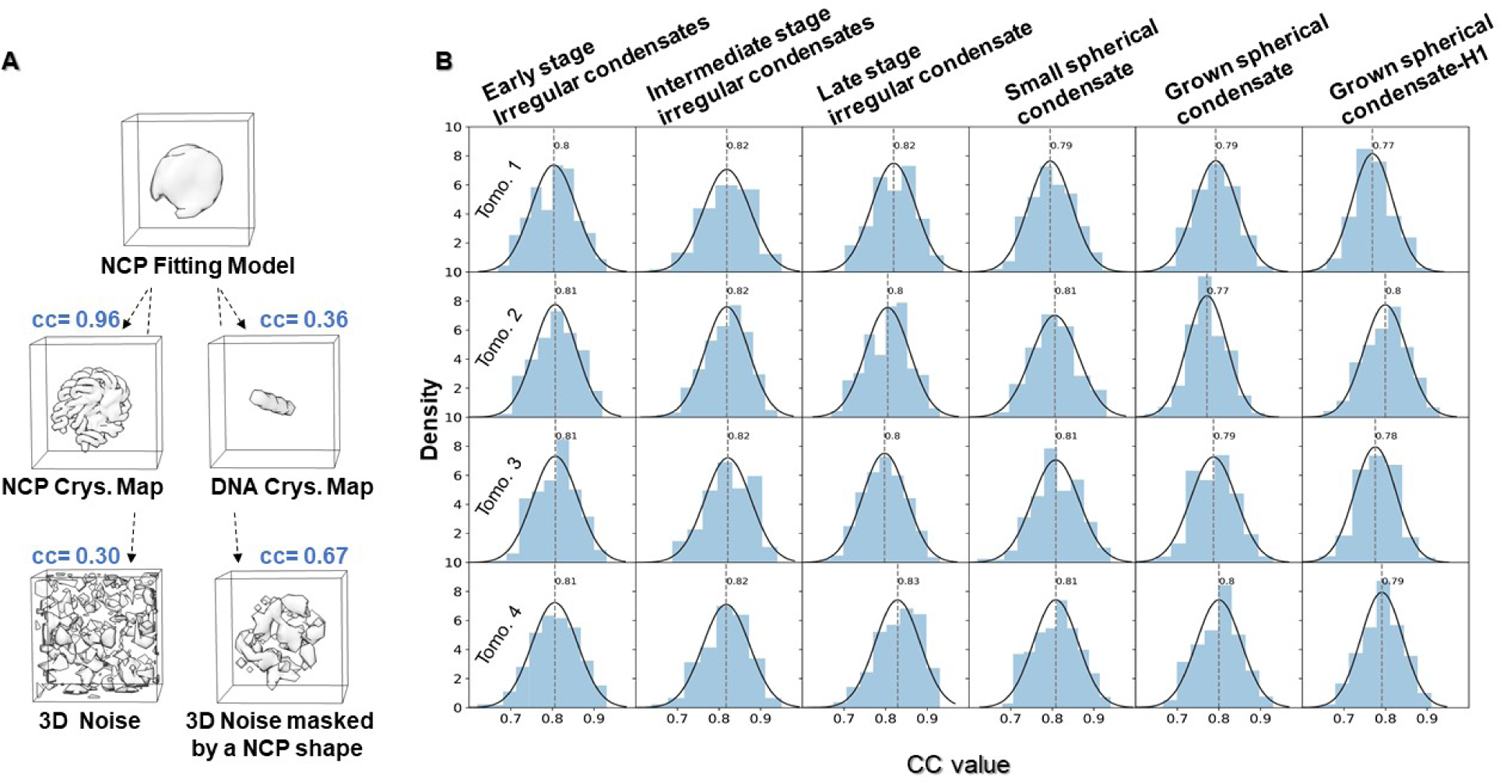
Distribution of map-model fitting scores. (A) Evaluation of cross-correlation coefficient between NCP fitting model with reference models, which yields a score of 0.96, 0.36, 0.30, and 0.67 for the NCP crystal structure map, DNA crystal structure map (40 bp), random noise map, and random noise enclosed by an NCP-shaped map, respectively. (B) Histograms showing the map-model fitting score distributions of all identified NCPs for four representative tomograms (rows). Each tomogram is analyzed in terms of early, intermediate, and late stage spinodal condensates (column one to three, respectively) as well as in terms of small spherical nuclei, grown spherical condensates, and grown spherical condensates with H1 (columns four to six, respectively). The fitting scores were calculated by measuring the cross-correlation between the local density map and the corresponding model.

**Figure S9.**
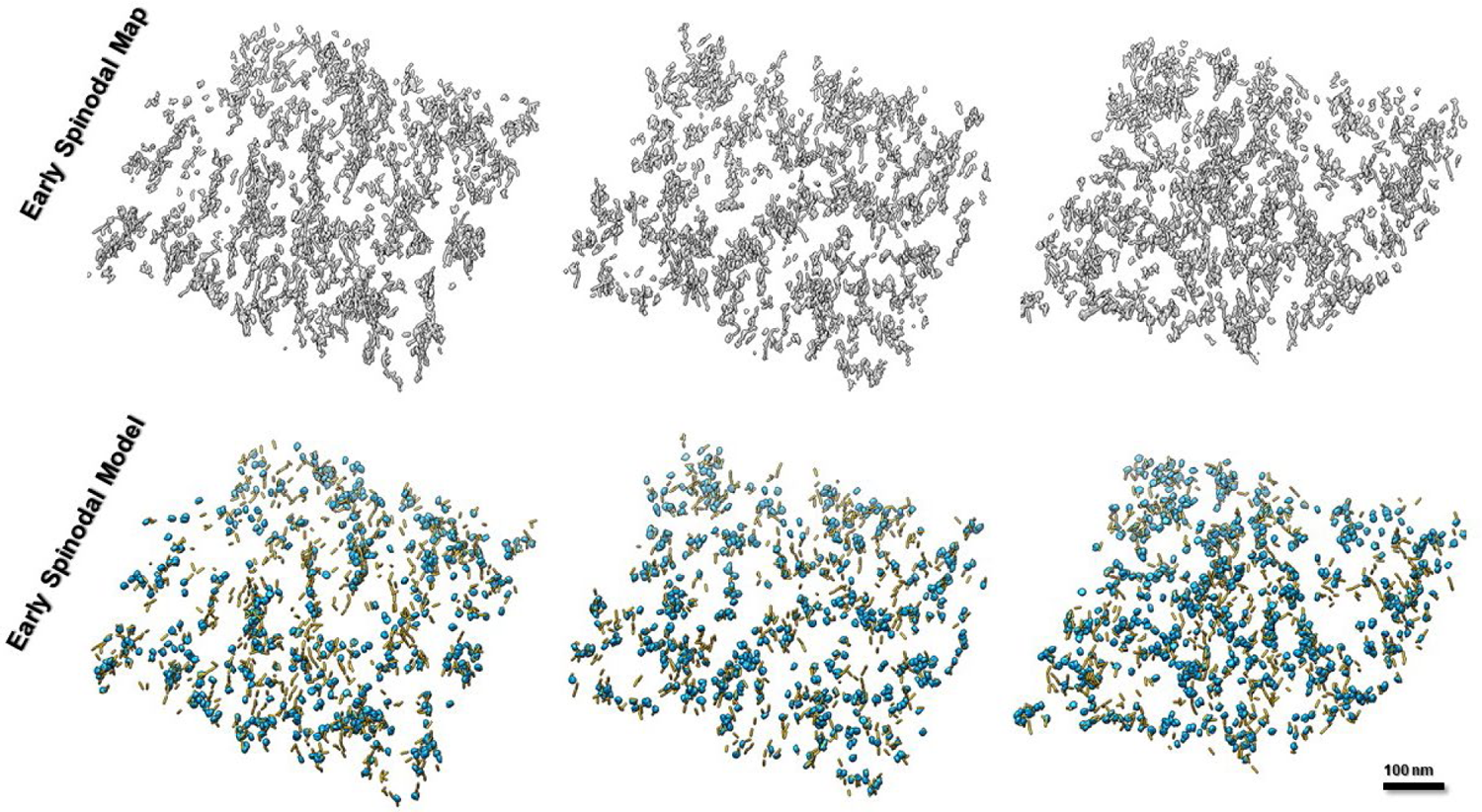
Fitted models of early irregular condensates. Comparison of three representative final density maps (top panels) with the corresponding fitted models (bottom panels) for early irregular condensates. Cyan indicates NCPs, and yellow indicates flanking DNA.

**Figure S10.**
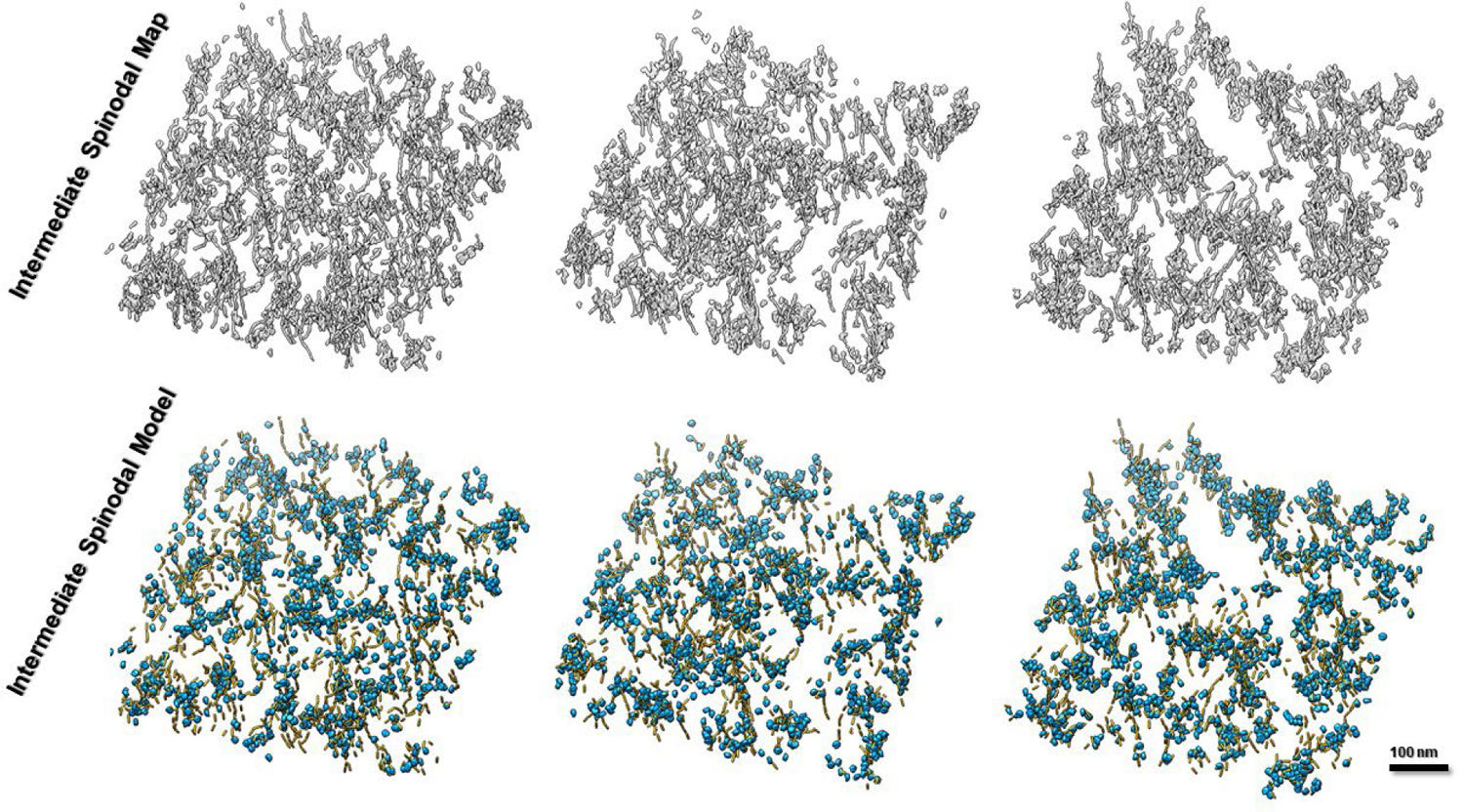
Fitted models of intermediate irregular condensates. Comparison of three representative final density maps (top panels) with the corresponding fitted model (bottom panels) for intermediate irregular condensates. Cyan indicates NCPs, and yellow indicates flanking DNA.

**Figure S11.**
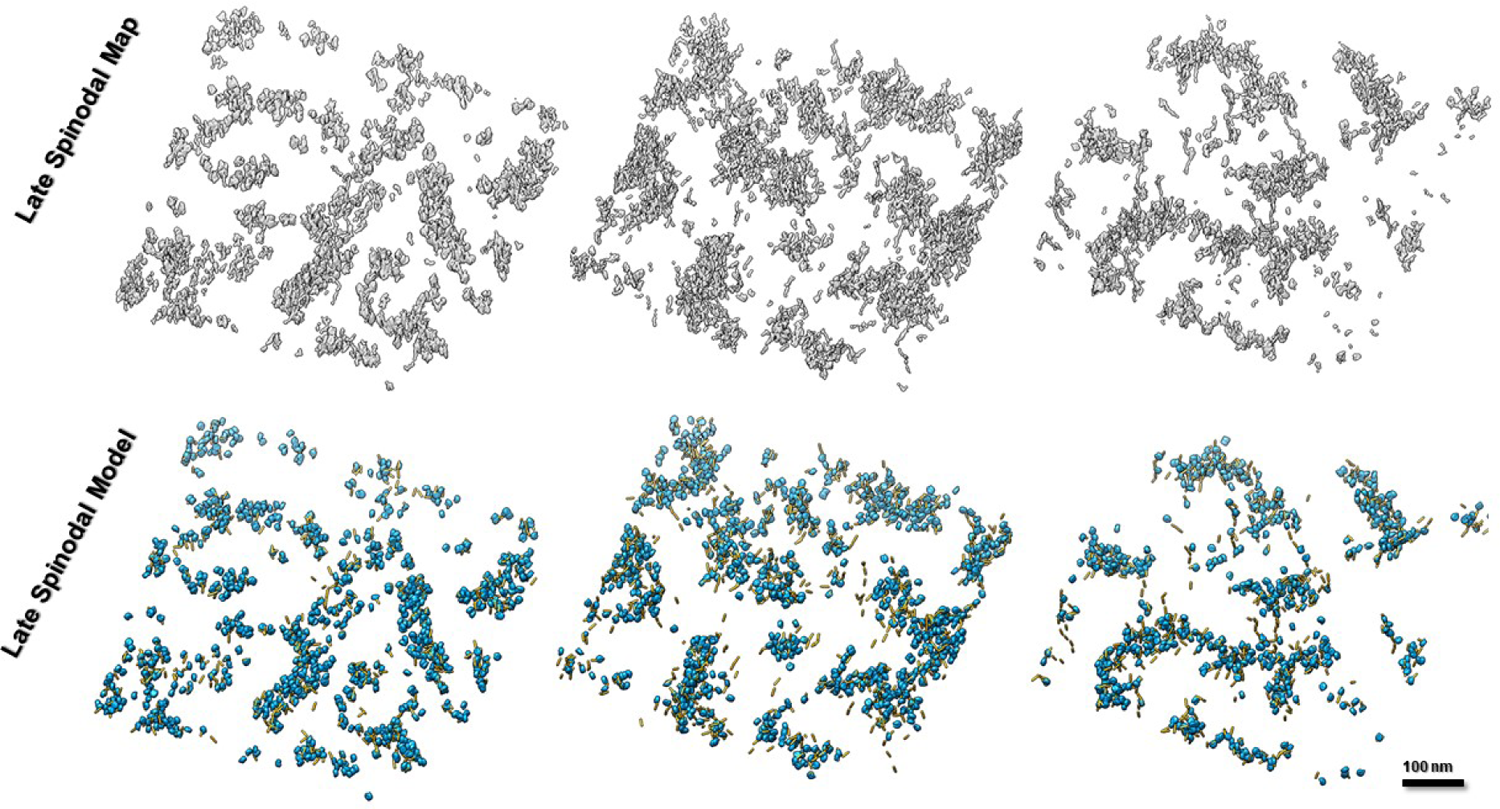
Fitted models of late irregular condensates. Comparison of three representative final density maps (top panels) with the corresponding fitted models (bottom panels) for late irregular condensates. Cyan indicates NCPs, and yellow indicates flanking DNA.

**Figure S12.**
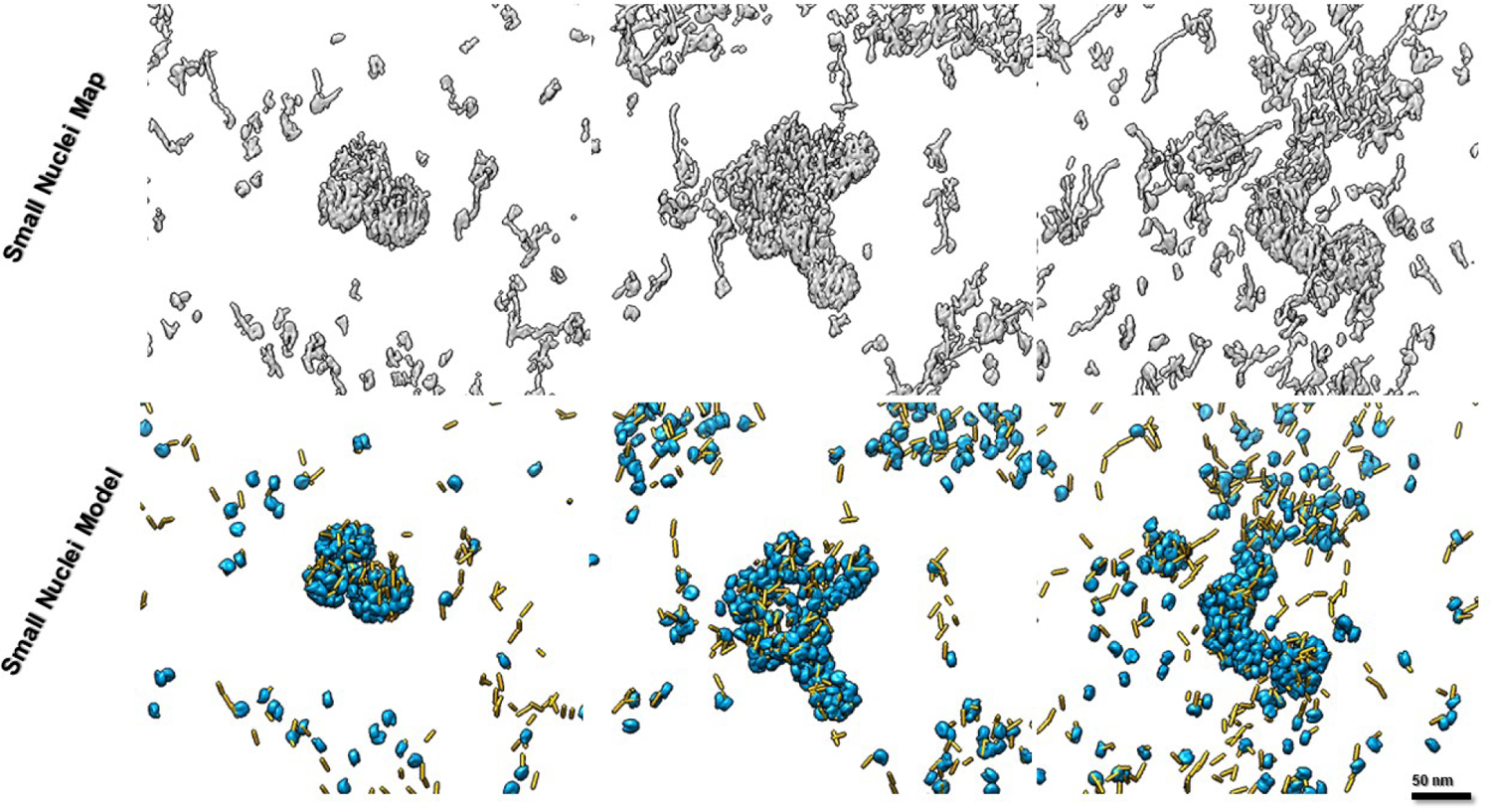
Fitted model of small spherical condensates. Comparison of three representative final density maps (top panels) with the corresponding fitted models (bottom panels) for small spherical condensates (nuclei). Cyan indicates NCPs, and yellow indicates flanking DNA.

**Figure S13.**
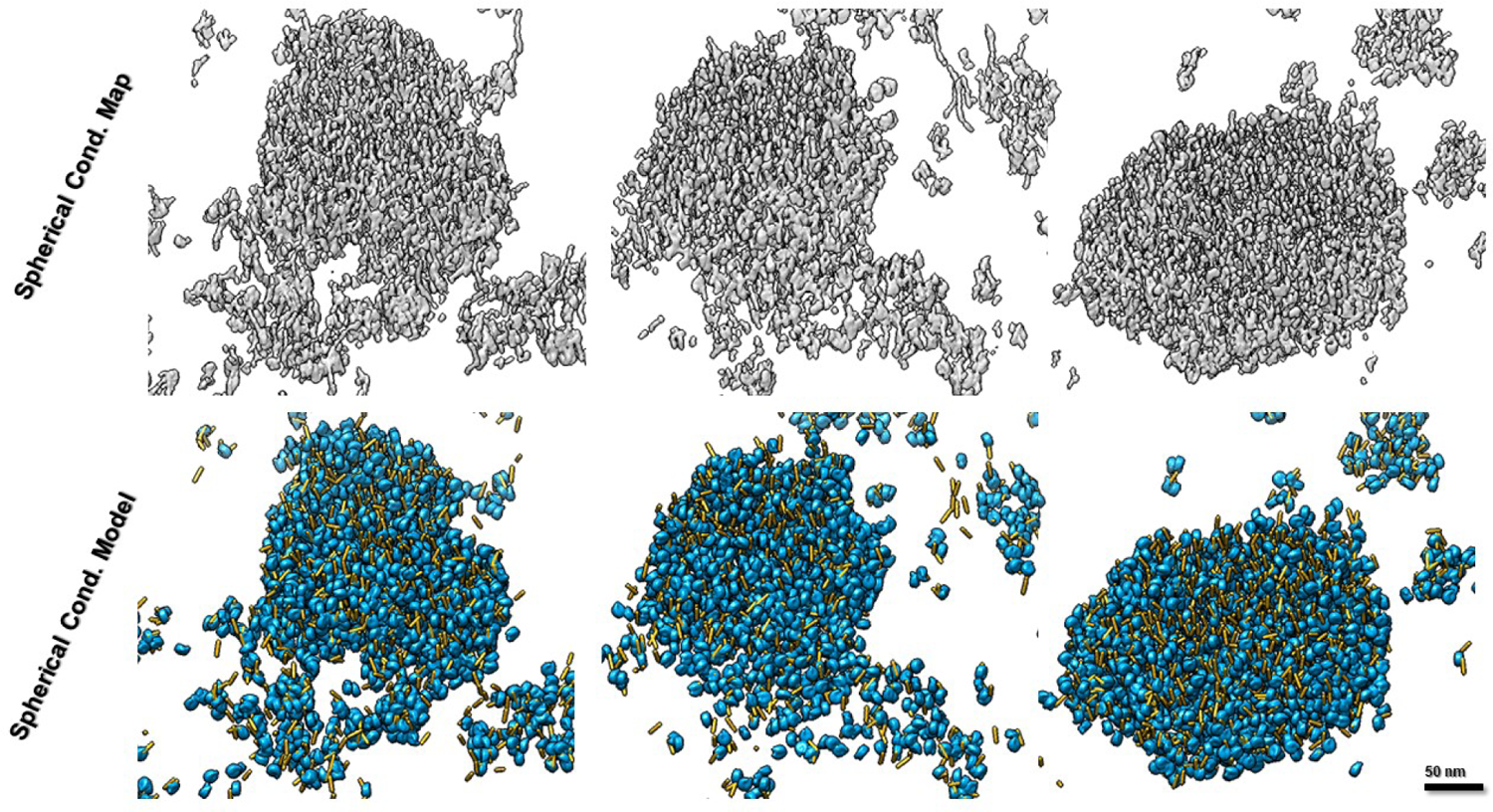
Fitted model of spherical condensates. Comparison of three representative final density maps (top panels) with the corresponding fitted models (bottom panels) for spherical condensates. Cyan indicates NCPs, and yellow indicates flanking DNA.

**Figure S14.**
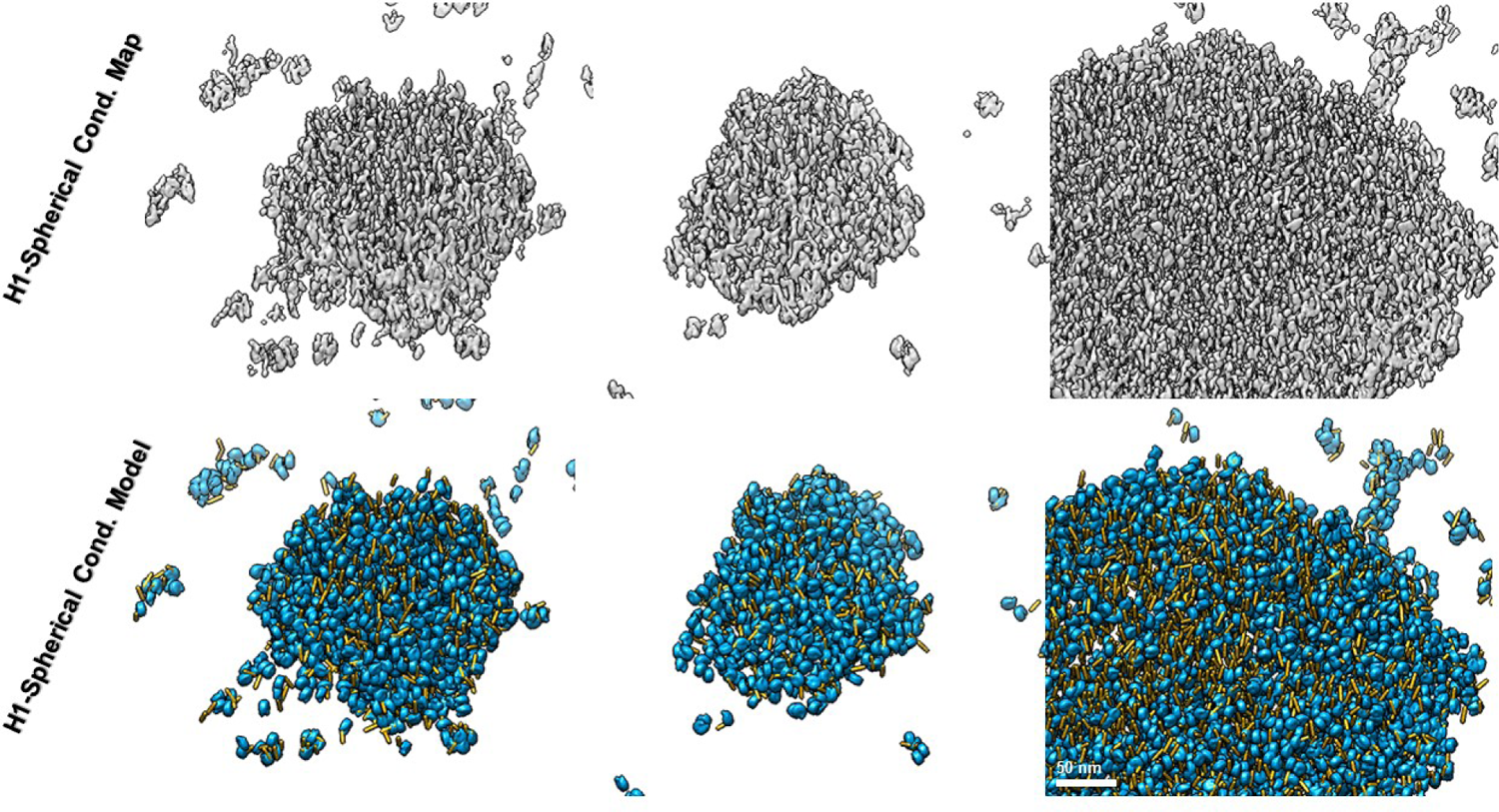
Fitted model of spherical condensates with H1. Comparison of three representative final density maps (top panels) with the corresponding fitted models (bottom panels) of spherical condensates in the presence of H1. Cyan indicates NCPs and yellow indicates flanking ADN.

**Figure S15.**
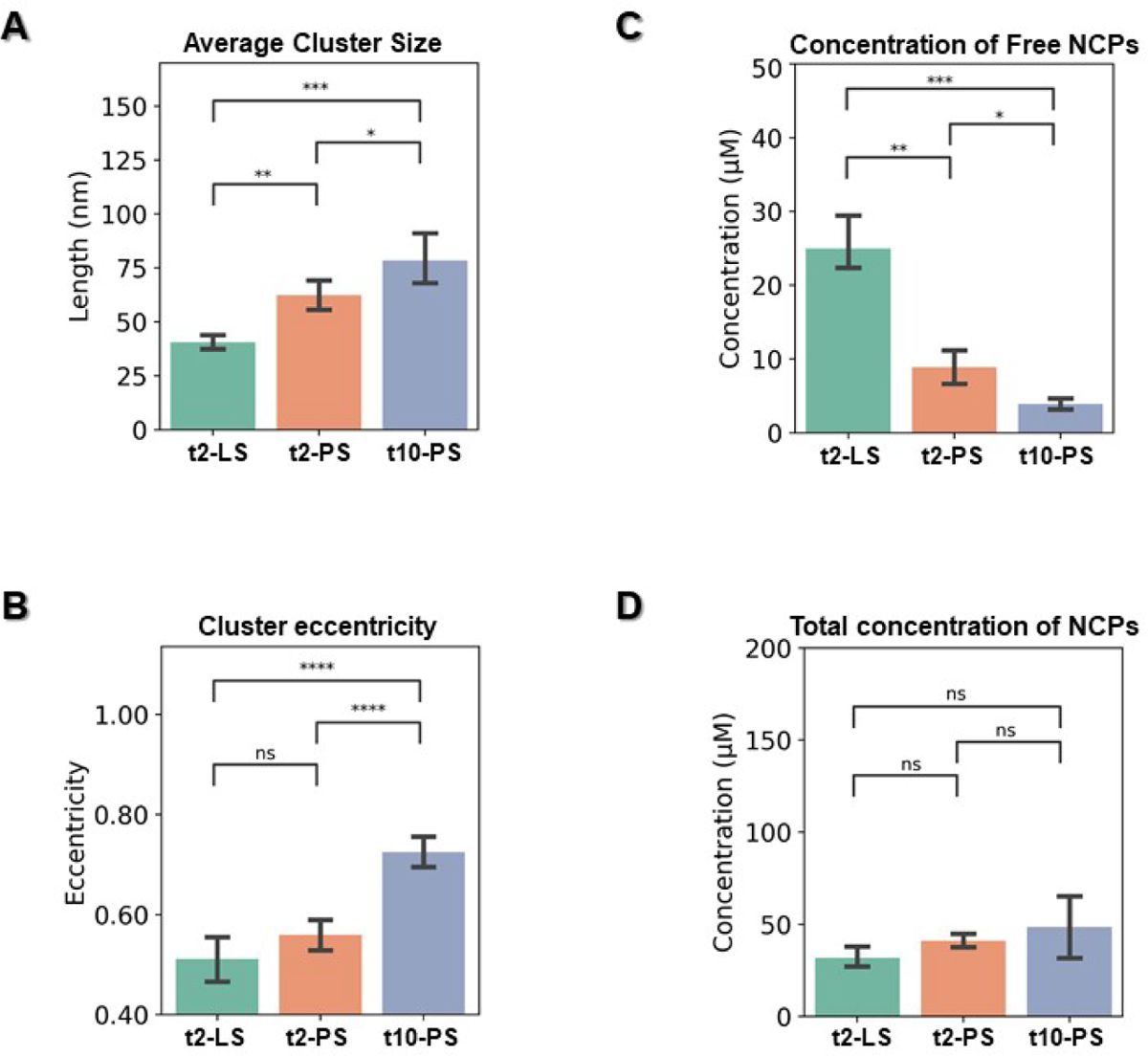
Statistical analysis of average cluster size, eccentricity, and NCP concentration for irregular condensates. (A) Average dimensions of irregular condensates expressed as a mean between the long (PC1) and short (PC3) axes obtained from a Principal Component Analysis (PCA). (B) Quantitation of irregular condensates eccentricity, expressed as (1 – PC3/PC1). (C) Concentration of free NCPs within the ice slab (number of free NCPs divided by the ice volume of the reconstructed tomogram). (D) Total NCP concentration within the ice slab (number of all identified NCPs divided by the ice volume). All data are plotted as mean ± SEM and measurements are compared using a two-tailed unpaired t-test, where ∗p < 0.05, ∗∗p < 0.01, ∗∗∗p < 0.001, ∗∗∗∗p < 0.0001; ns, not significant.

**Figure S16.**
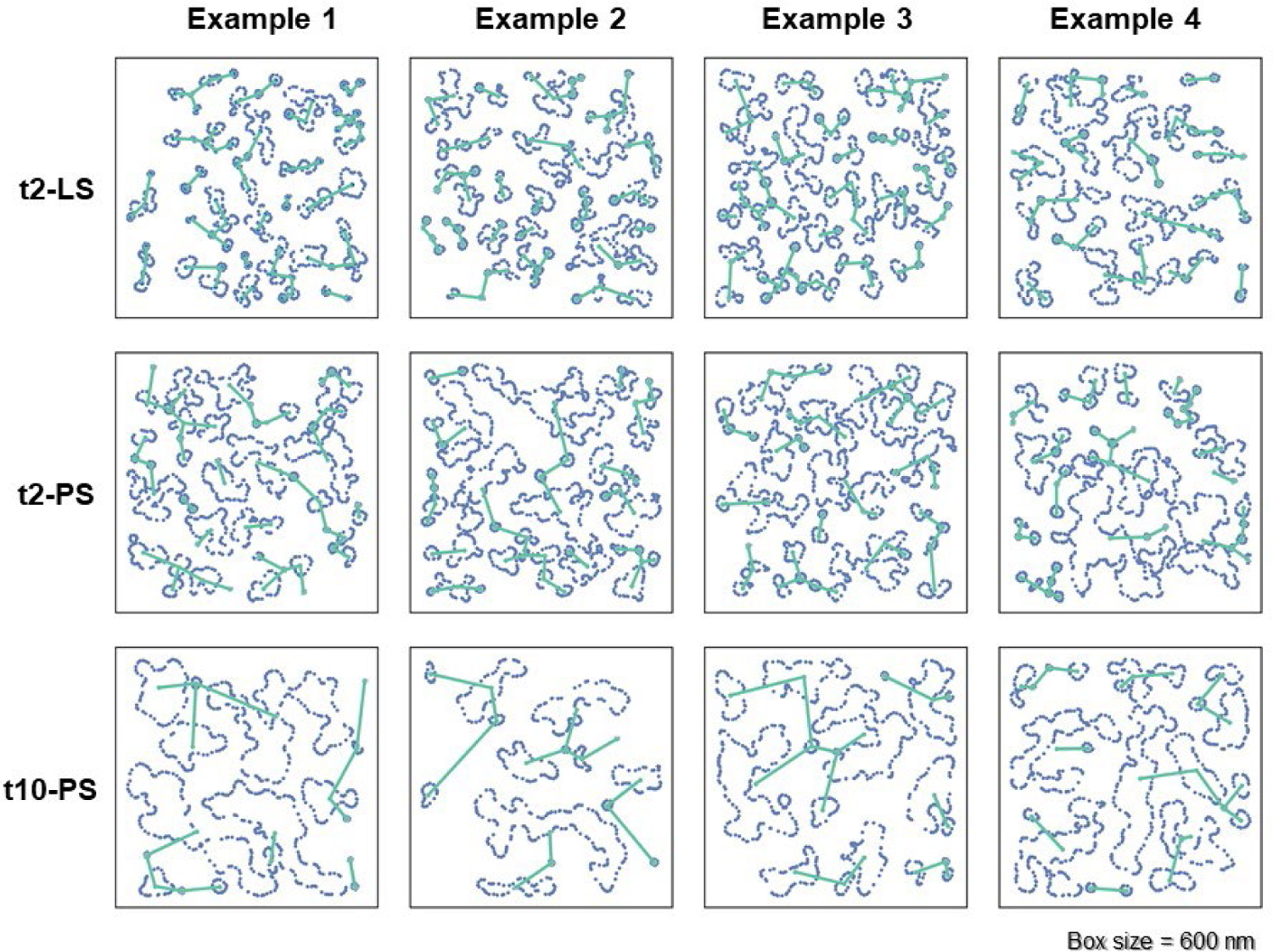
Measurement of the nearest neighbor distance among irregular condensate clusters. Identification of condensate-shaped contours (blue dots) within the z-dimension central slice for early (t2-LS), intermediate (t2-PS), and late irregular (t10-PS) condensates (top to bottom). Four examples of these three conditions are presented (columns). Nearest-neighbor condensates are connected by green lines, whose mean distances were used to represent the change of the critical wavelength, describing the evolution of spinodal decomposition in time.

**Figure S17.**
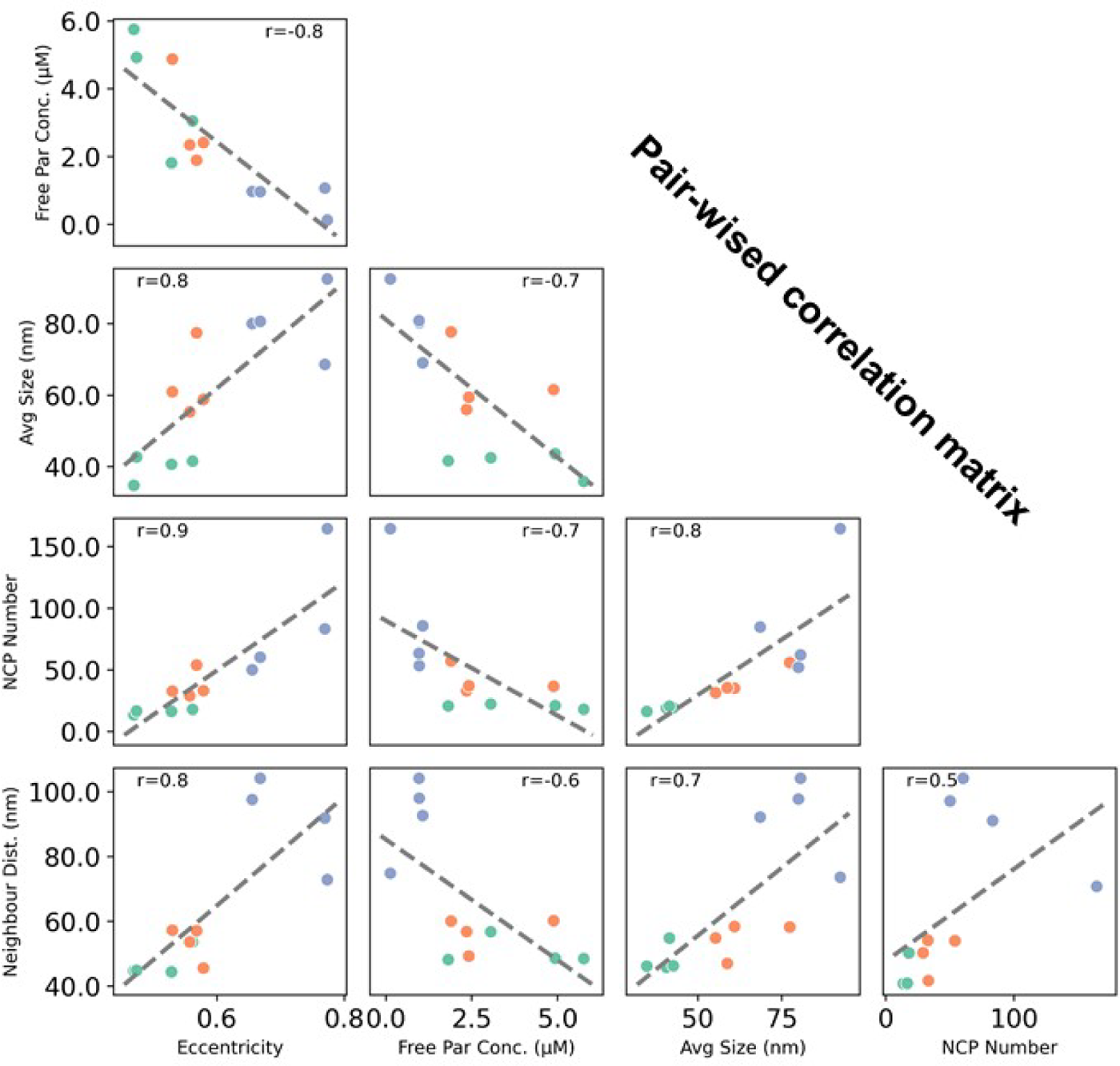
Pair-wise correlation matrix between independently determined number of NCPs per condensate, condensate size, eccentricity, neighbor distance among condensates, and concentration of free NCPs surrounding condensates. The mean value of the above five independent measurements from tomograms obtained for early, intermediate, and late stage irregular condensates are presented in green, orange, and blue points, respectively. The positive correlation observed between all parameters except with the concentration of free NCPs surrounding condensates, is consistent with a spinodal decomposition process.

**Figure S18.**
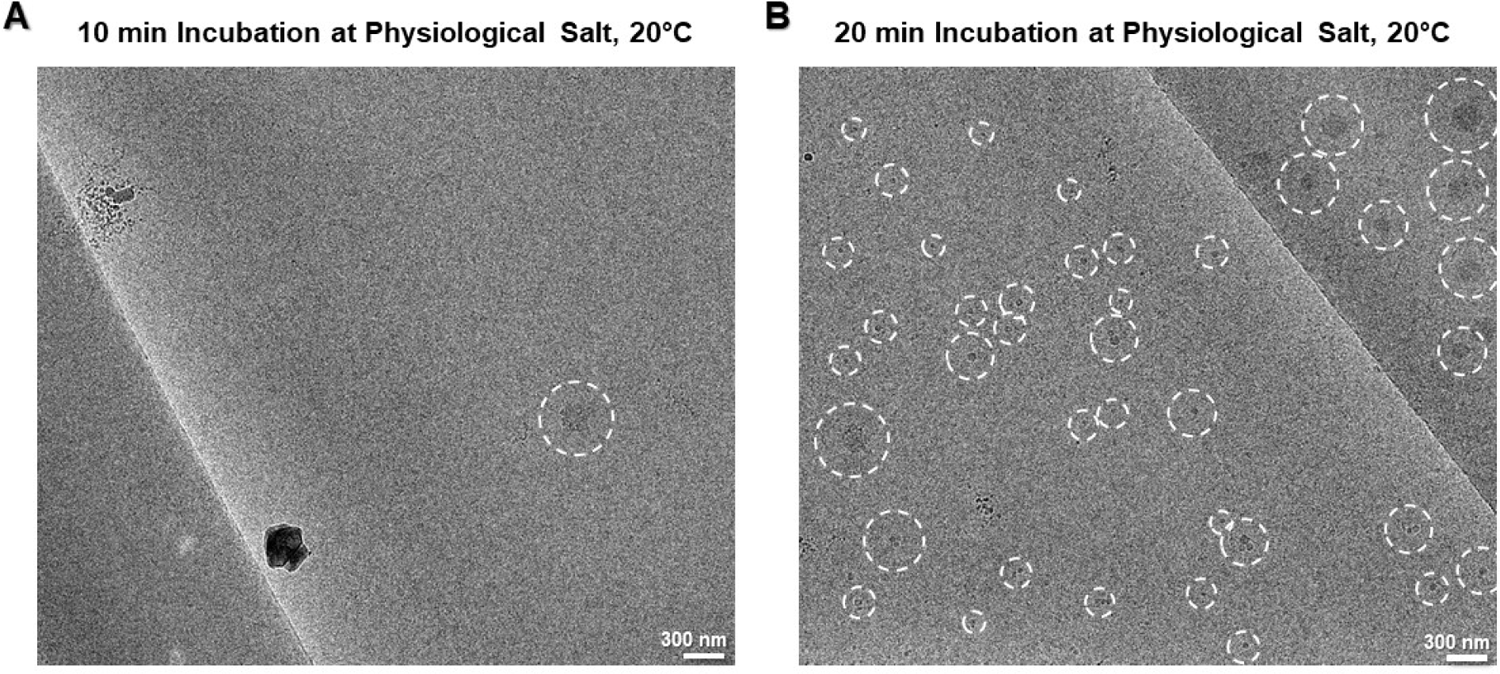
Spherical condensate visualized by cryo-EM. (A) Representative low magnification cryo-EM images showing the sparsely distributed spherical condensates on the SA-crystal surface after 10 min incubation in physiological salt at 20°C. (B) Representative low magnification image showing the appearance of small spherical condensates of varying sizes under the same conditions after 20 min incubation.

**Figure S19.**
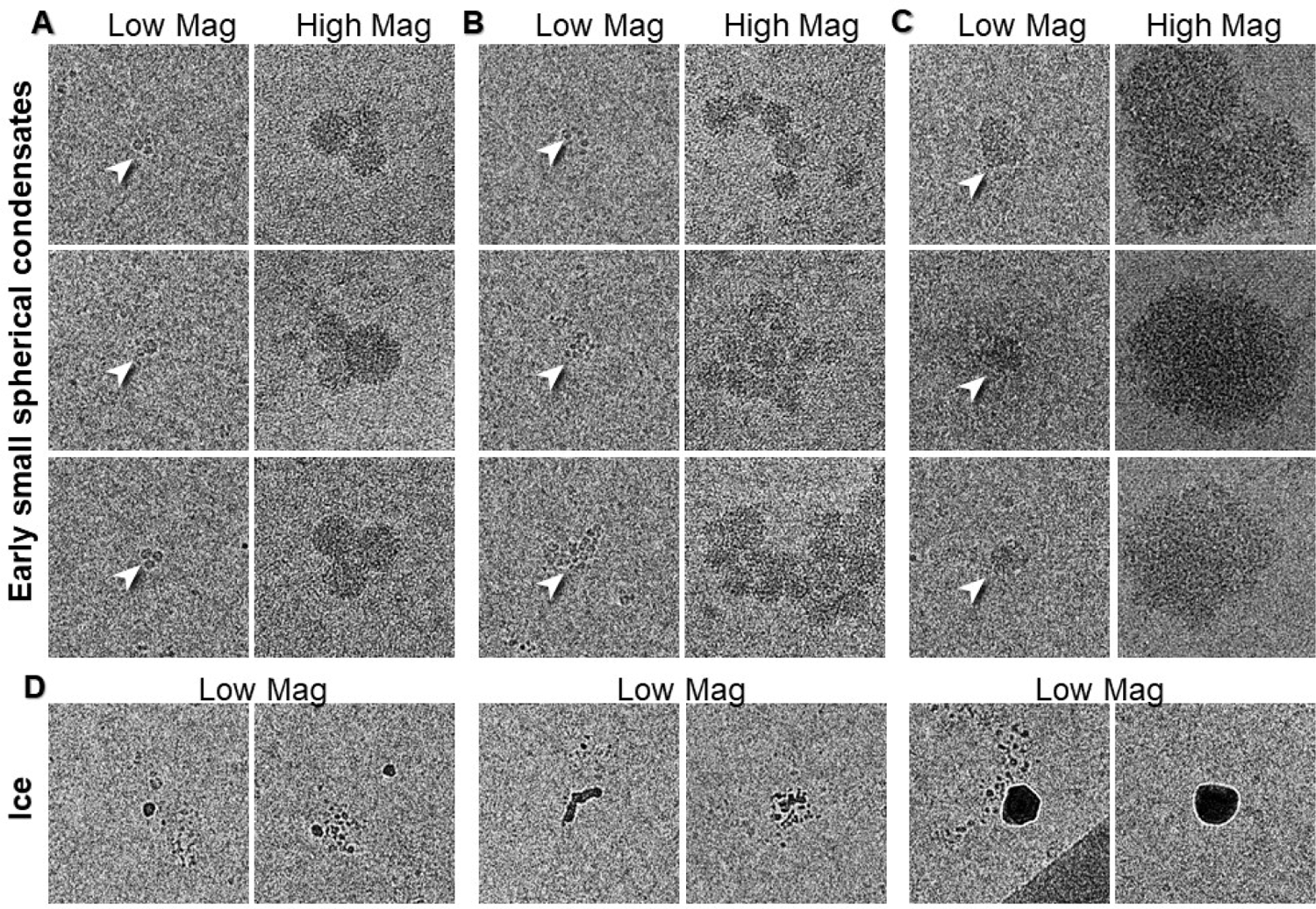
Distinction between the early small spherical condensates and ice contaminates from cryo-EM images. Representative cryo-EM images of a small (A) and a large bundle of attached small spherical condensates (B), and larger spherical condensates (C). These spherical condensates were imaged at both low (3,600 X; left column) and high (53,000 X; right column) magnification, where the latter was used to confirm the detailed nucleosomal pattern within the spherical condensates. (D) Ice contaminates observed in the low magnification images displayed a distinct high contrast and white fringe when compared to nucleosome condensates at similar sizes. These ice particles are excluded from the following statistical analysis of small condensate size distribution. Box size for Low Mag= 1,000 nm; Box size for High Mag= 220 nm.

**Figure S20.**
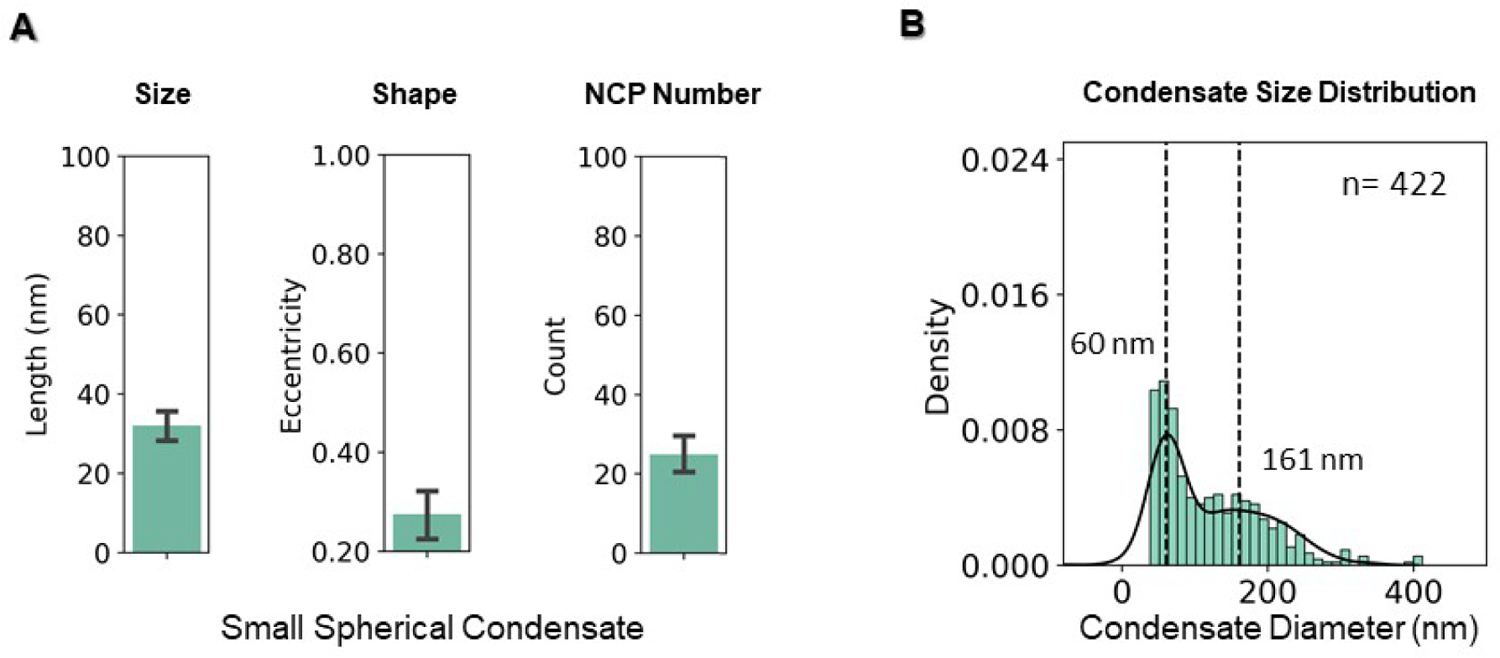
Quantitative analysis of small spherical condensate geometry and size distribution after 20 min incubation in physiological salt at 20°C. (A) Statistical analysis of the size (defined as the mean of the PC1 and PC3 axes after PCA analysis), eccentricity (1 – PC3/PC1 axis), and number of containing NCPs of small spherical condensates, measured from the reconstructed tomogram 3D models (n*=* 27). (B) Histogram of spherical condensate size distribution measured from low magnification 2D cryo-EM images (Fig. S18B; n*=* 440). Data was fitted using two Gaussians distributions. Dashed lines correspond to the mean of the two distributions (60 and 161 nm).

**Figure S21.**
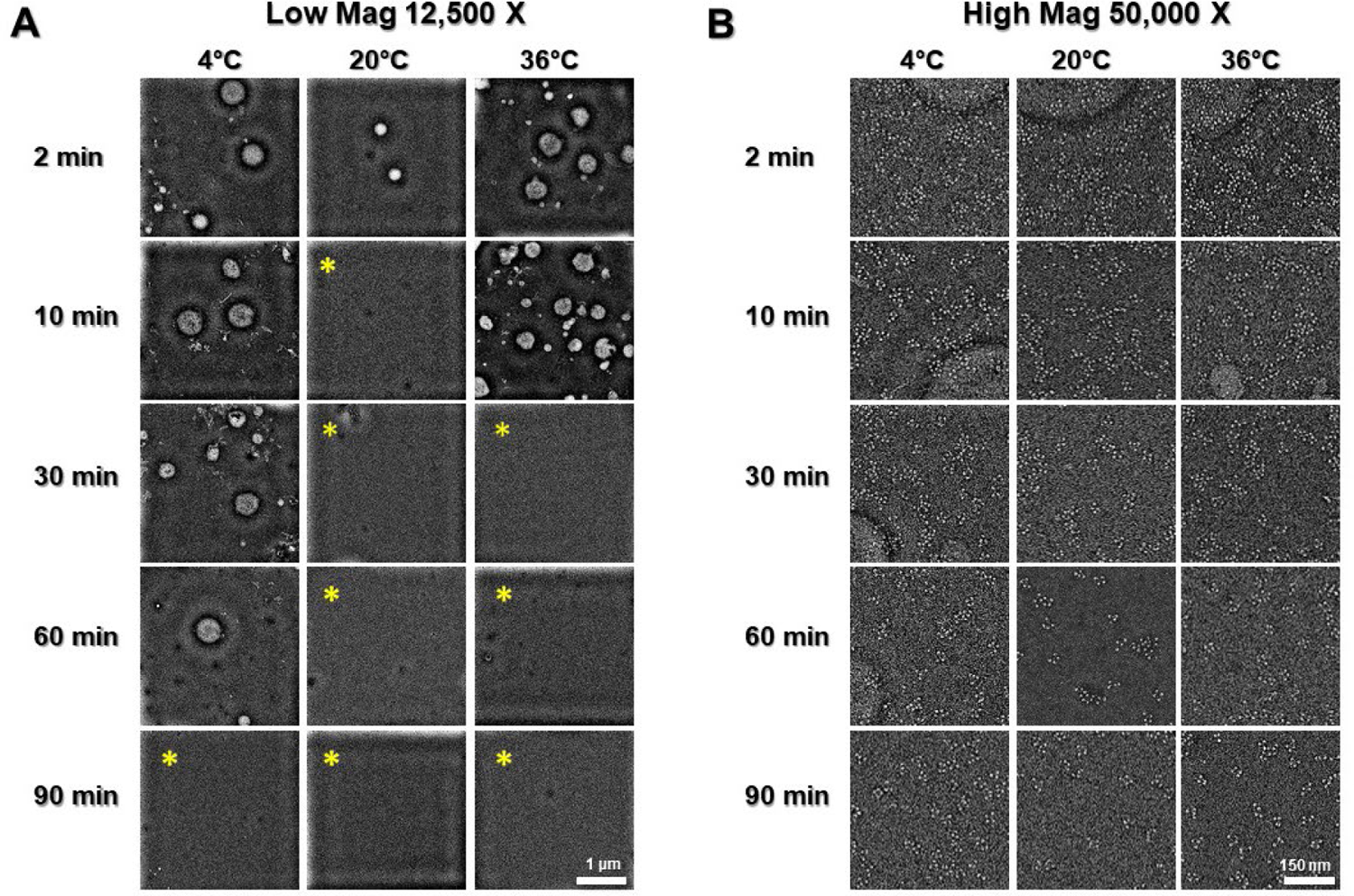
Tetranucleosome phase transition in the presence of H1 visualized by OpNS. Representative time series images of 30 nM tetranucleosome in physiological salt at 4°C, 20°C, and 36°C, in the presence of 120 nM of H1. Images were taken at 12.5k (A) and 50k (B). The low chance of finding spherical condensates on the grid surface after longer incubation times (yellow asterisk) should be an inevitable outcome in the presence of H1. This is because H1 can dramatically speed up the merging of small spherical condensates, which yield larger but fewer droplets given the limited amount of nucleosomal material in solution. This inference is supported by 1) the spherical condensate grid surface number density decreased gradually; 2) a significant amount of spinodal material was depleted at 90 min of reaction; 3) the disappearance of spherical condensate at earlier time points is consistent with a faster material depletion rate (20°C > 36°C > 4°C; Fig. S22); 4) no spherical condensates over >1.8 µm were observed in all conditions maybe because they were too large to be retained on grid surface.

**Figure S22.**
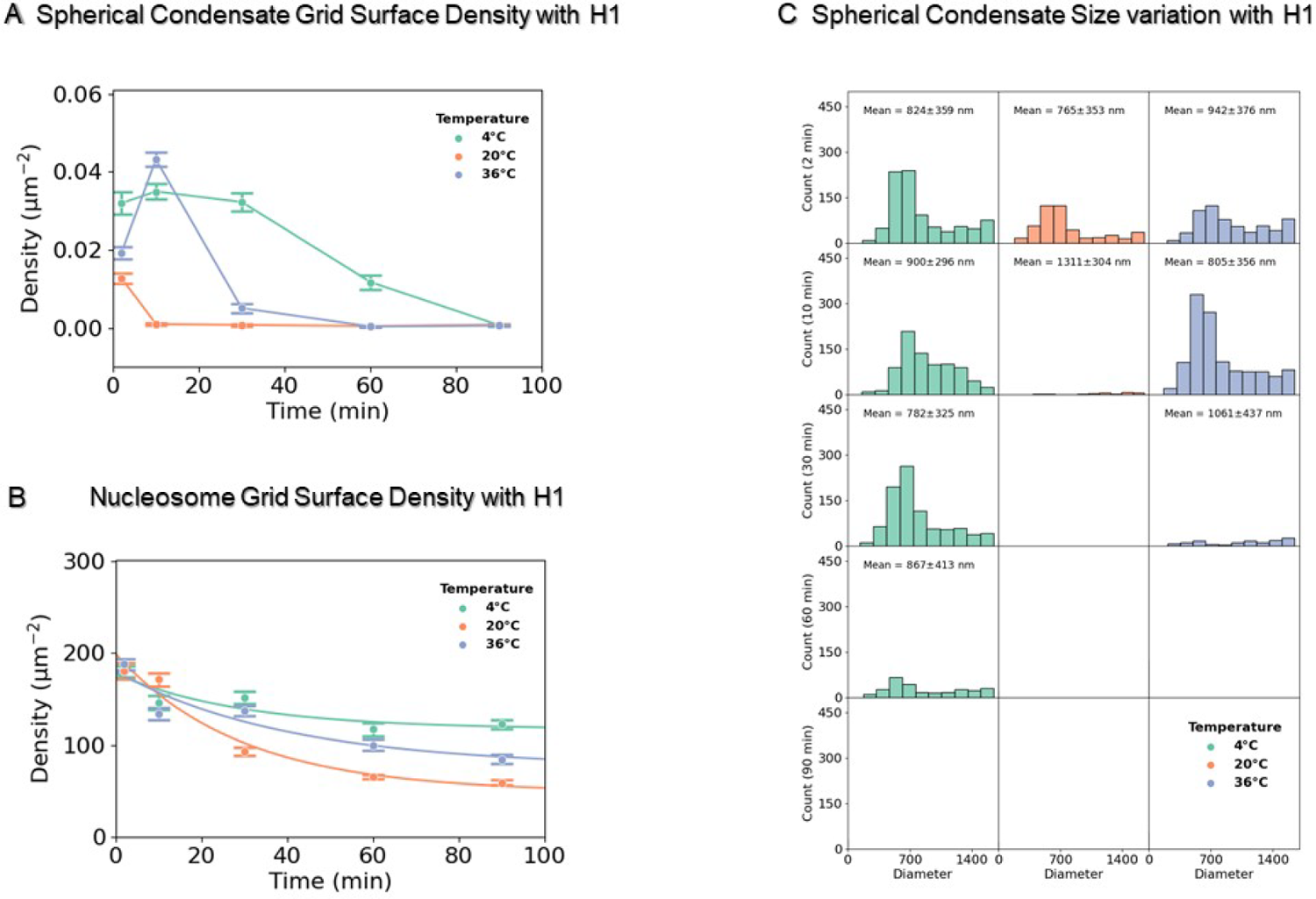
Quantitative measurement of tetranucleosomes and spherical condensates grid surface density and spherical condensates size in the presence of H1. (A) Statistics of spherical condensate density on the carbon grid surface in the presence of H1 (number of condensates divide by the imaging area) measured from incubation time series of OpNS samples (Fig. S21A). The incubation time series were conducted at 4°C (green), 20°C (orange), and 36°C (blue). (B) Statistics of nucleosome density in the presence of H1 (number of nucleosomes within spinodal condensates over the corresponding imaging area) measured from the same time series samples at higher magnification (Fig. S21B). For both (A) and (B), between 10-12 images were collected to generate the statistics which are presented as mean ± SEM. The data in (B) was fitted with an exponential decay function. (C) Size (diameter) distribution of spherical condensate in the presence of H1 (measured from images of (A)).

**Figure S23.**
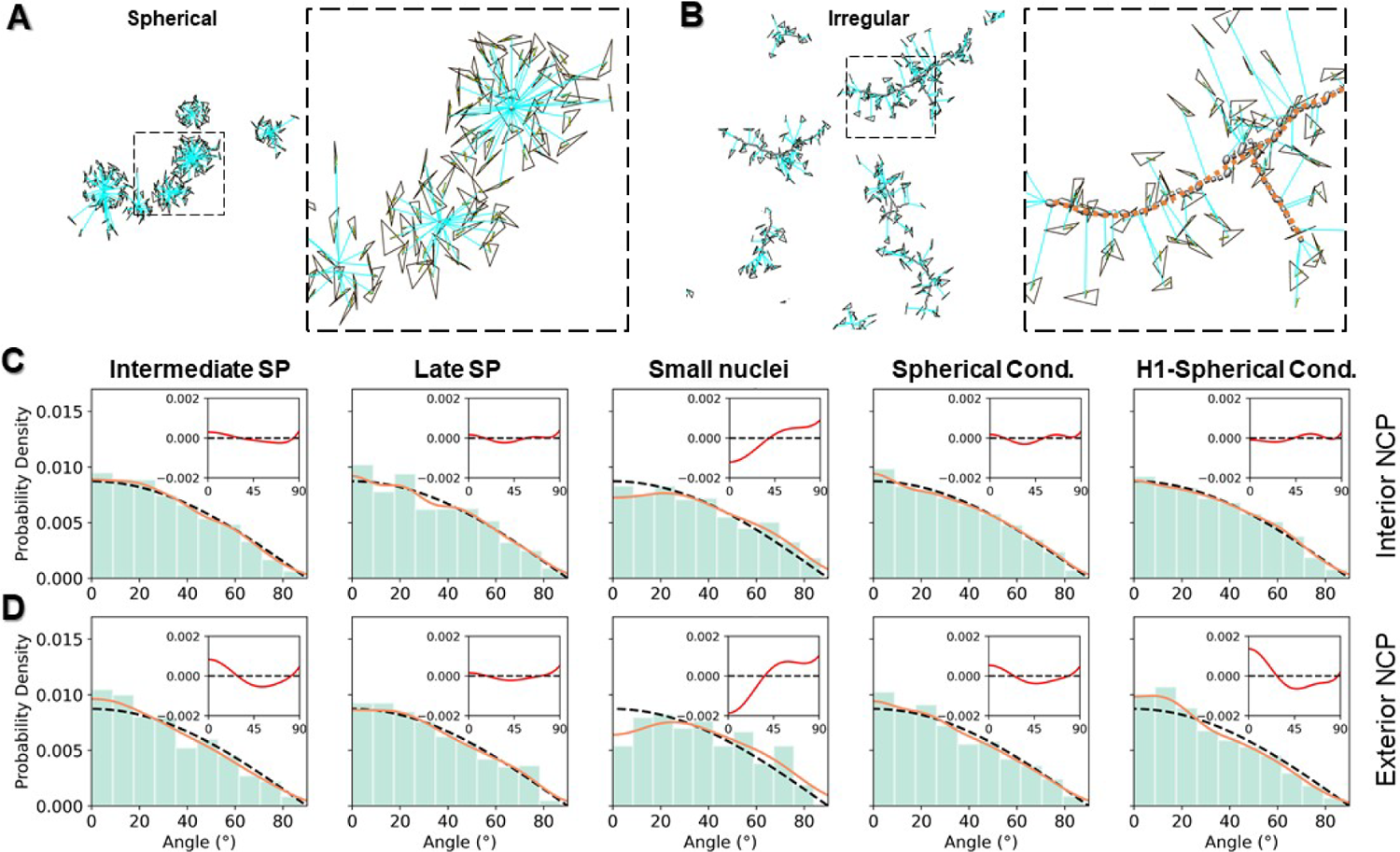
Quantitative analysis of NCP orientation on the surface and in the interior of condensates. (A) Angle measurement of NCP disk plane (black triangle) relative to the radius of the spherical condensate (cyan line) that crosses the center of the triangle (left panel). The zoom-in view of the dashed-line box region is shown on the right panel. (B) Angle measurement of nucleosomal disk plane (back triangle) relative to the radial line (cyan line, which is perpendicular to the axial axis of the irregular condensate) that crosses the center of the triangle (left panel). The axial axis of the irregular condensate was determined as the skeleton of the density map (orange dashed-line, on the right panel). (C and D) Histogram of nucleosomal disk angle distribution on the (C) surface of the condensate and (D) in the interior of the condensates. The angle distribution was fitted with an 8^th^ power polynomial (orange line) and compared with a *cos(*θ) function (black dash line, the angle distribution of a random rotated plane against a fixed axis following a cosine distribution). The deviation of experimental measurements from the *cos* was plotted on the top insets (red curves) indicating the preferred nucleosomal disk orientation found within each type of condensate.

**Figure S24.**
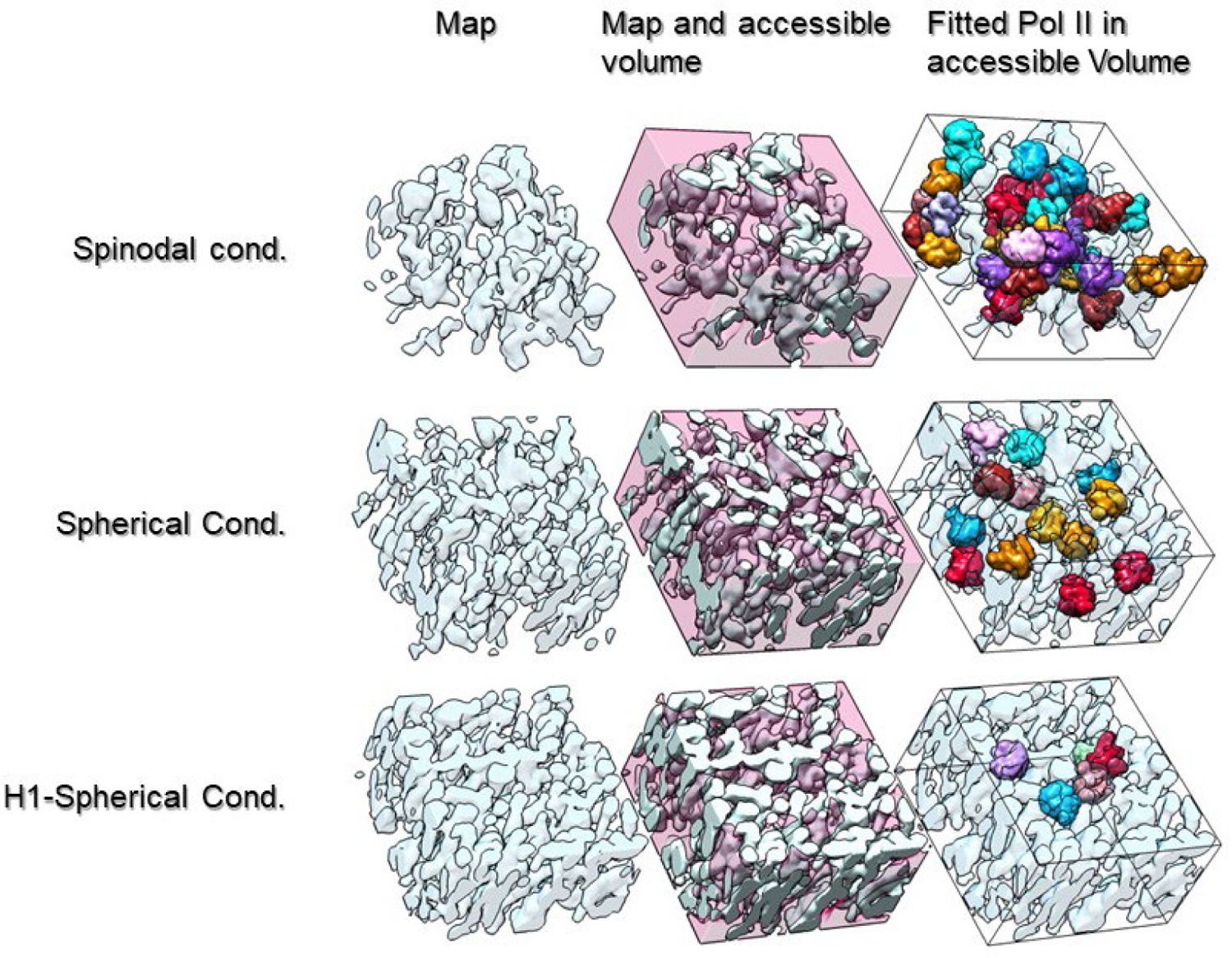
Evaluation of chambers capacity to RNA Polymerase II (Pol II) for different types of condensates. Sampling volumes of 2.88 x 105 nm^3^ from the final density maps (first column) of late stage spinodal condensates, spherical condensates, and spherical condensates in the presence of H1 (from top to bottom rows, respectively) were inverted and 3 nm low-pass filtered to show the detected empty chambers (second column). The capacity of the chambers was evaluated by quantitating the number Pol II molecules that fit within each condensate (third column, each Pol II was labeled with a random color). The number of Pol II molecules contained in the chambers from the late stage spinodal condensates, spherical condensates, and spherical condensates in the presence of H1, was 38,13, and 5, respectively.

**Figure S25.**
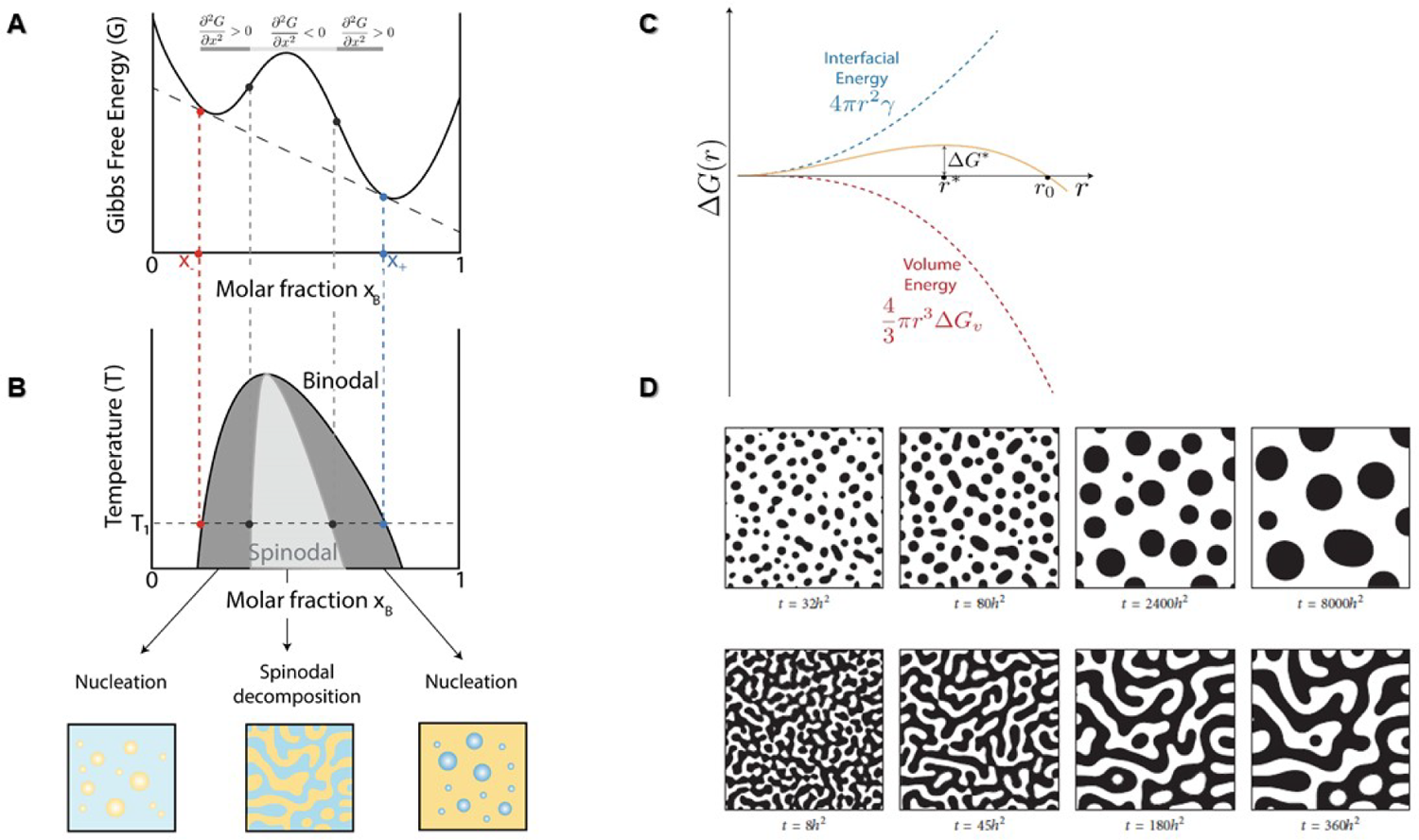
Schematics of phase transition mechanism. (A) Gibbs’ free energy as a function of molar fraction for a system that shows phase separation for a composition range X - < XB < X+. The straight dashed line is the only common tangent. (B) The light gray color area is the instability region in which the system undergoes demixing through spinodal decomposition. In the area in between the binodal and spinodal, the system demix via nucleation and growth (Alberti et al., 2019; Clerc and Cleary, 1995). (C) Gibbs free energy diagram for nucleation indicating the presence of a critical nucleus (*r\**) and a nucleation barrier (ΔG^*^) to overcome for growth to occur (Karthika et al., 2016). (D) Examples of spinodal decomposition predicted by the CH equation with different values of average composition and double-well potential function (ğ(ø) = 0.25(ø^2^ − 1)^2^), where ø is the composition of the system. The timescale *h* is a parameter used in the simulation and has the value of spatial step (similar to the area splitted in a grid). For this particular simulation, *h*= 0.03 (Image was taken from (Kim et al., 2016)).

---

**Movie S1. Simulation of two-step process of nucleosome phase separation.** Morphing trajectory of NCPs was linearly interpolated between two sets of 3D coordinates, which were obtained from two cryo-ET tomography docking models. To ensure that each NCP can find its counterpart from the first to the second tomograph, tomography containing similar number of NCPs (∼1100-1500) were selected as morphing pairs. These morphing pairs correspond to earlier and late stages of condensation. The movie morphs consecutively through six intermediate stages of phase separation, including the i) early, ii) intermediate, and iii) late stage of spinodal decomposition, iv) the formation of small nuclei clusters, their v) fusion into larger spherical condensates followed by the process of vi) accreting surrounding spinodal materials.

## Supplemental Theory Section

Phase separation is a non-equilibrium thermodynamic process in which a uniformly mixed system can lower its free energy (*G*) by segregating into two or more phases with distinct compositions that reach chemical equilibrium. The conditions at which distinct phases occur and coexist at equilibrium are described in a phase diagram. How do these conditions fulfill minimization of free energy (G) and satisfy the criteria for phase equilibrium? For a binary system exhibiting phase separation, the Gibbs free energy of a two-component solution as a function of composition exhibits a region of downward concavity or a negative curvature 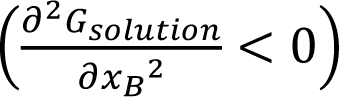 in which the mixed system is unstable (Fig. S25A; Supporting Supplementary Information). In this instability region, the single-phase system spontaneously undergoes demixing into two phases α and β, whose compositions must fulfill the criterion for phase equilibrium, i.e., that the chemical potential of each component of the binary mixture is the same in both phases (See equation 1, Supplemental Supporting Information). Since the chemical potential of component *i*, *µi*, is the first derivative of the free energy with respect to the concentration of component *i*, *ni*, the equilibrium condition implies that 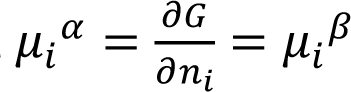, which can be obtained graphically at every temperature by finding the concentrations of both components for which the free energy has a common tangent (Fig. S25A) (Clerc and Cleary, 1995). The common tangent defines the points of the binodal curve, (Fig. S25B) which, in turn, defines the end point composition of phase separation, which may occur by different mechanisms.

## Spinodal Decomposition vs Nucleation and Growth

At compositions where 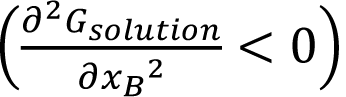, the mixed or single-phase system is unstable to both small and large fluctuations in composition. In this case, phase separation takes place without having to cross an energetic barrier and, therefore, it will occur throughout the entire system in a process known as spinodal decomposition (Fig. S25B). A detailed description of the equations involved in spinodal decomposition first formulated by Cahn and Hilliard (Cahn and Hilliard, 1958, 1959) is presented in the Supplemental Supporting Information section.

On the other hand, in the region in which the free energy as a function of composition has a positive curvature 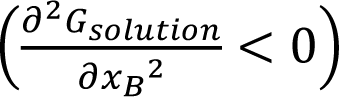 (Fig. S25A), the mixture is said to be metastable; in this region, phase separation only happens in discrete locations throughout the mixture where some rare, large fluctuations in composition spontaneously occur, because the system must overcome an energetic barrier. These localized regions, which appear sparsely in the mixture, are called nucleation sites (Clerc and Cleary, 1995; Schmelzer et al., 2004). Classical Nucleation Theory (CNT) describes phase separation in terms of a process involving the formation of a thermodynamically unfavorable phase separated ‘nucleus’ and an increasingly thermodynamically favorable process of ‘growth’. In CNT, the processes of nucleation and growth involve the interplay between the cost of an interfacial surface energy, or surface tension, and a favorable volume energy resulting from molecular interactions in the segregated phase that permits the nucleus to grow by the addition of more material to the new phase. (A general description of the parameters involved in the classical nucleation theory is presented in the supplementary information). However, in complex biological systems, such as the phase separation of lysozymes during crystallization, the CNT does not predict correctly the experimentally measured crystallization rates, which are ten orders of magnitude higher than the values predicted by CNT (Erdemir et al., 2009; Loh et al., 2017b; Vekilov, 2010). In these cases, non-classical nucleation models have been proposed, in which an additional step involving, for example, spinodal decomposition occurs before nucleation. For instance, a three-step mechanism has been proposed for the solidification and crystallization of gold nanocrystals based on liquid cell TEM studies (Loh et al., 2017a), which predicts that spinodal decomposition occurs before solidification and crystallization. Specifically, first, gold-rich spinodal structures form that later condense into amorphous nanoclusters, which crystallize into nuclei that can support nanoparticle growth because of their stable size (Ji et al., 2007; Loh et al., 2017b; Pong et al., 2007).

## Supplemental Supporting Information

In a multicomponent system of two or more phases, the phase equilibrium criterion is that the chemical potential of a given component *i* must be equal in all phases where *i* is present:

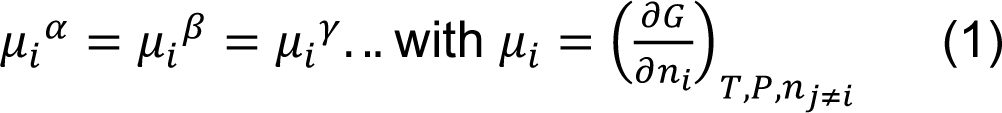

where μ_i_ represents the chemical potential of the *i*th component and α, β, γ, … represent the different phases where *i* is present at equilibrium (Clerc and Cleary, 1995). The compositions of those two phases α and β (expressed as the mole fraction of component B in these phases in figure S25B) are fixed at a given temperature but change as a function temperature. The collection of points defining the composition as a function of temperature at which these two phases coexist defines the binodal curve in the phase diagram (Fig. S25B) (Alberti et al., 2019; Clerc and Cleary, 1995; Shin and Brangwynne, 2017).

## Classical Nucleation Theory

Nucleation is the process by which a different thermodynamic phase with low free energy is formed from a parent phase with high free energy. The most common theoretical model to describe nucleation is the classical nucleation theory (CNT) that was formulated to explain the condensation of vapor into a liquid and that can be used also in liquid-solid equilibrium systems. In homogeneous nucleation, large supersaturations are usually needed to initiate the process because there are no preferential nucleation sites and the nuclei tend to form with equally low probability anywhere in the original mix (Karthika et al., 2016; Thanh et al., 2014).

For a spherical nucleus with radius *r*, the Gibbs free energy change ΔG(r) comprises the contributions of the volume free energy and the interfacial or surface energy cost of creating a nucleus inside the other phase (Equation 2; Fig. S25C),

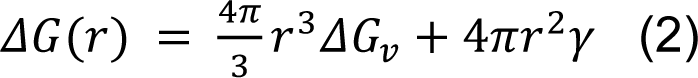

where γγ is the surface tension around the nucleus and ΔG_v_ is the free energy per unit volume gained from molecular interactions. The unfavorable interfacial energy dominates at small nucleus sizes *r*, while the favorable volumetric contribution to the nucleation barrier dominates at large nucleus sizes. The critical size or radius (*r*\*) for the nucleus corresponds to the minimum size beyond which the particle can be present in solution without being redissolved (Fig. S25C). From equation (2), 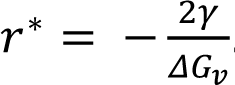. As shown in figure S25C, ΔG^∗^ indicates the nucleation barrier for the appearance of the separated phase. Clusters with values below *r*\* tend to redissolve and the ones with values above *r*\* tend to grow, thus, beyond *r*\*, ΔG(r) decreases with increasing *r* and at 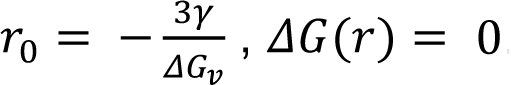. For r > r_0_, ΔG(r) is negative and particle growth leads to the formation of a new phase. Thus, phase separation from these nuclei can proceed through a process of growth that takes place when the surface energy price of creating a discrete phase in the medium is overcome by the energy gained inside the nucleus by the addition of more material to the phase. Several authors have estimated, using experimental data and simulations, that the size of the critical nucleus is in the range of ∼10-1,000 molecules. For instance, Yau and Vekilov reported that the size of the critical nucleus of protein apoferritin is about 40 nm in aqueous solution (Karthika et al., 2016; Yau and Vekilov, 2001).

## Spinodal Decomposition and the Cahn-Hilliard Equation

To evaluate the thermodynamics and kinetics underlying spinodal decomposition, it is necessary to use a theoretical framework that incorporates fluctuations in the composition field c(x, t). The Cahn-Hilliard (CH) model considers spatial inhomogeneities by adding a correction to the free energy function of a homogeneous medium. Based on the Ginzburg-Landau free energy theory (Cahn and Hilliard, 1958, 1959; Lee et al., 2014), the total free energy of a volume *V* of an isotropic system with a non-uniform composition is given by equation (3):

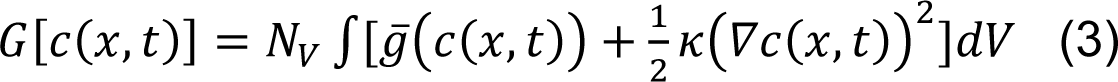

where N_v_ is the number of molecules per unit volume, ğ(c(x, t)) is the free energy per molecule of the homogenous system, ∇c(x, t) is the local composition gradient, and *K* is a parameter that controls the free energy cost of variations in concentration. Thus, the free energy of a volume of a non-uniform solution can be expressed as the sum of the free energy that this volume would have in a homogenous solution, and a gradient energy that is a function of the local composition. In this theoretical framework, the chemical potential is redefined as the functional derivative of *G* (equation (3)):

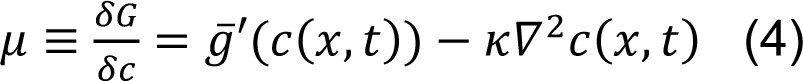

where ğ′ is the first derivative of the free energy per molecule with respect to concentration. During phase separation, fluctuations in the concentration of components will lead to the increase of concentrations of certain components at the expense of depletions on other regions. Therefore, a flux of components will be produced. The flux of components in mixture (*J*) can be described by an alternative formulation of Fick’s first law considering the variations in chemical potential (*μ*) (equation (5)):

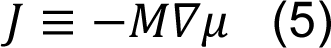

where *M* plays a role of the mobility (similar to the diffusion constant D used in Fick’s first law) defined as an interface parameter that indicates a measure of the transport kinetics across the interface of the phases (Kim et al., 2016). Using a continuity equation (conservation of mass):

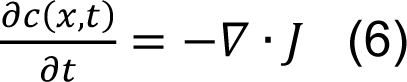

and by replacing equation (5) in the flux term (*J*), we obtain:

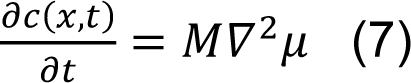

The replacement of equation (4) in equation (7) generates the Cahn-Hilliard equation, which describes the temporal evolution of the concentration field c(x, t) during spinodal decomposition:

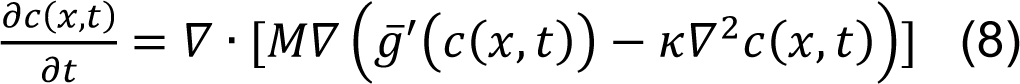

The CH equation is nonlinear and usually requires numerical approximations such as finite element, finite difference, or spectral methods to solve it. Spectral methods are common methods involving the use of fast Fourier transforms to solve differential equations. Computational simulations on the evolution of spinodal decomposition depicted by the CH equation are shown in figure S25D (Kim et al., 2016; Lee et al., 2014).

Some general implications can be obtained by evaluating particular conditions. For instance, for the double-well potential Gibbs free energy function (as the one depicted in figure. S25A), the concentration profile on the interface between the phases is diffuse and has a sigmoidal shape. In particular, for the one-dimensional case, it has the form:

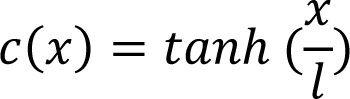

with 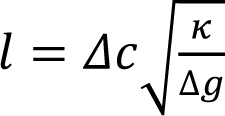 representing the interfacial width, where Δc = x_+_− x_−_, the width of the miscibility gap and where Δg ≥ kT. By evaluating how the CH equations responds to small fluctuations in concentration of form δc = Acos(kx − ωt) where k is given by 2*π*/λ and where λ is the wavelength of the concentration perturbation (as in a Fourier series), it can be shown that the critical wavelength has the following expression:

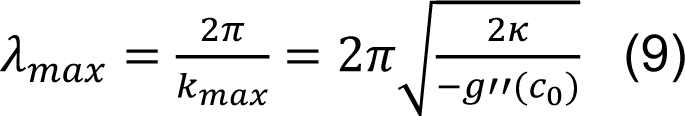

where *g*^”^ is the second derivative of the free energy with respect to concentration and the critical or maximum wavelength (λ_max_) is the most unstable wavelength in which random fluctuations decompose into characteristic patterns (Fig. S25D). There are two important conclusions from equation (9): i) λ_max_ only has a physical meaning when g˝(c_0_)<0 (a requirement for spinodal decomposition as discussed previously), and ii) λ_max_ gives a sense of the scale of the fluctuations and the size of domains formed during spinodal decomposition.

When one phase is at higher concentration, the Cahn–Hilliard equation predicts the process of Ostwald ripening, which is a thermodynamically-driven spontaneous process where the lower concentration phase forms spherical particles, and the smaller particles are absorbed through diffusion into the larger ones. The radii of these particles grow in time as in t ^1/3^(Lifshitz–Slyozov law) and the volume grows linearly with time. This process is favorable because the smaller particles are less energetically stable due to their high surface to volume ratio meaning high surface energy (Kim et al., 2016; Lee et al., 2014).

## Abbreviations

LLPS: Liquid-liquid phase separation

HP1: heterochromatin protein 1

cryo-ET: cryo-Electron Tomography

OpNS: optimized negative staining

NCPs: nucleosome core particles

IPET: individual-particle electron tomography

cc-score: cross-correlation score

DBSCAN: density-based clustering non-parametric algorithm

PCA: Principal Component Analysis

r: radial distribution functions g

AFM: Atomic Force Microscopy

Pol II: RNA Polymerase II

## Notes

### Competing Interest Statement

The authors have declared no competing interest.

